# Small Extracellular Vesicle Signaling and Mitochondrial Transfer Reprograms T Helper Cell Function in Human Asthma

**DOI:** 10.1101/2024.04.30.589227

**Authors:** Kenneth P. Hough, Jennifer L. Trevor, Shaheer Ahmad, Yong Wang, Balu K. Chacko, Kayla Goliwas, John G. Strenkowski, Nathaniel B. Bone, Young-il Kim, Renita Holmes, Yuelong Liu, Shia Vang, Alexandra Pritchard, Jay Chin, Sandeep Bodduluri, Veena B. Antony, Sultan Tousif, Mohammad Athar, Diptiman Chanda, Kasturi Mitra, Jaroslaw Zmijewski, Jianhua Zhang, Steven R. Duncan, Victor J. Thannickal, Susanne Gabrielsson, Victor M. Darley-Usmar, Jessy S. Deshane

## Abstract

Small extracellular vesicles (sEVs) are known to orchestrate cell-cell communication, but the role of sEV signaling via mitochondria in perpetuating asthmatic airway inflammation is unknown. Myeloid-derived regulatory cells (MDRCs) are known to control CD4^+^ T cell responses in asthma. We demonstrate that airway MDRC-derived sEVs from asthmatics mediate T cell receptor engagement and transfer of mitochondria that induce antigen-specific activation and polarization of Th17 and Th2 cells; these cells are drivers of chronic airway inflammation in asthma. CD4^+^ T cells internalize sEVs containing mitochondria predominantly by membrane fusion, and blocking mitochondrial oxidant signaling in MDRC-derived sEVs mitigates T cell activation. Reactive oxygen species-mediated signaling that elicits T cell activation in asthmatics is sEV-dependent. Additionally, a Drp1-dependent mechanism in pro-inflammatory MDRCs promotes mitochondrial packaging within sEVs, which then co-localize with the polarized cytoskeleton and mitochondrial networks in recipient T cells. Importantly, intranasal transfer of mitochondria packaged sEVs enhances airway inflammation and Th polarization *in vivo* in a murine model of asthma. Thus, our studies indicate a previously unrecognized role for mitochondrial fission and sEV-mediated mitochondrial transfer-mediated signaling in dysregulated T cell activation and Th cell polarization in asthma which could constitute a novel therapeutic target.

## Introduction

CD4^+^ T cells orchestrate the immune response in diverse chronic inflammatory diseases, including asthma^1^. Pathogenic T helper type 2 (Th2) and T helper type 17 (Th17) subsets drive airway inflammation, remodeling and smooth muscle hyperplasia in asthmatics^2-11^. Recent advances implicate cellular crosstalk between the innate and adaptive immune systems in the initiation and propagation of the allergic immune response. In allergic asthma, antigen presenting cells (APCs) internalize antigens, present processed peptides bound to class II molecules^12-14^ and engage the T cell receptor (TCR), triggering a multi-molecular signaling cascade to activate CD4^+^ T cells^15^. HLA-DR expressing APCs, including immature myeloid cells, regulate T cell function through TCR engagement^16-21^. In human asthma, the pro-inflammatory HLA-DR^+^ subsets of myeloid-derived regulatory cells (MDRCs) promote proliferation of T cells via redox signaling pathways ^21^. Although antigen presentation by MDRCs is not well-defined, TCR engagement and redox mechanisms may synergize to regulate MDRC-mediated T cell activation.

Recently, acellular antigen presentation by exosomes, has been described as an unconventional mechanism of immune activation^22-25^. Small extracellular vesicles (sEVs) that are endosomally-derived class of extracellular vesicles, known for facilitating cellular crosstalk, are decorated with scaffolding proteins for immune receptors called tetraspanins^24, 26-31^. Tetraspanins cluster at the immune synapse, and aid in antigen presentation and immune signaling^30^ and, thus, sEVs may modulate the immune response via the immune synapse. While professional APC-derived MHC-peptide bound sEVs induce Th2 cytokines in experimental asthma^22, 24, 32^, antigen presentation by MDRC-derived sEVs as a driver of dysregulated CD4^+^ T cell responses has not been demonstrated.

We reported that MDRC-derived sEVs containing polarized mitochondria are internalized by CD4^+^ T cells^33^. Increasing evidence for mitochondrial components and respiration in sEVs suggests that functional mitochondria packaged within the sEVs may elicit diverse physiologic responses^33-39^. This unconventional signaling mechanism may compensate for the altered bioenergetics of recipient cells and/or provide an alternative mechanism to outsource autophagy/mitophagy during cellular stress^36, 37, 40, 41^.

Here, we show that MDRC-derived sEV mitochondria with intact membrane potential activate CD4^+^ T cell proliferation and polarization into Th2 and Th17 cells via mitochondrial oxidant signaling; this event is triggered by membrane fusion of sEV-derived mitochondria and downstream complex formation of polarized cytoskeleton and mitochondrial network in recipient T cells. We demonstrate that mitochondrial fission, regulated by Drp1 oligomerization, is required for packaging functional mitochondria within sEVs and implicates MDRC-derived sEV-dependent mitochondrial signaling in Th cell polarization and immune regulation of chronic airway inflammation in mice with asthma and in human asthmatics.

## Results

### Bronchoalveolar lavage fluid (BALF) sEVs from asthmatics drive CD4^+^ T cell proliferation and Th polarization

We determined the concentration and size distribution of human BALF EVs (Figure S1) from healthy controls and asthmatics (Table I: Characteristics of study subjects) BALF EVs were predominantly 65-150 nm with a median size consistent with sEVs (Figure S1A-D). Immune regulation by sEVs is attributed to expression of MHC-II (HLA-DR) and co-stimulatory molecules CD81, CD86 and CD54^25, 42^. We and others reported that the % HLA-DR^+^ EVs and HLA-DR expression is increased in airways of asthmatics compared to healthy controls^24, 33, 43, 44^. Additionally, we showed that sEVs derived from human BALF and airway MDRCs were MitoTracker Green^+^(MitoTG^+^) and transferred mitochondria to T cells^33^. Consistent with this, BALF sEVs that were CD63^+^ and CD63^+^MitoTG^+^ were HLA-DR^+^ and were increased in asthmatics; sEVs from both healthy controls and asthmatics expressed HLA-DR (Figure S1E-J). We utilized nanoimaging studies to further confirm the size (Figure S2A) and EV marker distribution of BALF sEVs (Figure S2B) and observed that a proportion of these sEVs were Tomm20^+^ (mitochondrial outermembrane protein) in both study groups (Figure S2B). High resolution imaging of BALF sEVs with Tomm20 antibody confirmed expression of this mitochondrial marker in both healthy and asthma groups (Figure S2C). Western blot analyses of BALF sEVs showed expression of sEV marker CD81 and TIM23 (an inner mitochondrial membrane protein) (Figure S2D).

**Table 1.** Characteristics of volunteer subjects enrolled in the study.

| **Characteristics of Enrolled Patients** |  |  |  |
| --- | --- | --- | --- |
| **Characteristics** | **Healthy**<br>**(N=22)** | **Asthmatics**<br>**(N=21)** | **p-value** |
| **Sex** |  |  |  |
| *Male* | 8 | 4 |  |
| *Female* | 14 | 17 |  |
| Median Age (IQR) | 44 (23) | 40 (26) | 0.3054 |
| Median IgE Titer (IQR) – IU/L | 24.71 (83.2) | 102.3 (556.2) | 0.0254 |
| Median Predicted % FEV1 (IQR) | 106.3 (20.14) | 85 (30) | 0.0001 |
| Median Eosinophils (IQR) | 123.6 (86.8) | 227 (146.2) | 0.0029 |
| Median Neutrophils (IQR) | 3.71 (1.61) | 3.59 (1.82) | 0.5371 |

BALF sEVs when co-cultured with peripheral autologous CD4^+^ T cells (10 sEVs:1 T cell ratio), induced their proliferation (CFSE dilution) (Figure 1A-1B, S3A). Furthermore, proliferation of autologous airway CD4^+^ T cells of asthmatics was increased in co-cultures with BALF sEVs (Figure 1C & Figure S3B). An increase in %CD69^+^ T cells (an early activation marker), and an increase in phosphorylated Zap70 (pZap70), a kinase downstream of the TCR (Figure 1D) was observed, suggesting an MHC Class II-TCR interaction.

**Figure 1.**
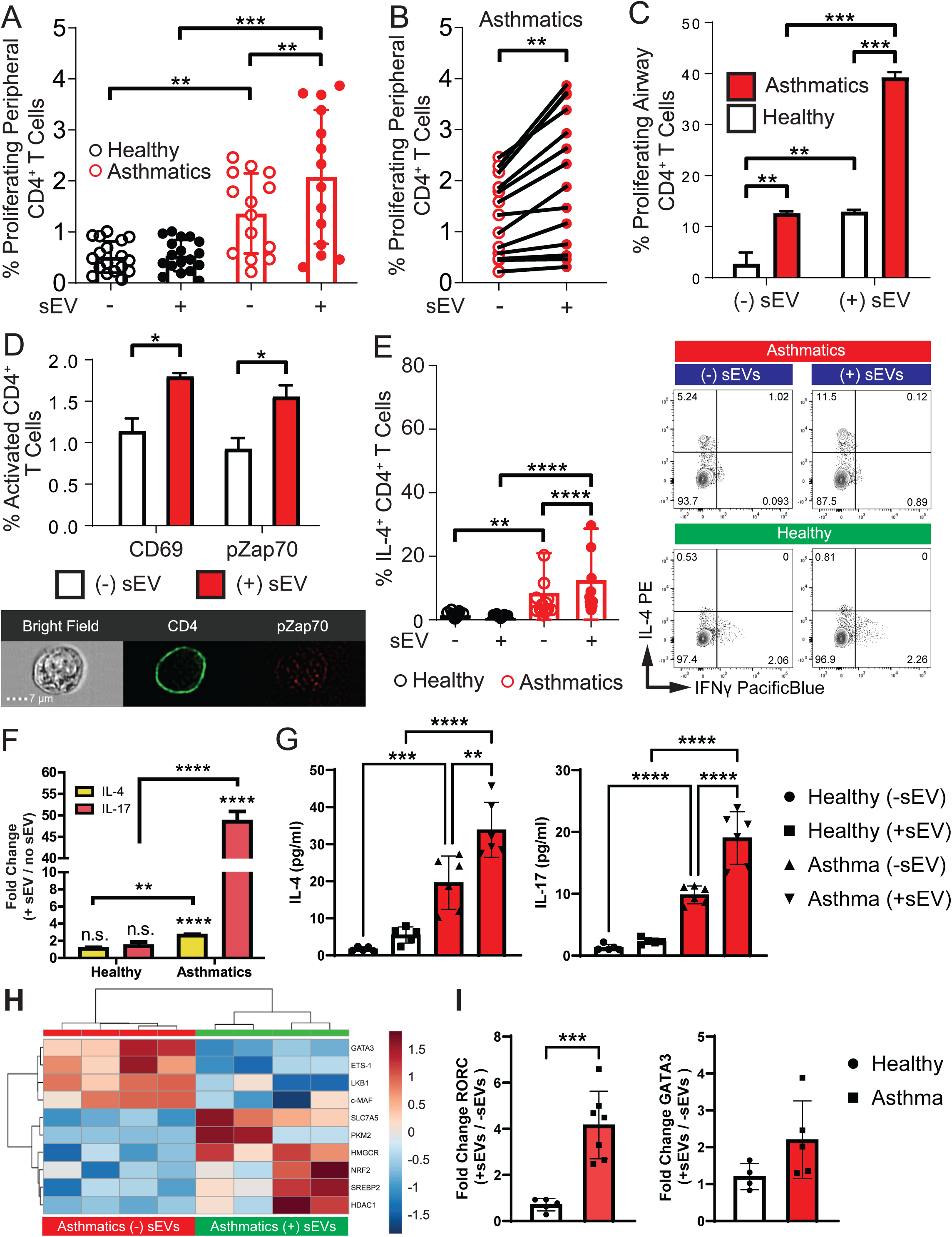
BALF sEVs from the airways of asthmatics promote proliferation and activation of CD4^+^ T cells, alters transcriptional signature and enhances T helper cell polarization. (A-B) Percentage of proliferating peripheral CD4^+^ T cells after co-culture with airway sEVs. Small extracellular vesicles (sEVs) from the BAL fluid (BALF) were co-cultured with CFSE labeled autologous peripheral CD4^+^ T cells at a ratio of 1:10 (T cells:sEVs) in the presence of rhIL-2 (50 IU/ml) for 7-days. Cells were harvested at day 7 for flow cytometry analysis on the BD LSRII. Mann Whitney T test for comparison between healthy and asthmatics, Wilcoxon matched-pairs signed rank test for intergroup (no sEVs vs sEVs), **<0.01, ***<0.001, ****<0.0001. (C) Percentage of proliferating airway CD4^+^ T cells after co-culture with airway sEVs. sEVs from the BALF were co-cultured as described above with autologous airway CD4^+^ T cells. Cells were harvested at day 7 for flow cytometry analysis on the BD LSRII. Mann Whitney T test for comparison between healthy and asthmatics, Wilcoxon matched-pairs signed rank test for intergroup (no sEVs vs sEVs), **<0.01, ***<0.001, ****<0.0001. (D) BALF sEVs were co-cultured with autologous CD4^+^ T cells (1:10 T cells:sEVs) for 15 minutes and percent CD69 and phospho-Zap70 was assessed by ImageStream imaging flow cytometry (representative image). Mann Whitney T test, *<0.05. (E) sEVs purified from the BALF were co-cultured with autologous peripheral CD4^+^ T cells in a ratio of 1:10 (T cells: sEVs) in the presence of rhIL-2 (50 IU/ml) for 7 days. Cells were harvested at day 7 for intracellular staining of cytokines and analyzed by flow cytometry using a BD LSRII. Left, percent of Th2 cells (assessed by IL-4 expression), and representative flow plots shown on the right. Mann Whitney T test for comparison between healthy and asthmatics, Wilcoxon matched-pairs signed rank test for intergroup (no sEVs vs sEVs), **<0.01, ****<0.0001. (F) Quantitative real-time PCR analysis of IL-4 and IL-17 gene expression of healthy and asthmatic CD4^+^ T cells co-cultured with and without sEVs for 7 days. Data are normalized to gene expression in T cells cultured without sEVs (ACTB as reference gene). Co-culture of asthmatic sEVs with T cells significantly increased IL-4 and IL-17 gene expression (**<0.01, ****<0.0001). (G) Cytokine measurements by ELISA of IL-4 and IL-17 in co-culture supernatants collected from CD4^+^ T cells co-cultured with autologous BALF sEVs at Day 7 of culture (n=6/study group/experimental group, 3 replicates each), One way ANOVA, **p=0.0016, ***p=0.0003 and ****p<0.0001 for IL-4, ****p<0.0001 for IL-17, (H) Clustered heatmap generated using MetaboAnalyst 3.0 from a custom NanoString panel comparing gene expression of CD4^+^ T cells from asthmatics co-cultured with BALF sEVs (right), and no sEVs control (left). Heatmap and clustering were generated using the nSolver Analysis Software 4.0 from nanoString, (I) Quantitative real-time PCR analysis of RORC and GATA-3 gene expression of healthy and asthmatic CD4^+^ T cells co-cultured with and without sEVs. Data are normalized to gene expression in T cells cultured without sEVs. Mann Whitney T test for comparison between healthy and asthmatic subjects ***p=0.004.

Additionally, airway sEVs from asthmatics modulated polarization of autologous CD4^+^ T cells into Th subsets. The increased baseline Th2 responses in asthmatics compared to healthy controls (Figure 1E) was further enhanced following co-culture with BALF sEVs (Figure 1E & S4). Th17 polarization was elevated in asthmatics with an increase in %IL-17^+^ T cells (Figure S3C & S4) and a robust increase in fold change of both IL-4 and IL-17 gene expression (Figure 1F) and cytokine production (Figure 1G). Modulation of transcription factors, metabolic genes, and epigenetic modifiers underlie Th polarization^45, 46^. Principal component analyses of gene expression in T cells from asthmatics co-cultured with or without sEVs indicated unique differences in gene programs (Figure S3D). Heatmaps with hierarchical clustering revealed differential modulation of metabolic genes, transcription factors, and cytokines (Figure S3E). The top 10 differentially expressed genes included sterol pathway genes (SREBP2, HMGCR), the glutamate transporter (*SLC7A5*), hexokinase 2 (*HK2*), Phosphoglycerate kinase 1 (*PGK1*) and histone deacetylase 1 (*HDAC1)*, in line with Th17 gene programming (Figure 1H and Figure S3E); *RORC*, the gene encoding the retinoic acid receptor-related orphan receptor gamma t (RORγt), a crucial transcription factor for Th17 cell development and function, was significantly increased in the asthma group (Figure 1I). The difference in GATA3 expression in these co-cultures from asthmatics did not reach significance (Figure 1I). Thus, airway sEVs from asthmatics enhance T cell proliferation, activation, and pathogenic Th polarization.

### MDRC-derived sEVs containing mitochondria are internalized by T cells via membrane fusion

We reported that the pro-inflammatory airway MDRC-derived sEVs containing mitochondria are internalized by peripheral autologous CD4^+^ T cells^33^. Although treatment of MDRCs with mitochondrial complex I inhibitor (rotenone) and uncoupler (FCCP) did not modulate the %MitoT Green^+^ sEVs, evaluation with a mitochondrial membrane potential dye, tetramethylrhodamine methyl ester (TMRE) showed that TMRE^+^MitoT Green^+^ BALF sEVs were present only in asthmatics (Figure S5A-S5C). Treatment of sEVs with FCCP or rotenone reduced the %TMRE^+^MitoT Green^+^CD63^+^ sEVs suggesting that the airways of asthmatics have an increased proportion of sEVs that maintain mitochondrial membrane potential (Figure S5B). Transduction of purified pro-inflammatory airway MDRCs with baculovirus constructs CellLight Mitochondria-GFP (Mito-GFP)^33^ yielded MDRC-derived GFP^+^ sEVs which were also internalized by T cells (Figure S5D).

Since, clathrin and dynamin-dependent endocytosis pathways or micropinocytosis, facilitate internalization of sEVs by immune and non-immune cells^47, 48^, we examined if their inhibition alters internalization of Mito-GFP^+^ sEVs by T cells. Although T cells are non-phagocytic, surface receptor recycling in CD4^+^ T cells is clathrin-dependent^49^. Treatment of CD4^+^ T cells with PitStop2 or Dynasore (inhibitors of clathrin and dynamin) decreased membrane- and cytoplasm-associated Mito-GFP^+^ sEVs only modestly (Figure 2A-2B), suggesting that endocytosis is not the primary mechanism for sEV internalization. To assess if sEV membrane fused with recipient T cell membranes, we pre-labeled Mito-GFP^+^ sEV with PKH26; in co-cultures, PKH26 signal remained restricted to the T cell membrane while the GFP signal was within the CD4^+^ T cells (Figure 2E). The percent of CD4^+^ T cells with membrane associated and internalized signals was not different between study groups (Figure 2C-2D). Thus, membrane fusion may be the primary mode of internalization of sEVs in lymphocytes.

**Figure 2.**
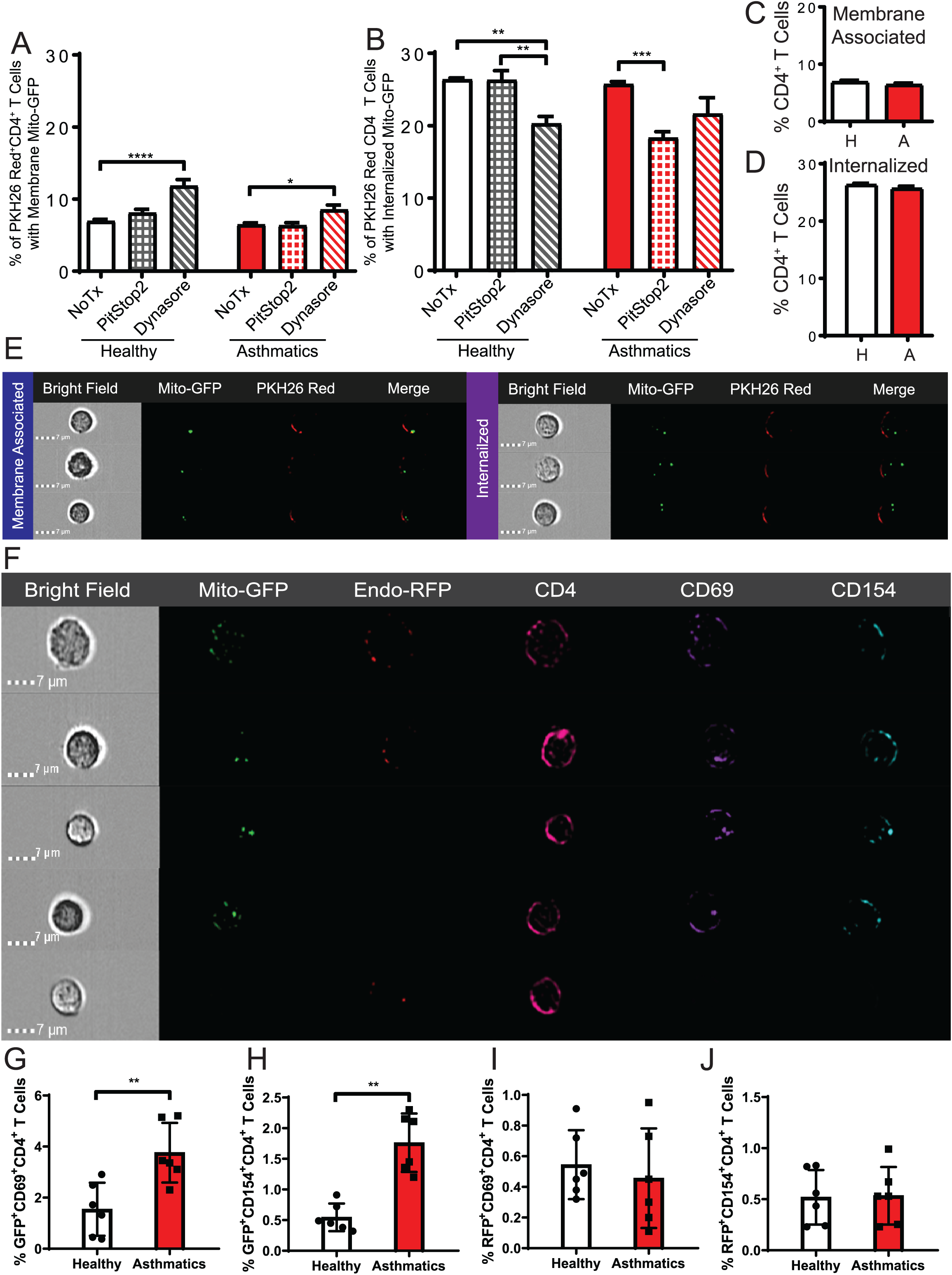
CD4^+^ T cells internalize MDRC-derived sEVs predominantly by membrane fusion and not by endocytosis. Activation of CD4^+^ T cells is associated with internalization of Mito-GFP^+^ sEVs derived from airway MDRCs. (A-B) Blockade of clathrin dependent and independent mechanism of endocytosis did not completely abrogate MDRC-derived Mito-GFP^+^ sEV internalization by CD4^+^ T cells. CD4^+^ T cells were treated with PitStop2 (50 nM), Dynasore (50 μM), or a combination of the two prior to co-culture with MDRC-derived sEVs for 24 hours and internalization assessed by ImageStream flow cytometry. (A) Quantitation of number of CD4^+^ T cells with membrane associated Mito-GFP^+^ MDRC sEVs. (B) Quantitation of number of CD4^+^ T cells with internalized Mito-GFP^+^ MDRC sEVs. (A-B) Mann Whitney T, *<0.05, **<0.01, ***<0.001, ****<0.0001. (C-E) MDRCs were transduced with Mito-GFP and membrane labeled with PKH26 red. sEVs derived from MDRCs were purified 48 hours later. Purified MDRC-derived sEVs were co-cultured with autologous peripheral CD4^+^ T cells for 24 hours in the presence of rhIL-2 (50 IU/ml) in a ration of 1:10 T cells:sEVs. Association of sEVs with the membrane or intracellular localization of sEVs were assessed by ImageStream flow cytometry. (C) Quantitation of membrane associated Mito-GFP^+^ MDRC sEVs. (D) Quantitation of intracellularly associated Mito-GFP^+^ MDRC sEVs. (E) Representative image strips from ImageStream flow cytometry analysis illustrating association of PKH26 red with the membrane and intracellular or membrane association of Mito-GFP. (F-J) MDRCs were transduced with CellLight Mito-GFP and CellLight Endo-RFP, and sEVs were purified from the supernatant 48 hours later. Autologous peripheral human T cells were cultured with MDRC sEVs for 24 hours, and internalization and activation assessed by ImageStream imaging flow cytometry. An sEV -generator-cell (MDRCs) to T cell ratio of 2.5×10^5^:1×10^6^ was maintained. (F) Representative images from ImageStream illustrating MDRC sEV internalization and T cell activation (CD69 and CD154). (G-H) Percent of GFP^+^CD69^+^CD4^+^ and GFP^+^CD154^+^CD4^+^ T cells that represent T cells that have internalized GFP^+^ sEVs and show early activation and antigen specific activation. (I-J) Percent of RFP^+^CD69^+^CD4^+^ and RFP^+^CD154^+^CD4^+^ T cells that represent T cells that have internalized RFP^+^ sEVs and show early activation and antigen specific activation (F-G) Mann Whitney T, **<0.01. (n=6/study group for (A-J) and 3 replicates each).

### Mito-GFP^+^ MDRC-derived sEVs are sufficient to trigger CD4^+^ T cell activation

We transduced MDRCs with baculovirus constructs Mito-GFP and CellLight Early Endosome-RFP (Endo-RFP) to confirm endosomal derivation of Mito-GFP^+^ sEVs. We observed two distinct populations of either Mito-GFP^+^ or Endo-RFP^+^ sEVs. The expression of CD81 and CD63 (Figure S5E) and %CD63^+^, CD81, or CD63^+^CD81^+^ sEVs were not different between Mito-GFP^+^ and Endo-RFP^+^ sEVs. Importantly, GFP and RFP signals did not overlap within sEVs, or co-localize following internalization in T cells (Figure 2F). Therefore, the Mito-GFP^+^ sEVs may have an alternate pathway of biogenesis than the Endo-RFP^+^ sEVs and/or they follow different paths following internalization by T cells. We investigated if internalized Mito-GFP^+^ sEVs elicit activation of co-cultured autologous peripheral CD4^+^ T cells, by assessing % CD69^+^ (early activation marker) and CD154^+^ (marker for antigen-specific activation) T cells. Only CD4^+^ T cells that internalized Mito-GFP^+^ or both Endo-RFP^+^ and Mito-GFP^+^ sEVs demonstrated a significant increase in both early and antigen-specific activation signals (Figure 2G-2H & 2F). Furthermore, activation was only seen in T cells of asthmatics and not in the healthy controls or in T cells that internalized only Endo-RFP^+^ sEVs (Figure 2I-2J). These data suggest that internalization of the Mito-GFP^+^ sEVs were sufficient for promoting T cell activation in asthmatics.

### Inhibition of MHC-II blocks MDRC-derived Mito-GFP^+^ sEV-mediated activation

We determined if MDRC-derived sEVs activate CD4^+^ T cells through MHC-II-TCR interaction (Figure 3A). When MDRC-sEVs were pre-treated with a pan-HLA antibody (targets HLA-DR, HLA-DP, and HLA-DQ), and then co-cultured with autologous peripheral CD4^+^ T cells, the activation of T cells that had internalized Mito-GFP^+^ (Figure 3B-3D) or both Mito-GFP^+^ and Endo-RFP^+^ sEVs (Figure S6A-S6D), was inhibited (Figure 3B-3D) in asthmatics. The internalization of sEVs was not abrogated (Figure 3D), suggesting that activation was independent of internalization. When MDRC-derived sEVs were pre-treated with LFA-1 antibody (blocks LFA-1-ICAM1 interaction), differences in early activation (CD69) were not observed (Figure 3C); however, antigen specific activation (CD154^+^) and CD69^+^CD154^+^ populations were reduced in asthmatics compared to sEV treatment alone.

**Figure 3.**
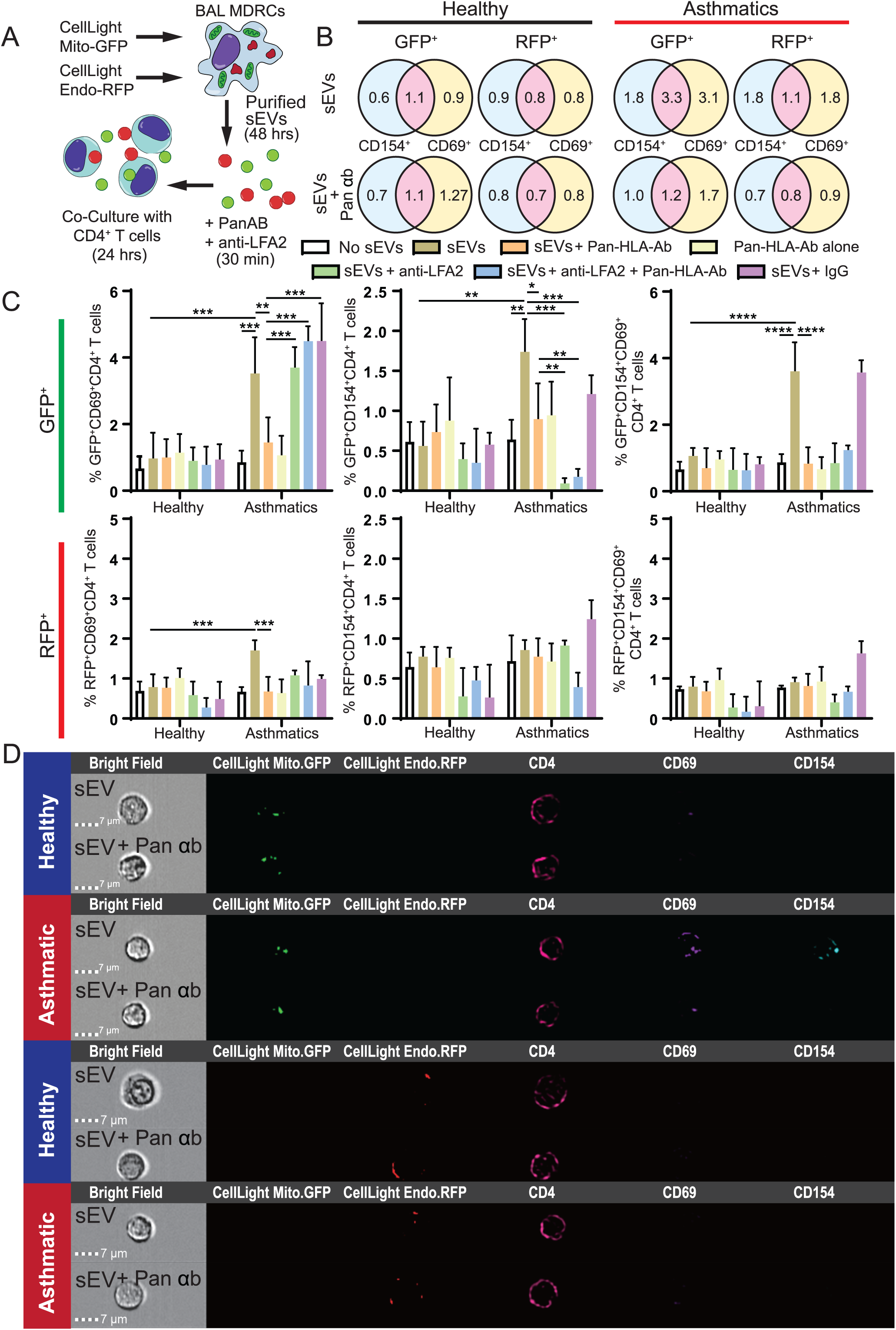
Blockade of class II molecules using a pan-HLA antibody (HLA-DR/DP/DQ) results in loss of MDRC sEV mediated activation of CD4^+^ T cells. (A) Illustration of experimental approach. (B-D) MDRCs were transduced with CellLight Mito-GFP and CellLight Endo-RFP, and sEVs were purified from the supernatant 48 hours later. sEVs derived from MDRCs transduced with Mito-GFP and Endo-RFP were co-cultured with autologous peripheral CD4^+^ T cells for 24 hours in the presence of rhIL-2 (50 IU/ml) in a ration of 1:10 T cells:sEVs. MDRC sEVs were pre-treated with pan-HLA-Ab (10 µg/ml), LFA2 antibody (1 µg/ml), IgG control, a combination of pan-HLA-Ab and anti-LFA2 for 30 minutes at room temperature. sEVs were washed and re-purified using the Invitrogen Total Exosome Isolation kit. ImageStream flow cytometry was used to assess early activation (CD69) and antigen-specific activation (CD154) marker expression. (B) Venn diagrams illustrating the percentages for CD154^+^, CD69^+^, or CD154^+^CD69^+^ T cells that internalized either Mito-GFP^+^ MDRC sEVs or Endo-RFP^+^ MDRC sEVs. The top row is untreated MDRC sEVs, and the bottom row is MDRC sEVs pre-treated with pan-HLA-Ab prior to co-culture. (C) Graphs illustrating the percentage of CD69^+^, CD154^+^, or CD69^+^CD154^+^ T cells that internalized either Mito-GFP^+^ MDRC sEVs (top row) or Endo-RFP^+^ MDRC sEVs (bottom row). Mann Whitney T, *<0.05, **<0.01, ***<0.001, ****<0.0001. (n=8-9/study group, 3 replicates each. (D) Representative image strips from ImageStream illustrating that Mito-GFP^+^ MDRC sEVs activate T cells in asthmatics, while treatment with pan-HLA-Ab abrogates this activation by MDRC sEVs.

### Inhibition of mitochondrial complex III and V abrogates MDRC-derived Mito-GFP^+^ sEV-mediated CD4^+^ T cell activation

We then determined if the mitochondria in Mito-GFP^+^ sEVs have functional effects on CD4^+^ T cell activation. Pre-treatment of MDRC-derived Mito-GFP^+^ sEVs with antimycin A, a high affinity complex III inhibitor (Figure 4A) before co-culture with T cells abrogated Mito-GFP^+^ sEV-mediated CD4^+^ T cell activation (Figures 4B-4D, S6C-S6D). To control for potential transfer of mitochondrial inhibitors from the sEV preparation to recipient cells, an sEV free isolation media with inhibitors was included which had no effect (Figures 4B-4D, S6C-S6D). To determine if the inhibition was dependent on ATP synthesis by sEV-mitochondria, we pre-treated MDRC-derived sEV with oligomycin, a mitochondrial ATP synthase inhibitor. Oligomycin pre-treatment abrogated Mito-GFP^+^ sEV-mediated activation of CD4^+^ T cells (Figure S7A-S7D), consistent with a requirement of a functional oxidative phosphorylation system within the sEV mitochondria to activate T cells.

**Figure 4.**
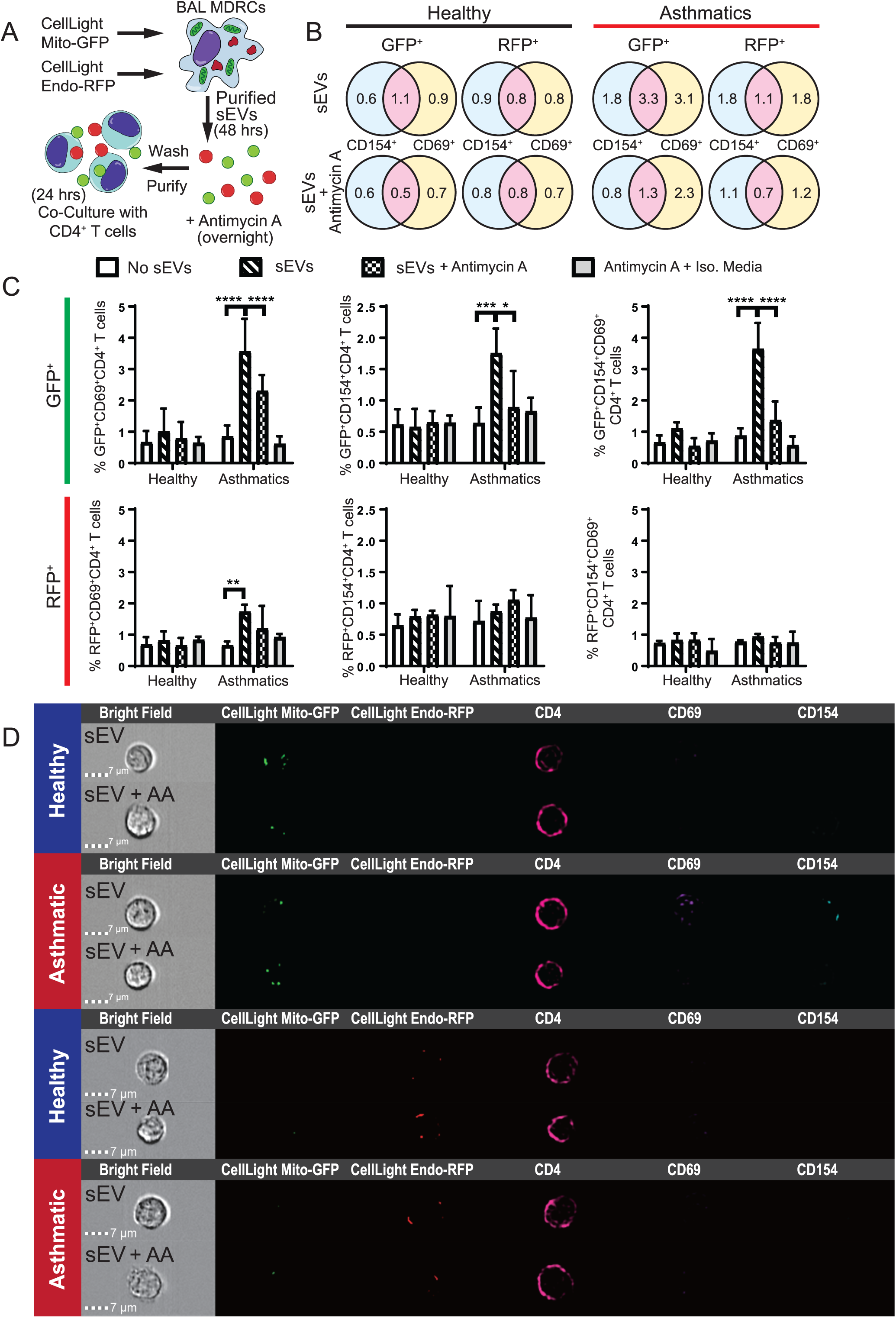
Inhibition of complex III by antimycin A in MDRC sEVs diminishes CD4^+^ T cell activation in asthmatics. (A) Illustration of experimental approach. (B-D) MDRCs were transduced with CellLight Mito-GFP and CellLight Endo-RFP, and sEVs were purified from the supernatant 48 hours later. sEVs derived from MDRCs transduced with Mito-GFP and Endo-RFP were pre-treated with antimycin A (10 µM) overnight or untreated prior to co-culture with autologous peripheral CD4^+^ T cells for 24 hours in the presence of rhIL-2 (50 IU/ml) in a ration of 1:10 T cells:sEVs. sEVs were washed and re-purified using the Invitrogen Total Exosome Isolation kit. ImageStream flow cytometry was used to assess early activation (CD69) and antigen-specific activation (CD154). (B) Venn diagrams illustrating the percentages for CD154^+^, CD69^+^, or CD154^+^CD69^+^ T cells that internalized either Mito-GFP^+^ MDRC sEVs or Endo-RFP^+^ MDRC sEVs. (C) Graphs illustrating the percentage of CD69^+^, CD154^+^, or CD69^+^CD154^+^ T cells that internalized either Mito-GFP^+^ MDRC sEVs (top row) or Endo-RFP^+^ MDRC sEVs (bottom row). Mann Whitney T, **<0.01, ***<0.001, ****<0.0001. (n=6-8/study group, 3-4 replicates each) (D) Representative image strips from ImageStream analysis illustrating that Mito-GFP^+^ MDRC sEVs activate T cells in asthmatics, and antimycin A blocks activation in asthmatics.

Interesingly, pre-treatment of MDRC-derived sEVs with the complex I inhibitor, rotenone, enhanced Mito-GFP^+^ sEV-mediated activation of peripheral CD4^+^ T cells in asthmatics and importantly, induced activation of healthy T cells (Figure S8A-S8D). This aberrant activation of CD4^+^ T cells, short circuiting the conventional MHC-II-TCR activation pathways, is unlikely to be solely due to effects on oxidative phosphorylation since oligomycin, antimycin A, and rotenone inhibit mitochondrial ATP synthesis but differ in their mechanisms. Oligomycin, rotenone and Antimycin generate ROS through different mechanisms and sites in the mitochondria^50, 51^.

### Activation of CD4^+^ T cells requires ROS generated within MDRC-derived sEVs

Next, we determined if rotenone affects ROS levels within sEVs and if the T cell activation was dependent on sEV ROS. We used MitoTEMPOL, a superoxide dismutase mimetic to scavenge ROS within the MDRC-derived sEVs following rotenone treatment (Figure 5A). To test if blocking complex II resulted in a similar effect to rotenone or antimycin A, we used thenoyltrifluoroacetone (TTFA), a complex II inhibitor. When MDRC-derived sEVs were treated with rotenone and MitoTEMPOL, the rotenone-induced activation of CD4^+^ T cells was inhibited (Figure 5A-5B) with a reduction in %GFP^+^CD69^+^CD4^+^, GFP^+^CD154^+^CD4^+^ and GFP^+^CD69^+^CD154^+^CD4^+^ T cells, specifically in CD4^+^ T cells that have internalized Mito-GFP^+^ or both Mito-GFP^+^ and Endo-RFP^+^ (Figure 5A-5B) but not just the Endo-RFP^+^ MDRC-derived sEVs (not shown). Furthermore, treatment of MDRC-derived sEVs with TTFA did not induce CD4^+^ T cell activation suggesting that complex II is not involved.

**Figure 5.**
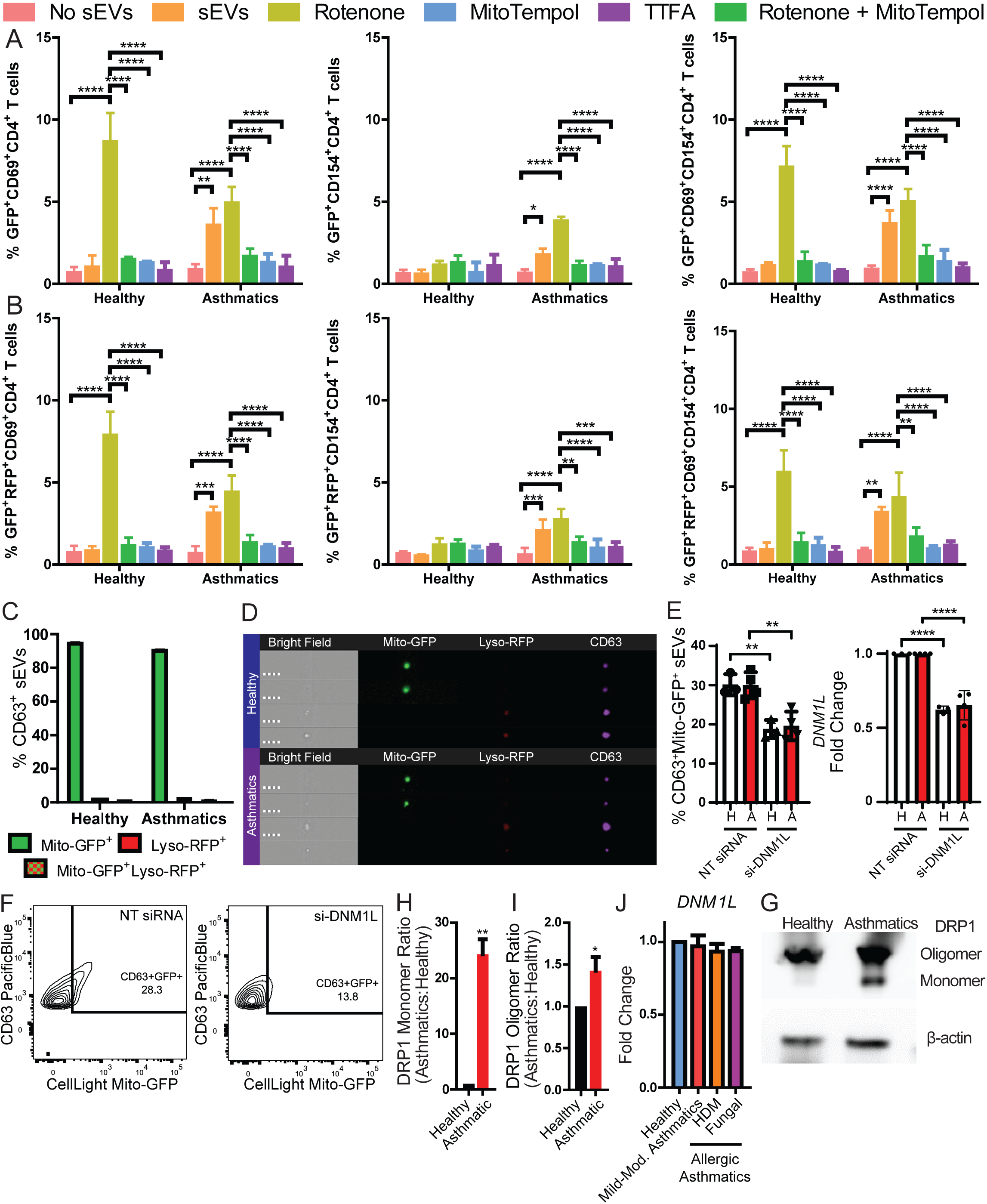
Reduction of mitochondrial ROS within sEVs by MitoTEMPOL inhibits activation of CD4^+^ T cells by MDRC-derived Mito-GFP^+^ sEVs from asthmatics. Packaging of mitochondria inside MDRC-derived sEVs is mediated by mitochondrial fission. (A-B) MDRCs were transduced with CellLight Mito-GFP and CellLight Endo-RFP, and sEVs were purified from the supernatant 48 hours later. sEVs derived from MDRCs transduced with Mito-GFP and Endo-RFP were pre-treated with rotenone (10 nM), MitoTEMPOL (1 µM), rotenone + MitoTEMPOL, thenoyltrifluoroacetone (TTFA, 10 µM) overnight, or untreated prior to co-culture. sEVs were washed and re-purified using the Invitrogen Total Exosome Isolation kit. MDRC-derived sEVs were co-cultured with autologous peripheral CD4^+^ T cells for 24 hours in the presence of rhIL-2 (50 IU/ml) in a ration of 1:10 T cells:sEVs. ImageStream flow cytometry was used to assess early activation (CD69) and antigen-specific activation (CD154). (A-B) Graphs illustrating the percentage of CD69^+^, CD154^+,^ or CD69^+^CD154^+^ T cells that internalized (A) Mito-GFP^+^ MDRC sEVs, (B) both Mito-GFP^+^ and Endo-RFP^+^ MDRC sEVs. Mann Whitney T, *<0.05, **<0.01, ***<0.001, ****<0.0001. (C-D) MDRCs were transduced with CellLight Mito-GFP and CellLight Lyso-RFP, and sEVs were purified from the supernatant 48 hours later. Purified sEVs were characterized using ImageStream flow cytometry to assess if the Mito-GFP and Lyso-RFP co-localized. (C) Quantitation of CD63^+^ MDRC sEVs that are Mito-GFP^+^Lyso-RFP^-^, Mito-GFP^-^Lyso-RFP^+^, or Mito-GFP^+^Lyso-RFP^+^ (D) Representative image strips from ImageStream analysis illustrating Mito-GFP^+^Lyso-RFP^-^ MDRC sEVs, and Mito-GFP^-^ Lyso-RFP^+^ MDRC sEVs. (E-F) MDRCs were transfected with either non-target (NT) control siRNA, or siRNA that targets DNM1L using lipofectamine 3000, and cultured for 48 hours in 10% human serum media that has been depleted of EVs including sEVs. Culture supernatant was isolated for sEV isolation, and cells for gene expression analysis of *DNM1L* (*ACTB* as reference). (E) Left, quantitation of percent CD63^+^Mito-GFP^+^ MDRC sEVs (n=3 Healthy, n=4 Asthma). Right, fold-change of *DNM1L* (Drp1) expression in transfected MDRCs to confirm siRNA efficacy(n=3 Healthy, n=4 Asthma). One way ANOVA, **p=0.006 for comparisons between healthy samples, **p=0.004 for comparison between asthma samples, ****p<0.0001. (F) Representative flow plots illustrating reduction of CD63^+^Mito-GFP^+^ MDRC sEVs after transection with siRNA for *DNM1L*. (G) Protein level and oligomerization of Drp1 was assessed in HLA-DR^+^ MDRCs sorted from the BAL of healthy and asthmatics by Native-PAGE with β-actin as control. (samples were pooled from n=3/study group, quantitation from n=3 western blots) **p<0.01 Mann Whitney test (H) Quantitation of Drp1 monomer ratio in samples from asthmatics to healthy. (I) Quantitation of the Drp1 oligomer ratio in samples from asthmatics to healthy. *p<0.05 (J) Quantitation of real-time PCR analysis of *DNM1L* gene expression in purified HLA-enriched peripheral myeloid cells from healthy, mild-to-moderate asthmatics, and allergic asthmatics (house dust mite or fungal).

Since epithelial cells are the first responders to environmental cues that induce oxidative stress and influence sEV production, we investigated if their sEVs modulate CD4^+^ T cell activation. Although CD45^neg^ cells produced Mito-GFP^+^ sEVs and T cells internalized these sEVs, T cell activation was not observed (Figure S9A-S9D). Therefore, sEVs derived from MDRCs or other antigen presenting cells (APCs), are important in facilitating sEV-mediated T cell activation.

### NF-KB signaling is enhanced in CD4 T cell co-cultures with Human BALF sEVs treated with rotenone, a Complex I inhibitor

As CD4 T cell activation required ROS and as Complex I inhibitor, rotenone promoted T cell activation, we evaluated signaling mechanisms influenced by ROS that underlie T cell activation and polarization. NF-κB family proteins are transcription factors that are important in inflammation and ROS impacts NF-kB signaling; NF-kB targets also may regulate levels of ROS^52^. Canonical NF-κB members, RelA(p65) and c-Rel, have a central role in mediating TCR signaling and T-cell activation^53^. NF-κB also regulates T-cell differentiation and effector function through proinflammatory cytokines^54, 55^, with RelA recently been identified, as a requirement for Th17 polarization^56, 57^. We evaluated sEV mediated effects on NF-kB signaling in presence or absence of rotenone. In CD4^+^ human T cells that internalized MitoTrackerGreen^+^ sEVs (Figure S10 A-C), the proportions of NF-kB p65(pS529)^+^ cells increased in presence of rotenone more in asthmatics compared to healthy subjects (Figure S10 D-E). RelA expression was also increased in T cells of asthmatics in presence of rotenone (Figure S10F). These data suggest that reverse electron transport (RET) does not mediate these effects, since rotenone inhibits RET. Alternative mechanisms are not clear at this point, but could include the impact due to specific sites of mitochondrial ROS production on RelA activation or accumulation of TCA substrates capable of mediating cell signaling^58^.

### Mitochondrial fission is required for sEV packaging of mitochondria

We explored if mitochondrial dynamics is involved in packaging mitochondria within sEVs. We first investigated mitophagy, which is shared with endosomal sorting complexes required for transport (ESCRT) pathway^59^. We transduced MDRCs with CellLight Mito-GFP and Lyso-RFP, a baculovirus construct that encodes lysosomal localization of RFP protein. The Mito-GFP^+^ and Lyso-RFP^+^ signals were mutually exclusive in the purified sEVs, indicating that Mito-GFP^+^ sEVs were not generated by mitophagy (Figure 5C-5D).

Mitochondria may be packaged in sEVs by dynamin-dependent mitochondrial fission, which is regulated by Drp1. We transduced MDRCs with CellLight Mito-GFP, transfected them with either non-target control siRNA or siRNA targeting *DNM1L*, the gene encoding Drp1, and confirmed knockdown of *DNM1L* by gene expression analysis (Figure 5E right panel). The proportion of Mito-GFP^+^ sEVs was significantly reduced following *DNM1L* knockdown (Figure 5E left panel & 5F). As dynamin proteins, such as Drp1, oligomerize to function as facilitators of mitochondrial fission, we compared Drp1 oligomerization in HLA-DR^+^ MDRCs from healthy individuals and asthmatics. Reduced levels of Drp1 monomers and oligomers were noted in HLA-DR^+^ MDRCs from healthy individuals compared to asthmatics (Figure 5G-5I). Gene expression of DNM1L was not significantly different comparing healthy, mild to moderate, and allergic asthmatics (Figure 5J). This enhanced mitochondrial fission in MDRCs from asthmatics may alter Drp1 oligomerization (Figure 5G) and promote packaging of mitochondria in sEVs and elicit sEV-mediated activation in CD4^+^ T cells.

### Mito-GFP^+^ MDRC-derived sEVs internalize at the site of cytoskeletal polarization and mitochondrial trafficking in CD4^+^ T cells

We determined if the internalization of Mito-GFP^+^ sEVs occurs at the CD4^+^ T cell immune synapse, where cytoskeletal structures polarize with the mitochondria at the site of TCR engagement. When PKH-26 labeled MDRC-derived Mito-GFP^+^ sEVs were co-cultured with autologous peripheral T cells, the Mito-GFP^+^ sEVs co-localized with both α-tubulin marking the CD4^+^ T cell cytoskeleton and the polarized CD4^+^ T cell mitochondria (Figure 6A). In CD4^+^ T cells labeled with both α-tubulin and Mitoview (labels mitochondria), the Mito-GFP^+^ sEVs co-localized with mitochondria and cytoskeleton (Figure 6B). Confocal analysis of phalloidin and 4′,6-diamidino-2-phenylindole (DAPI) labeled CD4^+^ T cells co-cultured with Mito-GFP^+^ sEVs, revealed both intracellular and membrane co-localization of the sEVs with actin (Figures 6C & S11). A z-stack slice of the confocal images indicated co-localization (yellow) of the Mito-GFP^+^ sEVs with cytoplasmic actin in the T cells (Figure S11 & Movie SV1). Live confocal imaging of Tubulin-RFP transduced and Mitoview-labeled T cells, co-cultured with Mito-GFP^+^ sEVs also showed co-localization of Mito-GFP with the polarized cytoskeleton and mitochondrial network of the recipient CD4^+^ T cells (Figure 6D & Movie SV2). These data suggest that sEVs are internalized at the site of the immune synapse, which may aid in the organization of the synapse through cytoskeletal and mitochondrial reorganization.

**Figure 6.**
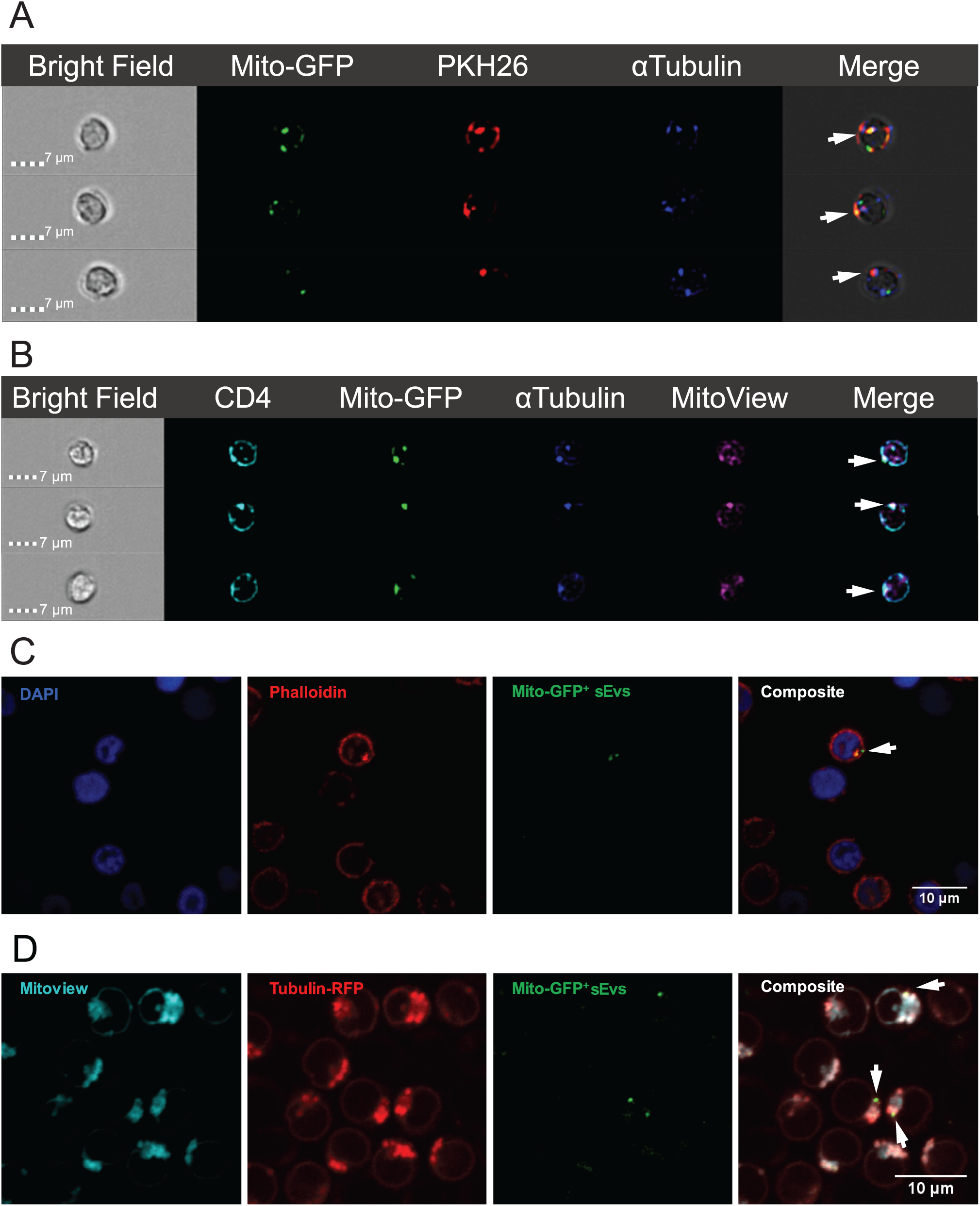
Internalization of Mito-GFP^+^ MDRC sEVs occurs at site of tubulin, actin, and mitochondrial polarization in T cells. (A) Representative images from ImageStream flow cytometry analysis illustrating polarization of α-tubulin at the site of internalization of Mito-GFP^+^ sEVs. MDRCs were transduced with CellLight Mito-GFP and sEVs were purified from the supernatants 48 hours later. Mito-GFP^+^ MDRC sEVs were membrane labeled with PKH26 red dye. T cells were transduced with CellLight Tubulin-RFP 24 hours prior to co-culture with sEVs. MDRC sEV co-culture with T cells (1:10 T cell:sEV) was carried out for 24 hours and analyzed on the ImageStream flow cytometer. (B) Representative images from ImageStream flow cytometry analysis illustrating polarization of α-tubulin and host mitochondria to the site of MDRC sEV internalization. T cells were transduced with CellLight Tubulin-RFP and labeled with MitoView Deep Red dye 24 hours prior to co-culture with MDRC sEVs. sEVs derived from MDRCs transduced with Mito-GFP were co-cultured with T cells (1:10 T cell:sEV) for 24-hours and analyzed on the ImageStream flow cytometer. (C) Confocal image illustrating internalization of MDRC sEVs inside the cytoplasm of the CD4^+^ T cells. sEVs derived from MDRCs transduced with Mito-GFP were co-cultured with T cells (1:10 T cell:sEV) for 24-hours in the ibidi µ-dish. Cells were fixed with 2% PFA and permeabilized with 0.5% triton-X in PBS. The fixed T cells were labeled with Phalloidine-Rhodamine for actin, and DAPI for the nucleus. Cells were imaged on the Nikon A1 confocal. Experiments were performed in n=6-8/study group. (D) Confocal image illustrating polarization of α-tubulin and host mitochondria to the site of MDRC sEV internalization. T cells were transduced with CellLight Tubulin-RFP and labeled with MitoView Deep Red dye 24 hours prior to co-culture with sEVs. sEVs derived from MDRCs transduced with Mito-GFP were co-cultured with T cells (1:10 T cell:sEV) for 24 hours in the ibidi µ-dish and imaged on the Nikon A1 confocal.

### Mito-GFP^+^ MDRC-derived sEVs drive Th2, Th17, and Th2/17 polarization

As BALF sEVs drive Th2 and Th17 polarization (Figure 1E-1I, S3C & S4), we assessed if airway MDRC-derived Mito-GFP^+^ sEVs directly promote Th polarization. Flow cytometry analyses showed IL-4 producing Th2 and IL-17 producing Th17 responses only in T cells that have internalized Mito-GFP^+^ and not the Endo-RFP^+^ sEVs alone (Figure 7A-7F & S12). Additionally, healthy MDRC-derived Mito-GFP^+^ sEVs induced IFN-γ producing Th1 responses (Figure S13A-S13C & S13G). Th2/Th17 hybrid cells that produce both IL-4 and IL-17 are also potent inducers of airway inflammation in asthmatics^11, 60^. The MDRC-derived sEVs from asthmatics induced Th2/17 hybrid responses only in T cells that internalized Mito-GFP^+^ and not Endo-RFP^+^ sEVs (Figure S13D-S13F & S13H).

**Figure 7.**
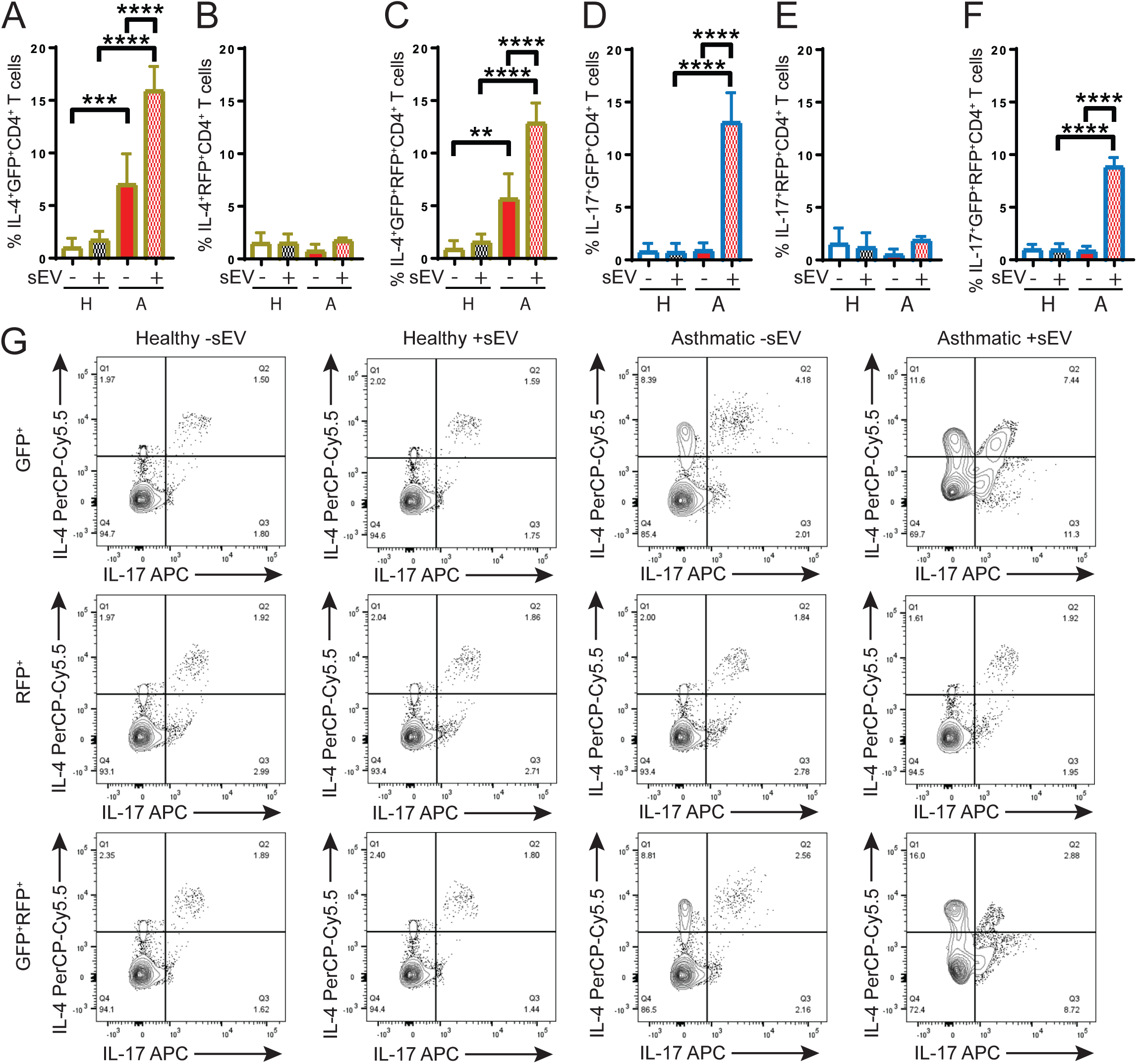
T cells that internalize MDRC-derived Mito-GFP^+^ sEVs from asthmatics polarize to Th17 and Th2 subsets. Mito-GFP^+^ sEVs from asthmatics may activate immune cells through DAMP signaling. sEVs from asthmatics induced pro-inflammatory cytokine production by THP-1. (A-F) Airway MDRCs were transduced with CellLight Mito-GFP and CellLight Endo-RFP, and sEVs were purified from the supernatant 48 hours later. Purified MDRC sEVs were co-cultured with autologous peripheral CD4^+^ T cells for 7 days in the presence of rhIL-2 (50 IU/ml) in a ration of 1:10 T cells:sEV. T helper subsets were assessed by flow cytometry 7 days later by staining intracellular for IL-4 (Th2) and IL-17 (Th17) cytokines. Cells were gated for CD4^+^GFP^+^IFNγ^-^ cells, and then identified as either IL-4^+^IL-17^-^ or IL-4^-^IL-17^+^. Similar gating strategies were used for CD4^+^RFP^+^cells and CD4^+^GFP^+^RFP^+^cells as shown in Supplementary Figure S12. Experiments were performed in (n=11 for healthy and n=13 for asthmatics). Mann Whitney T test for comparison between healthy and asthmatics, Wilcoxon matched-pairs signed rank test for intergroup (no sEVs vs sEVs), *<0.05, **<0.01, ***<0.001, ****<0.0001. (G) Representative flow cytometry plots of cells gated as above showing %Th2 and % Th17 cells shown as %IL-4^+^ and %IL-17^+^CD4^+^ T cells from the co-cultures with sEVs. Flow plots are shown for these Th cell populations that are GFP^+^, RFP^+^ or GFP^+^RFP^+^ in experimental groups of Healthy -sEV, Healthy + seV, Asthmatic –sEV and Asthmatic +sEV.

### Mito-GFP^+^ MDRC-derived sEVs may elicit both innate and adaptive immune responses

Mitochondrial danger-associated molecular patterns (DAMP) signaling^61, 62^ via the sEV mitochondria may trigger T cell responses. We first determined if T cell activation was HLA-specific or due to allogeneic responses. We co-cultured HLA-DR^+^ MDRC-derived sEVs from asthmatics with CD4^+^ T cells from healthy subjects, and vice versa. Modest CD69 activation without unchanged CD154 was seen in CD4^+^ T cells that have internalized Mito-GFP^+^ or both Mito-GFP^+^ and Endo-RFP^+^ MDRC-derived sEVs (Figure S14A-S14C) compared to autologous co-cultures (Figures 3, 4, S5-S8). To investigate potential mitochondrial DAMP signaling, MDRC-derived sEVs from asthmatics and healthy subjects were co-cultured with a human myeloid cell line, THP-1 for 24 hours. Pro-inflammatory cytokines increased in THP-1 cells treated with sEVs of asthmatics (Figure S14D-S14I), while healthy sEVs reduced MCP-1 levels (Figure S14E). These results indicate that immune activation by sEVs can promote both innate and adaptive immune responses and may not be dependent on mitochondrial DAMP signaling.

### In vivo transfer of lung sEVs with stable mitochondrial network enhanced overall airway inflammation and pathogenic Th responses in allergic recipients

To ascertain that it is not mitophagy-mediated shuttling of mitochondria in sEVs that elicit pathogenic Th responses *in vivo*, we utilized Mito-QC mice. Quenched GFP and stable mCherry signal marks mitophagy in Mito-QC mice^63^. Both purified BALF EVs (Figure S15A-S15D) and MDRC-derived EVs from lungs of asthmatic or OVA sensitized Mito-QC mice (Figure S15E-S15H) were predominantly 65-150 nm with a median size consistent with sEVs. We utilized nanoimaging studies to further confirm the size (Figure S16A) and EV marker distribution of MDRC-derived sEVs (Figure S16B) and observed subpopulations of sEVs co-expressing sEV markers including CD81, CD63 and CD9. The predominant sEVs were either CD63^+^ or CD63^+^CD9^+^ in both study groups. High resolution imaging of MDRC-derived sEVs confirmed the co-expression of sEV markers in both groups (Figure S16C). Cryo-electron microscopy of MDRC-derived sEVs from lungs of both asthmatic and OVA sensitized mice showed lipid bilayered vesicles within the size range of sEVs with electron dense inclusions; multi-vesicular bodies were present in asthma group (Figure S16D-S16E). We observed that the intranasal transfer of these lung MDRC-derived sEVs from Mito-QC asthmatic or OVA sensitized donor mice enhanced overall inflammatory cell infiltration in the lungs, airways and lung draining lymph nodes of asthmatic recipient mice compared to OVA sensitized controls (C57BL/6) (Figure S17A-S17C). Followng transfer of MDRC-derived sEVs, OVA-specific IgE in both BALF and serum was increased along with levels of Muc5AC in asthmatic recipient mice compared to OVA sensitized controls (Figure S17D-S17F). MDRC-derived sEVs isolated from OVA sensitized controls elicited inflammatory responses of much lower magnitude when transferred to OVA challenged mice and little to no response in OVA-sensitized mice (Figure S17A-S17F). BALF differential also showed increased airway infiltration of eosinophils, neutrophils, macrophages and lymphocytes in asthmatic recipient mice of MDRC-derived sEVs (Figure S17G-S17J). The absolute numbers and frequencies of lung MDRCs including Ly6C^+^, Ly6G^+^ and Ly6C^+^Ly6G^+^ subpopulations as well as neutrophils and eosinophils also showed similar higher levels in asthmatic recipient mice following sEV transfer (Figures 8A-8E, S18A-S18E, S19). Importantly, the absolute numbers and frequencies of Th2, Th17 cells (Figures 8F-8G, S18F-S18G) as well as Type 2 innate lymphoid cells (Figures 8I, S18I) also increased in asthmatic recipient mice following sEV transfer compared to relevant controls; the absolute numbers and frequency of Th1 cells declined in mice with asthma (Figures 8H, S18H).

**Figure 8.**
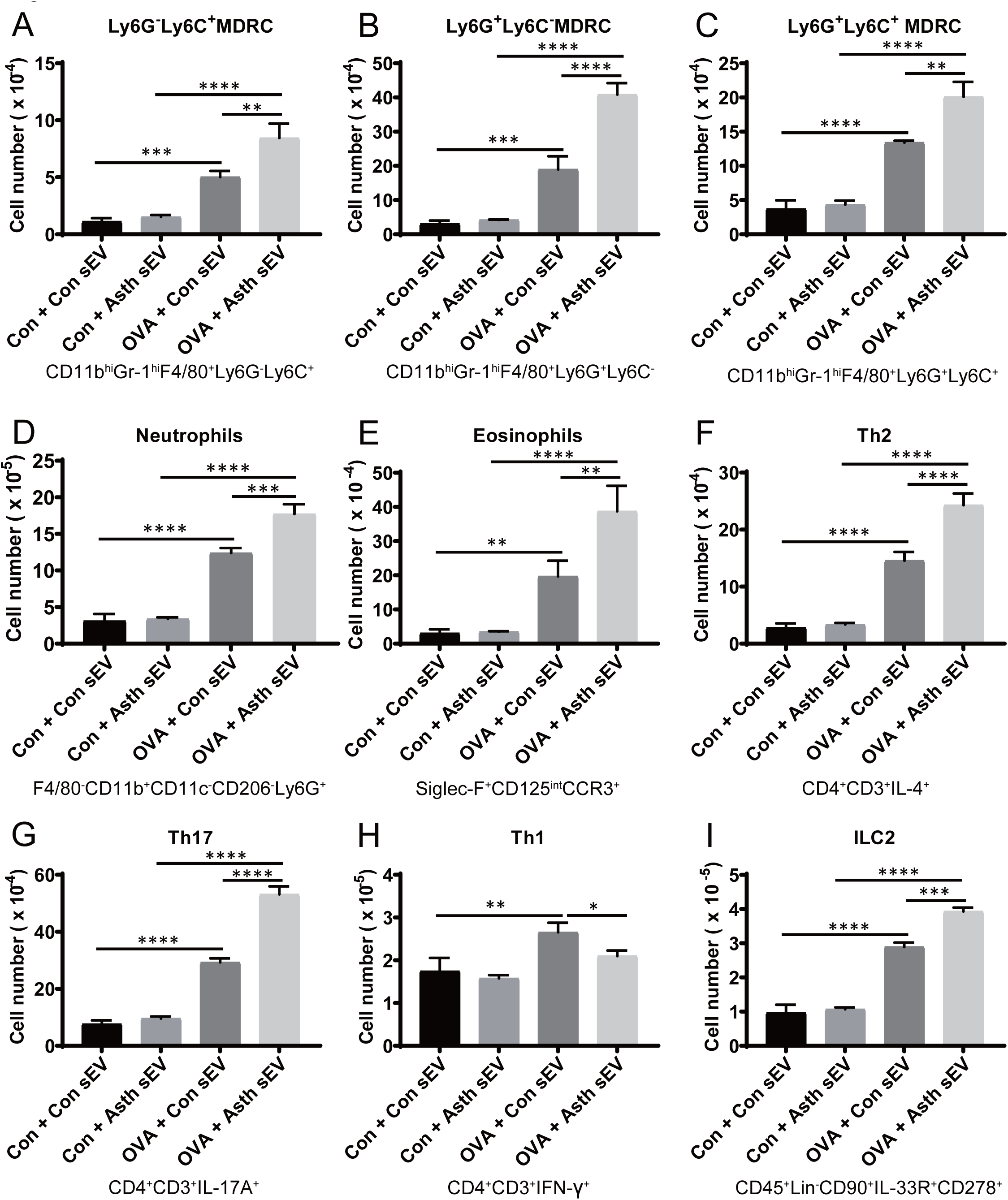
Intranasal transfer of pro-inflammatory lung MDRC-derived sEVs from sensitized and challenged donor Mito-QC mice with asthma exacerbates Th2 and Th17 responses and allergic airway inflammation in sensitized and challenged recipients. Mice were sensitized by intraperitoneal injection on d0 and d7 with 50 *μ*g of alum-adsorbed OVA. On d14, d15 & d16 mice were challenged once i.n. with 15 *μ*g OVA in 30 *μ*l PBS or PBS alone. On d16, i.n. delivery of lung MDRC-derived sEVs (1 x 10^8^ particles/mouse in 30 μl PBS) from control or OVA challenged Mito-QC mice were carried out as before. Lung infiltration of immune cells in lung tissue was determined by FACS analyses. Cell numbers of Ly6G^-^Ly6C^+^ MDRC (A), Ly6G^+^Ly6C^-^ MDRC (B), Ly6G^+^Ly6C^+^ MDRC (C), Neutrophils (D), Eosinophils (E), Th2 (F), Th17 (G), Th1 (H), and ILC2 (I) in the lung tissue were determined by flow cytometry. (n=9 mice/group) Statistical significance was evaluated using one-way ANOVA with Tukey’s multiple comparison testing. ** P < 0.01, *** P < 0.005, **** P < 0.001.

We then confirmed that the activation of CD4^+^ T cells are elicited by internalization of Mito-QC^+^ sEVs *in vivo* during allergic airway inflammation. The oligomerization of Drp1 was enhanced in pro-inflammatory MDRCs sorted from OVA challenged Mito-QC mice compared to sensitized controls (Figure 9A). The frequency of CD81^+^MHCII^+^GFP^+^ sEVs produced by pro-inflammatory MDRCs sorted from OVA challenged Mito-QC mice was higher compared to those from sensitized controls (Figure 9B-9C). Image Stream analyses confirmed the characterization of GFP^+^MHC-II^+^CD81^+^ sEVs isolated from proinflammatory Ly6G^+^ MDRCs (Figure 9C). Further, ImageStream analyses showed that lung-derived CD45^+^CD4^+^CD69^+^ activated T cells were also GFP^+^ in transfer recipient asthmatic mice compared to controls (Figure 9D). Correlation analyses showed that CD69 MFI correlated with % GFP^+^ CD4^+^ T cells as well as % GFP area within T cells of asthmatic recipient mice (Figure 9E-F).

**Figure 9.**
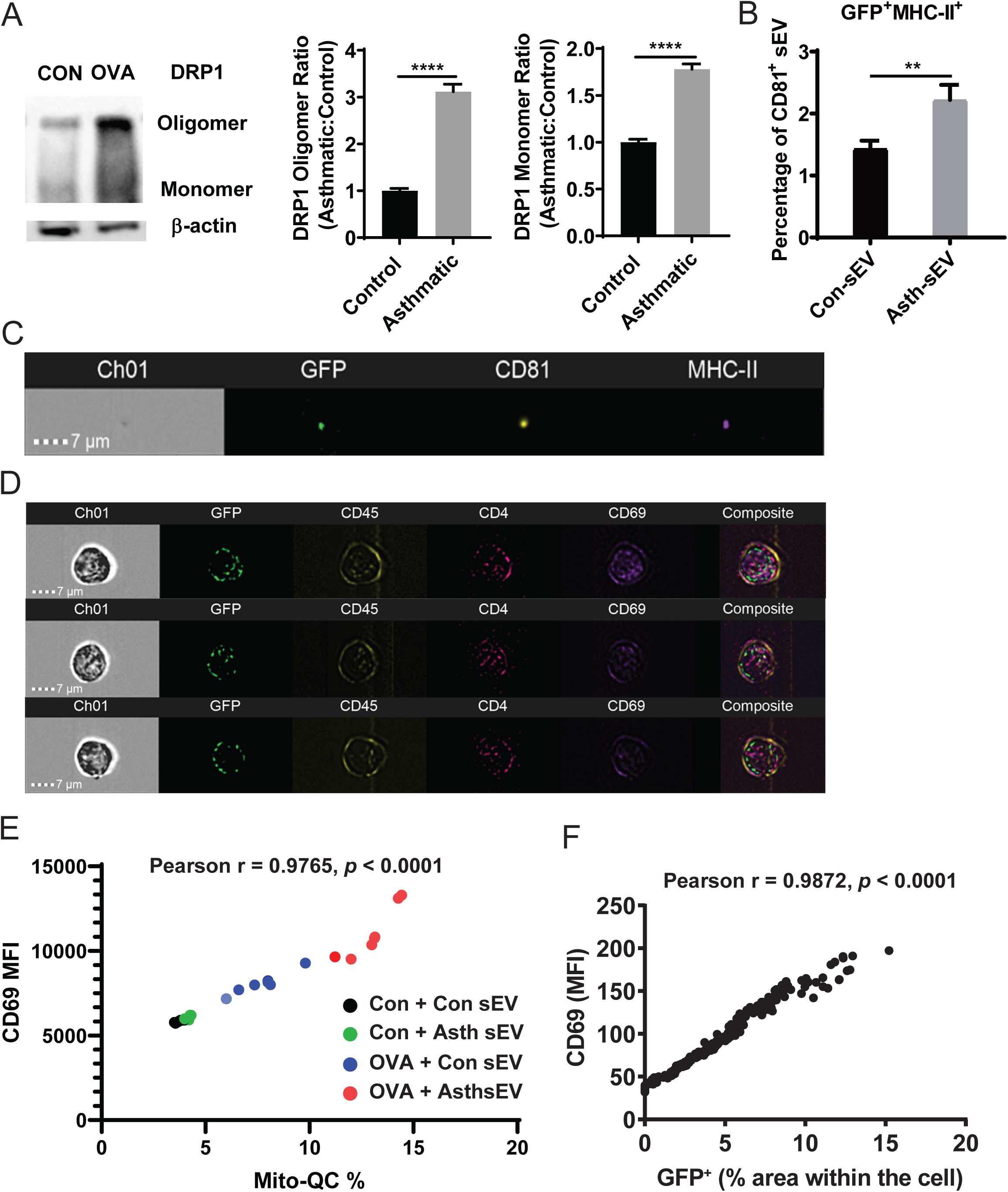
Activation of CD4^+^ T cells by internalization *in vivo* of Mito-QC^+^ sEVs during allergic airway inflammation. (A) Protein level and oligomerization of DRP1 was assessed in Ly6G^+^ MDRCs sorted from the lungs of control or OVA challenged mice by Native-PAGE. The relative expression of DRP1 oligomer or monomer was normalized with β-actin. (B). sEVs were isolated from conditioned media of proinflammatory Ly6G^+^ lung MDRCs sorted from control or OVA sensitized and challenged donor Mito-QC mice using the Total Exosome Isolation kit (n=7 mice/group). Frequency of GFP^+^MHC-II^+^ sEVs in CD81^+^ gate was characterized by ImageStream analyses. (C). Image strips showing GFP^+^MHC-II^+^CD81^+^ sEVs isolated from proinflammatory Ly6G^+^ MDRCs. (D). sEVs isolated from (B) were i.n. delivered (1 x 10^8^ particles/mouse in 30 *μ*l PBS) to recipient sensitized and challenged mice as before. Lung tissue was harvested and digested two days after sEV delivery. Image strips showing co-localization of GFP^+^ signal in CD45^+^CD4^+^CD69^+^ T cells. (E) Pearson correlation analysis between Mito-QC (%) and CD69 expression (MFI) on CD4^+^ T cells. The data were analyzed by IDEAS 6.2 software. The black dots represented Con + Con sEV group (n=5 mice), the green dots represented Con + Asth sEV (n=5 mice), the blue dots represented OVA + Con sEV group (n=7 mice), and the red dots represented OVA + Asth sEV group (n=7 mice). (F) Pearson correlation analysis between Mito-QC (%) and CD69 expression (MFI) on CD4^+^ T cells. The percentage area of GFP^+^ signal or CD69^+^ mean fluorescence intensity (MFI) within each cell, collected from 400 cells using MATLAB analysis. Each data point represents an individual cell. ** *p* < 0.01, **** *p* < 0.0001.

We then investigated if intranasal transfer of these MDRC-derived sEVs from OVA sensitized or challenged donor Mito-QC mice to sensitized or challenged recipients would enhance pathogenic Th responses and exacerbate allergic airway inflammation *in vivo* in recipients. As stable mitochondrial network without mitophagy is marked by binary signal of both mCherry and GFP in Mito-QC mice, we evaluated percentages and absolute numbers of mCherry^+^GFP^+^ Th2 and Th17 cells and found that both were increased in BALF, lung tissues, as well as lymph nodes (Figures 10A-10G, S20-S22) of asthmatic recipient mice compared to control recipents. Consistent with this, IL-4 and IL-17 levels were enhanced in the BALF of the recipient mice (Figure 10H-10I). These data comfirmed that stable mitochondrial network was transferred *in vivo* via sEVs to CD4^+^ T cell subsets to promote Th polarization. Additionally, increased proportion and absolute numbers of mCherry^+^GFP^+^ ILC2s were also noted in BALF of recipient mice with asthma compared to sensitized controls (Figure 10G, S22G).

**Figure 10.**
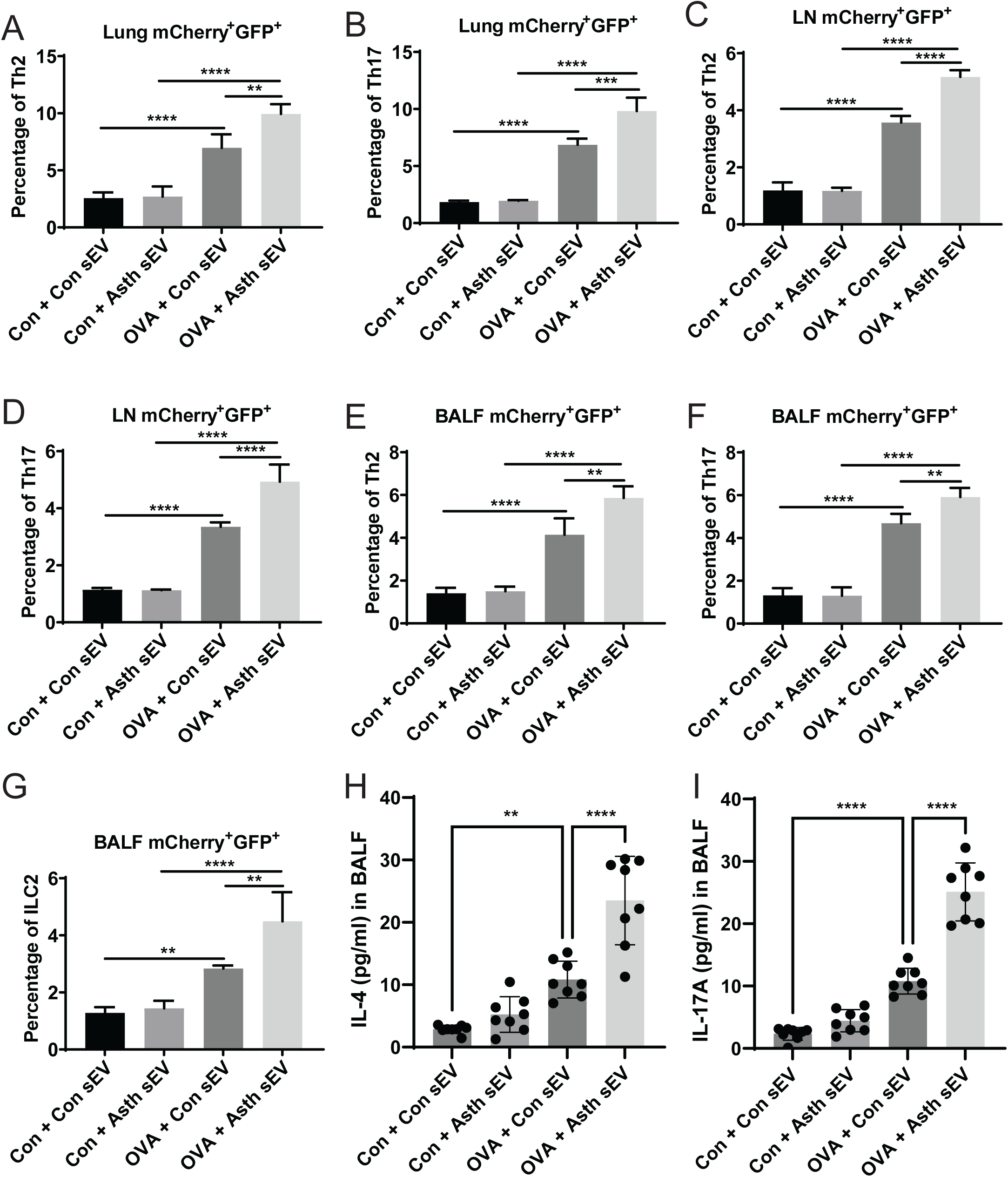
Intranasal transfer of pro-inflammatory lung MDRC-derived sEVs from sensitized and challenged donor Mito-QC mice with asthma enhances mCherry^+^GFP^+^ Th2 and Th17 cell infiltration in the lung tissue, draining LNs and BALF in sensitized and challenged recipients during allergic airway inflammation. Mice were sensitized by intraperitoneal injection on d0 and d7 with 50 *μ*g of alum-adsorbed OVA. On d14, d15 & d16 mice were challenged once i.n. with 15 *μ*g OVA in 30 *μ*l PBS or PBS alone. On d16, 4 hrs after i.n. challenge with OVA in or PBS, i.n. delivery of lung MDRC-derived sEVs (1 x 10^8^ particles/mouse in 30 μl PBS) from control or OVA challenged Mito-QC mice were carried out. Infiltration of immune cells in the lung, draining LNs and BALF were detected by FACS analyses at two days after sEV delivery (n = total 9-11/group). Frequencies of mCherry^+^GFP^+^ Th2 (A) and Th17 (B) in the lung tissue were determined by flow cytometry. Frequencies of mCherry^+^GFP^+^ Th2 (C) and Th17 (D) in the draining LNs were characterized by flow cytometry. Frequencies of mCherry^+^GFP^+^ Th2 (E), Th17 (F) and ILC2 (G) in BALF were determined by flow cytometry. (H) IL-4 levels in BALF determined by ELISA (I) Levels of IL-17A in BALF determined by ELISA (n=8 mice/group). Statistical significance was evaluated using one-way ANOVA with Tukey’s multiple comparison testing. ** *P* < 0.01, *** *P* < 0.005, **** *P* < 0.0001.

## Discussion

Small extracellular vesicles (sEVs) are novel intercellular messengers, both in homeostasis and disease. Our observations extend the role for sEVs as immune modulators in human asthma. We report that airway sEVs from asthmatics drive proliferation of Th cells, and their polarization to Th2 and Th17 phenotypes is supported by NF-kB signaling. Accompanying the changes in epigenetic programming, the airway HLA-DR^+^ MDRC-derived immunogenic sEVs containing mitochondria elicit antigen-specific activation that requires mitochondrial ROS signaling through the transfer of sEV mitochondria to recipient T cells; this transfer involves internalization of sEVs by membrane fusion that initiates TCR engagement/activation in CD4^+^ T cells potentially at the site of the immune synapse. We demonstrate that *in vivo* transfer of stable mitochondrial network carrying sEVs can enhance *in vivo* pathogenic Th polarization accompanied by Th cytokine production in allergic recipients.

CD4^+^ T cells use clathrin-dependent mechanisms to recycle cell surface receptors ^49^; however, transfer of the PKH26 signal from labeled sEVs to CD4^+^ T cell membranes, internalization of the mitochondrial GFP signal and the preservation of sEV internalization by CD4^+^ T cells in the presence of inhibitors of clathrin or dynamin, or LFA-1 antibody, suggest that membrane fusion is the primary mechanism of internalization of mitochondria-containing sEVs. Although CD4^+^ T cell activation correlated with Mito-GFP^+^ sEV internalization, blockade of class II or LFA-1 on sEVs failed to inhibit their internalization, but attenuated activation of recipient CD4^+^ T cells. LFA-1 helps bind T cells to APCs via ICAM-1 and ICAM-2, which aids in antigen presentation^64^ and CD8^+^ T cells utilize LFA-1 to internalize sEVs^65^. Synergistic signaling between class II and LFA-1 may facilitate MDRC-derived sEV-mediated antigen-specific T cell activation without APCs.

Our studies indicate that mitochondria within sEVs are bioenergetically functional and required for sEV-mediated TCR activation. Inhibition of mitochondrial complex II, complex III, and complex V abrogated Mito-GFP^+^ sEV-mediated T cell responses. Interestingly, pre-treatment of MDRC-derived sEVs with complex I inhibitor, rotenone, aberrantly activated healthy T cells (CD69^+^) and promoted antigen specific activation in asthmatics. Since these mitochondrial inhibitors block oxidative phosphorylation, this aberrant activation could be ATP-dependent. However, mitochondria-targeted antioxidant, mitoTEMPOL, scavenged the ROS within MDRC-derived sEVs after rotenone treatment; implicating ROS in the sEV-dependent activation of healthy T cells. Rotenone-induced activation was accompanied by NF-kB signaling and RELA upregulation, both implicated in T cell activation and Th differentiation, suggesting that it is not RET-mediated and may be due to rotenone-mediated ROS production altering normal T cell function^51, 58, 66, 67^. Although low rotenone doses were used on sEVs, and the high affinity of this inhibitor prevents it from leaving sEVs, these concentrations may cause off target effects; mechanisms for rotenone-dependent effects needs to be further clarified^68^. Despite these caveats, our studies implicate mitochondrial ROS as an essential mediator of T cell activation. T cells produce ROS via complex I after TCR engagement^69^, and complex I is important for T helper differentiation/polarization^70^. This is consistent with activation of Th transcriptional program in T cells by mitochondrial signaling via MDRC-sEVs. sEV-mediated activation of Th1 transcriptional regulators may be suppressed by robust activation of Th2 and Th17 programs in T cells from asthmatics and this threshold is different in healthy T cells; this difference may account for Th1 polarization seen in healthy T cells. Taken together, our studies suggest that CD4^+^ T cell-activation by sEVs in the context of asthma or other chronic inflammatory diseases may depend on both TCR-MHC engagement and transfer of functional mitochondria capable of generating ROS.

We interrogated mechanisms underlying mitochondrial packaging within airway myeloid cell-derived sEVs. Mesenchymal stem cells package mitochondria in extracellular vesicles to outsource mitophagy ^36^. Absence of co-localization of the packaged Mito-GFP^+^ signal with Lyso-RFP^+^ associated with mitophagy, demonstrates a novel role for MDRC-derived sEVs in mitochondrial retrograde signaling. Although TLR-mediated activation of immune cells by mitophagy is possible^71^, our results suggest that antigen presentation and mitochondrial ROS signaling are the primary mechanisms of immune activation.

Although transfer of mitochondria by sEVs has been reported^72^, mechanisms of packaging mitochondria within airway immune cell-derived sEVs are largely unknown. We provide insights for mitochondrial fission in the generation of sEVs that transport functional mitochondria. Our observations that intranasal transfer of Mito-QC mice-derived sEVs with binary signal of mCherry and GFP results in exacerbation of allergic airway inflammation and enhancement of pathogenic Th polarization in allergic models, provide *in vivo* evidence for the significance of sEV mitochondria in asthma.

Drp1-dependent mitochondrial fission requires its oligomerization^73^; here, Drp1 oligomerization was increased in MDRCs from asthmatics and Drp1 knockdown reduced Mito-GFP^+^ sEVs. Mitochondrial fragmentation alters immune function in cancer; this limited the function of NK cells^74^. In our studies, mitochondrial fission generates sEVs with mitochondria that facilitates sEV-mediated immune activation, and a role for mitochondrial fission in immune regulation. The extent of sEV signaling is different between healthy controls and asthmatics including ROS and NF-kB-mediated signaling. Genetic differences are, so far, not known to contribute to differences in proinflammatory MDRCs reported in asthmatics. Epigenetic modifications from lung damage/oxidative stress may alter MDRC-derived sEVs in asthmatics and/or differential presence/presentation of antigens by sEVs and/or may account for the increased mitochondrial fission noted in MDRCs from asthmatics; this differential response may not be restricted to asthma and may be important in COPD or sarcoidosis.

Additionally, we demonstrate that Mito-GFP^+^ sEV-mediated signaling and internalization occur in close proximity to the polarized cytoskeleton of activated T cells often associated with synapse formation in activated T cells, which provides an antigen-specific dimension to this cellular crosstalk. Our observation that polarized cytoskeleton and the mitochondria of the activated CD4^+^ T cell co-localizes with sEV mitochondria from donor MDRCs suggest that upon TCR engagement by the MHC-II on sEVs, reorganization of the immune synapse may occur at the site of sEV internalization.

Here, we uncover a novel mechanism by which sEVs activate CD4^+^ T cells, and mitochondrial transfer may function as a second signal in CD4^+^ T cell activation. Traditionally, T cell activation/polarization requires antigen presentation by APCs, co-stimulation and an optimal cytokine milieu^75-77^. Our studies support an acellular form of T cell activation by sEV mitochondria and a new mechanism of MDRC-T cell crosstalk *in vitro* and *in vivo* at the intersection of bioenergetics and cellular function. We propose that these intricate immunological mechanisms facilitated by sEVs participate in both normal physiological responses and may be dysregulated in pathological processes such as human asthma.

## Supporting information

Supplementary Media Legends

## Methods

**Please see supplementary information**

## Acknowledgements

We thank Prof. Ian Ganley, University of Dundee, UK for allowing us to access Mito-QC mice under a Material Transfer Agreement. Genotyped mice were provided by Dr. Martin Young’s laboratory at UAB. Mice were maintained and utilized for *in vivo* experiments according to the MTA guidelines. We gratefully acknowledge Dr. Regan Moore, Oxford Nano imaging, for collaborations for nanoimaging experiments with both human and MDRC-derived sEVs and for analyses of all nanoimaging data. We gratefully acknowledge Marion Spell at the UAB Flow Cytometry Core Facilities, Shawn Williams and Robert Grabski at the High Resolution Imaging Facility, James Kizziah at the Cryo-EM Core Facility, and Sagar Hanumanthu at the Shelby Comprehensive Flow Cytometry Core Facility for technical assistance in sEVs flow cytometry, confocal microscopy, and ImageStream flow cytometry, respectively.

