## Supplementary Media Legends for "Small Extracellular Vesicle Signaling and Mitochondrial Transfer Reprograms T Helper Cell Function in Human Asthma"

**Supplementary Media: Titles and Legends**

**SV1: MDRC-derived Mito-GFP$^+$ sEVs co-localize with actin in activated T cells of subjects with asthma.** A z-stack slice of the confocal images indicated co-localization (yellow) of the Mito-GFP$^+$ sEVs with cytoplasmic actin (red) in the T cells.

**SV2: MDRC-derived Mito-GFP$^+$ sEVs co-localize with polarized cytoskeleton and mitochondrial network of recipient CD4$^+$ T cells.** Confocal images of Tubulin-RFP transduced and Mitoview (labels mitochondria) labeled T cells, co-cultured with MDRC-derived Mito-GFP$^+$ sEVs showing co-localization of Mito-GFP$^+$ sEVs with the polarized cytoskeleton and mitochondrial network of the recipient CD4$^+$ T cells.
