## Supplementary Methods for "Small Extracellular Vesicle Signaling and Mitochondrial Transfer Reprograms T Helper Cell Function in Human Asthma"

**Human Subjects**

Asthmatic and healthy control subjects were enrolled through the University of Alabama at Birmingham Lung Health Center and screened for IgE titer, blood eosinophil frequencies, past medical history, and FEV_1_ (Table I). Healthy controls did not have histories of asthma, pulmonary infections or other known lung diseases. All asthmatic patients had a prior diagnosis of asthma and demonstrated a 12% or greater increase in FEV_1_ within 30 minutes of administrating 400 μg of albuterol, as outlined in the GINA guidelines (Global Strategy for Asthma Management and Prevention; <http://www.ginasthma.org/>) ^1^. Screened subjects with serum cotinine levels greater than 10ng/ml (smokers) and those subjects who received treatment with inhaled or systemic corticosteroids within the six weeks prior to or during the study were excluded. Previous exposure to secondhand smoke was determined by LC-MRM mass spectrometry for serum cotinine levels between 0.05-10 ng/ml ^2^. Healthy controls and asthmatics with serum cotinine levels below 0.05 ng/ml were considered non-SHS-exposed subjects.

Bronchoscopies were carried out as previously described ^3, 4^. In brief, a total of 200 ml of saline solution was instilled, with an average return of 123.3±39.09 ml among healthy, and 92.14±32.50 ml for asthmatics. Eighty (80 ml) of BAL fluid (BALF) was used for isolation of small extracellular vesicles (sEVs). A total of 32 volunteer subjects were assessed in this study, of which 17 in the healthy study arm, and 15 in the asthma study arm. The median IgE titer was significantly higher in the asthma study arm (Table 1). The study was approved by the University of Alabama at Birmingham Institutional Review Board (Protocol IRB-151209005), and written informed consent was obtained from all participants. All procedures performed in this study were performed in accordance with relevant guidelines and regulations.

**Primary Cell Culture**

Primary cells isolated from BALF or peripheral blood were cultured in RPMI 1640 supplemented with 1% Penicillin-Streptomycin, 1% L-Glutamine, and 10% human AB heat inactivated human serum. sEV depleted media was used (described in method details) when cells were cultured for sEV isolation or when sEVs were co-cultured with T cells. For primary T cell co-cultures, recombinant human IL-2 was supplemented at a final concentration of 50 IU/ml.

**Isolation of small extracellular vesicles from BALF**

sEVs were isolated using a previously described differential centrifugation method ^5, 6^. Briefly, 40 ml of BALF was centrifuged at 300 x g for 10 minutes at 4˚C to remove airway cells. Then the supernatant was further centrifuged at 2,000 x g for 10 minutes at 4˚C to remove any dead cells and large cellular debris. The supernatant was spun again at 10,000 x g for 30 minutes at 4˚C to remove any smaller cellular debris and finally filtered through a 0.2 μm cellulose acetate filter (Corning, cat# 430320). The filtrate was then centrifuged at 100,000 x g for 70 minutes at 4˚C and the pellet washed with fresh PBS to remove any contaminating proteins. Finally, the washed pellet was centrifuged at 100,000 x g for 70 minutes at 4˚C, and the final pellet resuspended in 100 µl of fresh PBS. The purified sEVs were stored at -80˚C.

**Isolation of sEVs from human airway HLA-DR^+^ MDRCs**

Human airway HLA-DR^+^ myeloid-derived regulatory cells (MDRCs) were sorted by FACS from the BALF of healthy and asthmatic volunteers as described before ^3^. Cells isolated from the BALF were first blocked with RPMI 1640 media containing 10% Corning Human AB heat inactivated human serum (Corning, NY) for 30 minutes on ice and then stained with the following antibodies: CD11b APC Cy7 (ICRF44, BD Biosciences, Franklin Lakes, NY), CD169 BV510 (7-239, BD Biosciences), HLA-DR APC (LN3), CD163 PE (eBioGHI/61), CD33 PE-Cy7 (WM53), CD14 Percpcy5.5 (61D3) and CD11c PECy5 (3.9) (all other antibodies from eBioscience, Waltham, MA). The pro-inflammatory HLA-DR^+^ MDRCs were identified as CD11b^+^HLA-DR^+^CD33^+^CD11c^+^CD163^+^CD14^lo^CD169^lo^ and sorted using a BD FACS Aria. Sorted cells were cultured in sEV-depleted human AB serum media for 48-hours. The conditioned media was collected and centrifuged twice at 2,000 x g to remove any cell pellets and apoptotic bodies. sEVs were purified from the supernatant using the Total Exosome Isolation Reagent for Cell Culture Media kit (ThermoFisher, Waltham, MA) following the manufacturer’s protocol. Purified sEVs were stored in 50 µl of PBS at -80˚C.

**Preparation of sEV-depleted media**

sEV-depleted media was prepared by supplementing RPMI 1640 media with 20% Corning Human AB Serum (Corning, NY) and centrifuged on an ultracentrifuge overnight at 100,000xg at 4˚C. The supernatant was transferred to a new tube and sterile filtered through a 0.2 μm cellulose acetate filter (Corning, New York). The sEV depleted media was then diluted 1:1 with RPMI to make a final concentration of 10% Human AB Serum. Additionally, the sEV depleted media was supplemented with 1% Penicillin-Streptomycin and 1% L-Glutamine. Media was stored at 4˚C until use.

**Quantitation of sEVs**

The concentration and size distributions of purified airway sEVs were determined using a NanoSight NS300 (Cambridge, MA). The instrument was calibrated using 100 nm polystyrene latex microspheres (cat #NTA4088). The sEVs were diluted 1,000-fold with PBS to make a final volume of 1 ml and loaded in to a 1 ml syringe, which was placed on a syringe pump attached to the NanoSight. The diluted sEVs were injected into the NanoSight at a flow rate of 25 μl/s at room temperature. A total of 5 videos were acquired per sample under the following conditions and settings: temperature: 22.4-22.6°C; viscosity 0.939-0.944 cP; camera level: 7; capture duration: 1 min/video; shutter speed: 11.12 ms; camera type: SCMOS; slider gain: 250; slider shutter: 250; minimum tracks completed: 2000-4000/video; frames processed: 1951/video; frames per second: 32.5 fps; blur: auto; detection threshold: 5.

sEV (human BALF sEVs, murine BALF sEVs and murine MDRC-derived sEVs from MitoQC mice) particle measurements and size distributions were performed using a Spectradyne’s nCS1 nanoparticle analyzer (Signal Hill, CA) with Hardware Version 1. The microfluidic system is primed with a solution of 1% Tween20 in PBS. To prepare a clean diluent (PBS) that does not generate false positive counts, diluents were filtered through a 0.01 filter membrane. C-400 or TS-400 cartridges (65-400nm range) were used to determine the concentration spectral density (CSD) of particles. For each measurement, 3 µl of sample was used. All experiments consisted of acquisitions of n>3000 particles (Error < 3.2), and experiments were repeated in multiple biological and technical replicates with new cartridges to quantify the cartridge variability. After each measurement was completed, data were combined, and (i) peak filters and (ii) background subtraction were applied as recommended by the manufacturer, in the nCS1 Data viewer. To avoid false positive counts due to electrical noise, background subtraction was applied to all measurements. To count the concentrations, particle size distribution, mean or width of the peak of particles over a certain range was obtained after plotting the combined CSD file. For the quantification the size range of interest was selected manually in the Viewer software and then “concentration” option was selected to report the concentration over the selected size range. Peaks in the combined CSD file were fitted to a normal distribution to quantify the particle size distribution. This was done in the Viewer software by selecting the size range of interest manually, and choosing “Gaussian Fit” to fit the selected region to a normal distribution. Fit results and the concentration of particles over the selected size range were then displayed on the plot. Display options were selected on the Viewer software, to display the D-values over the selected size ranges “D10, D50, D90”. After the independent measurements were completed, CSDs were exported from the nCS1 Data Viewer software.

**Cryo- Electron Microscopy**

Murine MDRC-derived sEV samples (10^8^ particles) were applied to 300 mesh copper lacey carbon grids (Ted Pella, Redding, CA) that had been glow-discharged at 30mA for 25s with a Pelco easiGlow glow discharger (Ted Pella). For each grid, 3mL sample were incubated on the grid for ~2min, the grid was manually blotted, another 3mL sample were added, and the grid was loaded into an Vitrobot Mark IV (Thermo Fisher Scientific, Waltham, MA) for the final blotting process. The Vitrobot was set to 12^o^C and 100% humidity, and the grids were blotted for 4s before plunging into liquid ethane. The samples were imaged using the EPU software on a Glacios 2 electron microscope operated at 200kV and equipped with a Falcon 4i direct detector (Thermo Fisher Scientific) at the UAB Cryo-EM Facility. Images were collected as TIFF files at a nominal magnification of 120,000 x (1.19 Å/pix) with 3mm defocus and either 40 or 60 e-/Å2 total dose. Using the software Fiji, a mean filter with a 3-pixel radius was applied to reduce high resolution noise in the images, and scale bars were added.

**Nanoimaging and quantitation of sEVs**

Nanoimaging and quantitation of sEVs were performed in collaboration with Oxford Nanoimaging Company (ONi, San Diego, CA). Purified human BALF sEVs were captured and stained using the Oxford Nanoimaging EV Profiler Kit 2 with the following modifications. Biotinylated antibodies against CD81, CD63, and CD9 were used to capture the human EVs in the kit’s flow chambers. They were stained using the kit’s lipophilic dye, PanEV markers and an antibody against TOMM20 (Abcam 56783) at a dilution of 1:200 and a secondary antibody also provided by ONi (donkey anti-mouse AZ647) at a concentration of 1:200. The secondary antibody was incubated for 30 minutes after washing the primary antibody away and prior to the post-fixation in the EV Profiler Kit 2 protocol. Imaging was completed using the AutoEV function on the ONi Nanoimager. All analysis was completed in the CODI software. The mouse MDRC-derived sEVs were captured using a solution of poly-L-lysine. They were stained using the EV Profiler Kit 2 with primary conjugated antibodies against CD81, CD63, and CD9. These antibodies were raised in a mouse host and therefore may have higher non-specific background signal. Their reactivity has not been extensively tested in murine samples. Imaging was completed using the AutoEV function on the ONi Nanoimager. Analysis was completed in the CODI software.

**Western Blot analyses of sEVs**

BALF-sEV samples isolated from 7 healthy human study subjects and 7 study subjects with asthma were pooled for a total concentration of 8 x 10^9^ sEVs each for both study groups. These sEV samples were lysed in 10 x RIPA buffer (Cell Signaling, 9806S) and 25 x protease inhibitor cocktail (Protease and Phosphatase inhibitor, Thermo Scientific Cat no A32961). Samples were then sonicated at 30 Amplitude on ice for 10 seconds (QSONICA Sonicator Q55), and then incubated on ice for 15 minutes, mixed with loading buffer and heated at 95°C for 8 min. Samples were then electrophoresed on a 15% acrylamide SDS-PAGE gel (Tris-Glycine). Samples were run into the stacking gel at 60 volts for 30 minutes, and then switched to 100 volts to run samples into the separating gel. The gels were then transferred on to a 0.45 Immobilon PVDF membranes (EMD Millipore, Burlington, MA, USA) which was pretreated in methanol for 2 mins at room temperature. Gels were transferred at 100mA in 20% methanol Tris-Glycine transfer buffer overnight at 4^o^C. The PVDF membranes were then blocked in 5% w/v BSA (Fisher, Cat no BP1600-100) in 1xTBST solution for 1 h at RT. The membranes were then probed with the following primary antibodies; Tim23 (Proteintech, Cat no11123-1-AP, 1:1000 diluted diluted in 5% w/v non-fat milk in 1xTBST); CD81 (Cell Signaling, Cat no 56039T, 1:500 in 5% w/v BSA in 1xTBST) with gently shaking at 4°C overnight in cold room. The membranes were then washed three times with 1X TBST for 10 min each at room temperature and secondary antibody anti-rabbit-HRP secondary Ab (1:5000 diluted in BSA/TBST) was added and incubated with gently shaking for 1 hour at room temperature. Then the membranes were washed three times for 10 minutes each. HRP substrate mixture (A and B solution at 1:1; Immobilon Western Chemiluminescent HRP Substrate, Millipore Cat no WBKLS0500) was then added to the membrane and incubated for 1 min. Membranes were then imaged using GeneSys software in SynGene PXi machine.

**ImageStream analysis of sEVs**

ImageStream combines flow cytometry with fluorescence imaging technology, and can resolve much smaller particles than conventional flow cytometry. CD63 eFlour450 (clone H5C6; Affymetrix, Inc., Santa Clara) was used as a positive marker to distinguish sEVs from other types of sEVs, cellular debris, and calibration beads. Other antibodies used in the study were, HLA-DR APC (clone LN3; Affymetrix, Inc., Santa Clara), CD54 PE (clone HA58; Affymetrix, Inc., Santa Clara), CD9 PE (clone M-L13; BD Biosciences, San Jose, CA), and CD81 PE-Cy7 (clone 5A6; BioLegend, Inc., San Diego, CA). For the PKH26 (Sigma, St. Louis, MO) and CellTrace CFSE (Invitrogen, Carlsbad, CA) experiments, sEVs were labeled with their respective dyes following the manufacturer’s protocols. CFSE labeling was stopped with 3% BSA in PBS instead of FBS, which contains exogenous EVs, such as sEVs. The stained samples were imaged at 60X magnification without extended depth of field (EDF), while acquiring data on channels Ch01, Ch03, Ch06, Ch07, Ch09, Ch11 and Ch12. Appropriate controls, single color stains, and calibration beads were used to adjust spectral compensation and to calibrate the machine. ApogeeBead Mix (Apogee Flow Systems Ltd., Hertfordshire, UK) was used to determine gating of 100 nm sized particles. A total of 5,000 events were acquired, gating on forward- and side-scatters, as well as aspect ratio and area. Channels Ch01 and Ch09 were used as brightfields, and Ch12 was used for side-scatter. The acquired data was analyzed using IDEAS software version 6.2 (EMD Millipore, Bellerica, MA).

**Flow cytometry analysis of sEVs**

A total of 1x10^7^ particles of airway sEVs were stained with antibodies specific for CD63, CD54, HLA-DR, CD9, CD81, and TSG101 Alexa Fluor 647 (clone 4A10; Novus Biologicals, Littleton, CO). Flow cytometry data was acquired on a BD Becton Dickinson LSRII (Franklin Lakes, NJ). This machine was configured to detect small particles with ApogeeBead Mix (Apogee Flow Systems Ltd., Hertfordshire, UK) by changing the photomultiplier tube (PMT) voltage for forward and side scatters to 600 volts and 286 volts respectively, and thresholding on both forward and side scatters to 500 volts. Particles scattering in the same range as beads between 50-150 nm were gated and total of 100,000 events were acquired. PBS with antibodies alone was used as a control to confirm the absence of a background signals due to antibodies in the samples. Fluorescence compensation was configured using single color stains of sEVs. Acquired flow cytometry data was analyzed using FlowJo X.

**ImageStream and Flow Cytometry analysis of CD4^+^ T cells**

Autologous peripheral CD4^+^ T cells were blocked for 30 min in sEV-depleted RPMI supplemented with 10% human AB serum. Cells were fixed and permeabilized using BD Cytoperm/Cytofix Kit (BD, Franklin Lakes, NJ). Cells were then labeled with CD4 PE-Cy7 (clone SK3; ThermoFisher, Waltham, MA), IL-4 PE (clone 8D4-8; ThermoFisher, Waltham, MA), IL-17A APC (clone eBio64DEC17; ThermoFisher, Waltham, MA), IFNγ BV421 (clone B27; BD Horizon, Franklin Lakes, NJ), CD69 eFluor450 (clone FN50; ThermoFisher, Waltham, MA), CD154 APC (clone 24-31; ThermoFisher, Waltham, MA), pZap70 Alexa Fluor 647 (clone 17A/P-ZAP70; BD Phosflow, Franklin Lakes, NJ), or α-tubulin eFluor 615 (clone DM1A; ThermoFisher, Waltham, MA) for 30 minutes on ice and washed twice. For flow cytometry, cells were resuspended in 100 µl of PBS. For ImageStream, cells were resuspended in 30 µl of PBS in a 1.5 ml microcentrifuge tube. PMTs were adjusted using unstained and compensation configured using single color controls.

**Assessment of NF-kB signaling in sEV-CD4^+^ T co-cultures**

Primary human peripheral T cells isolated from healthy and asthmatics (n=5-7/group) were cultured in a sEV free RPMI media (Corning, Product # -15-040-CV) supplemented with 20% FBS (ATLAS, Cat no # F-0500-D), 2% L-Glutamine (Corning, Product # 25-005-Cl), 1% Penicillin-Streptomycin (Cellgro, Cat no # 30-002-CI), For primary T cell co-cultures, recombinant human IL-2 was supplemented at a final concentration of 50 IU/ml and Recombinant IL-2 (5 IU/ul R&D Systems, Lot: AE6015081). Purified BALF sEVs labeled were treated with 10 µM rotenone, a mitochondrial Complex I inhibitor. sEVs were then washed and purified by centrifugation and then labeled with MitoTracker Green, following which they were washed and purified as described before. The treated and untreated sEVs were then cultured for 24 hours with autologous peripheral CD4^+^ T cells at a ratio of 10 sEV per T cell based on concentrations determined from Spectradyne analysis. CD4^+^ T cells were blocked for 30 min in sEV-depleted RPMI media as described above. Cells were then labeled with surface marker CD4 PE-Cy7 117 (clone SK3; Cat# 25-0047-41, ThermoFisher, Waltham, MA), and washed twice with prewarmed FACS buffer. T cells were then fixed (10 min, 37^o^c) with pre-warmed BD Cytofix Fixation Buffer (Cat .No. 554655). Cells were then permeabilized (30 min on ice) with Buffer (eBioscience Cat# 00-5523-00), and washed twice with the FACS Buffer. Then cells were stained with NF-kB p65(pS529) (BD Phosflow Cat#565446 BV421 ) in a permeabilization buffer for 30 min at room temperature. Finally, cells were washed twice and resuspended in 100 μl of PBS for flow cytometry. Percent of CD4^+^ T cells that were MitoTracker Green^+^ and NF-kB p65 (pS529)^+^ were determined from these sEV-T cell co-cultures.

**Labeling of sEVs with PKH26 Red**

The sEVs were labeled with PKH26 (Sigma, St. Louis, MO) in PBS. Final concentration of PKH26 is 2 µM. Samples were stained for 15 minutes at 37˚C in the dark, followed by isolation using Total Exosome Isolation from Cell Culture Media kit (ThermoFisher, Waltham, MA) per the manufacturer’s protocol. Final sEV pellet was resuspended in 50 µl of sterile PBS.

**Labeling of cells with PKH26 Red or MitoView 633**

MDRCs or autologous peripheral CD4^+^ T cells were labeled with PKH26 Red (Sigma, St. Louis, MO) at a final concentration of 2 µM following the manufacturer’s protocols. Samples were stained for 15 minutes at 37˚C in the dark, followed by two washes with pre-warmed sEV-depleted human serum RPMI media. For MitoView 633, cells were live stained with sEV-depleted RPMI with 10% human AB serum with 100 nM of MitoView 633 for 15 minute at 37˚C in the dark. Cells were then washed twice with pre-warmed sEV-depleted human serum RPMI media.

**Transduction of cells with CellLight constructs**

MDRCs or autologous peripheral CD4^+^ T cells were transduced with the CellLight Bacman reagent at a particle-per-cell value of 10 for 1x10^5^ cells/well in one well of a 96-well plate. An exception to this was with the transduction of CD4^+^ T cells with CellLight Tubulin RFP, where the reagent volume was doubled due to lower transduction efficiency. 10 µl of CellLight Bacman reagent (Mitochondria GFP, Early Endosome RFP, or Lysosome RFP) or 20 µl of CellLight Tubulin RFP was used per 1x10^5^ cells/well following the manufacturer’s protocol. The transduction was performed in sEV-depleted human serum RPMI media for 24 hours.

**Co-culture of sEVs with CD4^+^ T cells and inhibition assays**

Purified airway sEVs were cultured with autologous peripheral CD4^+^ T cells at a ratio of 10 sEV per T cell based on concentrations determined from NTA analysis. For sEVs generated from MDRCs, we normalized our conditions so that a ratio of sEV generating cells to CD4^+^ T cells of 2.5x10^5^ MDRCs (sEV generating cells) to 1x10^6^ CD4^+^ T cells was used. The sEV-T cell co-culture was performed for 24-hours in sEV depleted media. Mitochondrial inhibitors used to block complex I, complex II, complex III or complex V were used at the following final concentrations: 10 nM rotenone, 10 µM Thenoyltrifluoroacetone (TTFA), 10 µM antimycin A, 10 µM oligomycin. MitoTEMPOL was used at a concentration of 1 µM. The sEVs were treated with the inhibitors 48 hours prior to co-culture and purified twice using the Invitrogen Cell Culture Exosome Isolation kit to prevent carryover of inhibitors into the co-culture. For LFA-1 antibody (clone R7.1, Fisher Scientific) a final concentration of 1 µg/ml was used. For the Pan-HLA (DR/DP/DQ) antibody (clone TU39, BD Pharmigen) a final concentration of 10 µg/ml. Pitstop2 (Abcam, Cambridge, UK) was used at a final concentration of 50 nM. Dynasore (Sigma, St. Louis, MO) was used at a final concentration of 50 nM.

**Cytokine assessment in sEV-CD4^+^ T cell co-culture supernatants**

Co-culture supernatants collected following co-culture of sEVs isolated from BALF of asthma and healthy study subjects with autologous peripheral CD4^+^ T cells, as described above, were utilized to measure levels of IL-4 and IL-17. The quantikine human IL-17 immunoassay kit (R & D Systems, Cat# D1700) was used to measure levels of IL-17 following manufacterer’s recommendations. The quantikine human IL-4 immunoassay kit ( R & D Systems, Cat # D4050) was used to measure IL-4 levels following manufacturer’s recommendations.

**NanoString gene expression analysis of CD4^+^ T cells**

Autologous peripheral CD4^+^ T cells were cultured with or without sEVs for 7 days in sEV-depleted RPMI supplemented with 10% human AB serum and 50 IU/ml rhIL-2. Cells were harvested and processed for RNA purification using Invitrogen PureLink RNA Mini Kit (ThermoFisher, Waltham, MA). NanoString (Seattle, WA) differential gene expression analysis was conducted using a custom panel of 77 genes at the NanoString core facility at the University of Alabama at Birmingham. Differential gene expression analysis was performed using nSolver version 3 (NanoString, Seattle, WA), R, and Metaboanalyst 3.0 (Xia Lab, McGill University, Montreal, Canada).

**Real-time qPCR analysis of CD4^+^ T cells**

Autologous peripheral CD4^+^ T cells were cultured with or without sEVs for 7 days in sEV-depleted RPMI supplemented with 10% human AB serum and 50IU/ml rhIL-2. Cells were harvested and processed for RNA purification by Trizol. Purified RNA was reconstituted in nuclease-free water and the concentration quantitated. cDNA was synthesized using the PrimeScript 1^st^ Strand Synthesis Kit (Takara Bio, Mountain View, CA). The VeriQuest SYBR qPCR Master Mix with ROX (ThermoFisher, Waltham, MA) was used to prepare the qRT-PCR reactions with 100 ng of cDNA. ACTB was used as a reference gene. Gene expression of IL-4 and IL-17 was measured (primers and sequences listed in Key Resources Table). The qRT-PCR was performed on a StepOne system (ThermoFisher, Waltham, MA). For evaluation of gene expression of RELA, GATA-3 and RORC, total RNA from cells was isolated using the Takara isolation kit (NucleoSpin@RNA plus, Cat # 740984.50) and reverse transcribed to cDNA using the cDNA synthesis kit (Takara Cat# RR037A, USA) according to the manufacturer's protocol. Real-time PCR procedure was p erformed with TB Green Premix Ex Tag II (Takara Cat# RR820A, USA). The primer sequences used are listed in rge Key Resources Table. Gene expression data analysis was performed using the 2^-ΔΔCT^ method.

**DNM1L siRNA and qRT-PCR analysis**

MDRCs were transfected with ON-TARGETplus SMARTpool *DNM1L* siRNA (Dharmacon, Lafayette, CO) using Lipofectamine 3000 (ThermoFisher, Waltham, MA) per the manufacturer’s protocol. Cells were transduced with CellLight Mitochondria GFP and cultured for 48-hours in sEV-depleted RPMI supplemented with 10% human AB serum. Supernatants were harvested for sEV isolation and characterization by flow cytometry. Cell pellets were collected for RNA isolation using Trizol (ThermoFisher, Waltham, MA) following manufacturer’s protocol. RNA was quantified and cDNA generated using the Takara Bio PrimeScript 1^st^ Strand cDNA synthesis kit (Takara Bio, Kusatsu, Japan). The VeriQuest SYBR qPCR Master Mix with ROX (ThermoFisher, Waltham, MA) was used to prepare the qRT-PCR reactions with 100ng of cDNA. ACTB was used as a reference gene. The following primers used for *DNM1L* were designed using UCSC genome browser and *in silico* PCR for hg38 (forward: GCTCCAGGACGTCTTCAACA; reverse: TAGCACTGAGCTCTTTCCGC). The qRT-PCR was performed on a StepOne system (ThermoFisher, Waltham, MA).

**Native-PAGE and Western Blot Analyses**

Sorted HLA-DR^+^CD11b^+^ MDRCs from 3 asthmatics and 3 healthy subjects were pooled within each study group. The cells were washed in PBS, and pelleted. The final pellet was resuspended in 50 µl of PBS and sonicated on ice 5 times in short 5 second bursts to minimize heat. Protein was quantitated using the BCA method and 35 µg of sample was loaded on a 4-15% gradient gel (Mini-PROTEAN TGX StainFree; BioRad, Hercules, CA). A non-SDS native gel running buffer was used (1.92 M Glycine, 250 mM Tris-Base, pH 8.3 not adjusted). The gel was run at 100V for 5 hours at 4°C to minimize heat. After electrophoresis of the native-PAGE gel, the protein was transferred to a PVDF membrane. The PVDF membrane was activated with methanol. The transfer occurred in transfer buffer (1.92 M Glycine, 250 mM Tris-Base, pH 8.3 not adjusted) overnight at constant 100mA at 4°C. The transferred membrane was washed with Tris-buffered saline with 0.1% Tween 20 (TBS-T) and blocked in TBS-T with 5% non-fat milk overnight at 4°C. Membrane was probed with primary mouse anti-Drp1 (Cat: 611113, Clone 8; BD, Franklin Lakes, NJ) at a 1:1000 dilution in TBS-T with 5% non-fat milk overnight at 4°C. Membrane was washed with TBS-T 3 times for 5 min at room temperature on a rocker. Secondary antibody (anti-mouse HRP) was used at a dilution of 1:5000 in TBS-T with 5% non-fat milk and membrane probed for 2 hours at room temperature. Blot was washed 3 times with TBS-T and imaged.

**Cytokine and chemokine assay of sEV treated THP-1 cells**

IL-1β, TNF-α, IFN-γ, MCP-1, MIP-1β, MIP-1α, Eotaxin and VEGF in culture supernatant were assayed by a luminex magnetic bead based multiplexed (Biorad Bio-Plex 200 system, Luminex Bio, USA) assay using commercially available kits (Miliplex MAP Human cytokine/chemokine, Cat# HCYTOMAG-60K) according to the manufacturer's recommended protocol.

**Experimental Allergic Airway Inflammation and Intranasal Adoptive Transfer of MDRC-derived sEVs.**

Female C57BL/6J mice at 6 to 8 weeks of age were purchased from The Jackson Laboratory (Bar Harbor, ME). Mice were kept in pathogen-free conditions and handled in accordance with the Guidelines for Animal Experiments at the University of Alabama at Birmingham. Mito-QC mice were obtained from University of Dundee under a Material Transfer Agreement and were maintained and utilized according to the MTA guidelines. These mice express a functionally inert, tandem mCherry- GFP tag fused to the mitochondrial targeting sequence of the outer mitochondrial membrane (OMM) protein, FIS1 (comprising amino acids 101–152). Under steady-state conditions, the mitochondrial network fluoresces both red and green and can monitored mCherry+ GFP+. If there is mitophagy, mitochondria are delivered to lysosomes where mCherry fluorescence remains stable, but GFP signal will be quenched. Mito-QC mice were sensitized by intraperitoneal injection (i.p.) on d0 and d7 with 50 *μ*g of alum-precipitated OVA (Grade VII, Sigma Chemical, St Louis, MO; <1 ng lipopolysaccharide per mg) as previously described. On d14, d15 and d16, under anesthesia with Isoflurane (Schering-Plough Animal Health, Union, NJ), Mito-QC mice were challenged intranasally (i.n.) with 15 *μ*g OVA in 30 *μ*l PBS or PBS alone. On day 18, lung tissues were harvested from both asthmatic and control group. Gr-1^+^CD11b^+^Ly6G^+^F4/80^lo^ proinflammatory MDRCs were sorted and sEVs isolated from the culture supernatant after 24 hour culture. For intranasal transfer of isolated sEVs into mice with asthma, C57BL/6J mice were first sensitized by i.p. injection on d0 and d7 with 50 *μ*g of alum-precipitated OVA. On d14 and d15, C57BL/6J mice were challenged i.n. with 15 *μ*g OVA in 30 *μ*l PBS or PBS alone. On d16, C57BL/6J mice were challenged i.n. with 15 *μ*g OVA in 30 *μ*l PBS or PBS alone in the morning, followed by i.n. delivery of lung MDRC-derived sEVs (1 x 10^8^ particles in 30 *μ*l PBS) from control or OVA-challenged Mito-QC mice in the afternoon. At two days after OVA challenge and sEV delivery, BALF and sera were harvested for OVA-IgE (Biolegend, San Diego, CA) and Muc5AC analyses (MyBioSource, San Diego, CA) by ELISA. Lung tissues were digested with collagenase B (Sigma Chemical, St Louis, MO) and immunophenotyping was carried out as described before.

**Cytokine Assays of mouse bronchoalveolar lavage fluid**

Bronchoalveolar lavage fluid collected from sensitized and challenged mice following intranasal adoptive transfer of sEVs were utilized for assessment of Th2 and Th17 cytokines. Levels of IL-4 was measured using the quantikine mouse IL-4 ELISA kit, (R & D Systems, Cat# M4000B) and IL-17A levels were measured using quantikine mouse IL-17 ELISA kit (R & D Systems, Cat# M1700) following manufacturer’s recommendations.

**Preparation of sEV-depleted media**

sEV-depleted medium was prepared as previously described. Briefly, RPMI 1640 media supplemented with 20% FBS was centrifuged using an ultracentrifuge overnight at 100,000 x g at 4˚ C. Supernatant was filtered through a 0.2 μm cellulose acetate filter (Corning, NY). sEV depleted media were then diluted 1:1 with RPMI 1640 media to make a final concentration of 10% FBS.

**Isolation of MDRC-derived sEVs from conditioned media**

Immunosorted Gr-1^+^CD11b^+^Ly6G^+^F4/80^lo^ MDRCs were cultured in sEV-depleted RPMI 1640 media for 24 hrs. Conditioned media were collected and centrifuged at 2000 × g to remove any cell pellets and apoptotic bodies. The supernatant was then incubated with the Total Exosome Isolation Reagent for Cell Culture Media kit (ThermoFisher, Waltham, MA) per the manufacturer's protocol. Purified sEVs were stored in 50 µl of PBS at −80° C.

**NanoSight particle analysis for quantitation of sEV size and concentration**

The concentrations and size distributions of purified MDRC-derived sEVs were determined using a NanoSight NS300 (Cambridge, MA) as described previously for human sEVs above.

**ImageStream analysis of MDRC-derived sEVs and lung tissue cells after sEV transfer**

ImageStream flow cytometry analysis of sEVs purified from conditioned media was performed as previously described. sEVs were stained with the following antibodies: eFlour450-conjugated anti-mouse MHC-II (M5/114/15.2) and APC-conjugated anti-mouse CD63 (NVG2) antibodies were purchased from Life Technologies (Grand Island, NY). PE-conjugated anti-mouse CD81 (Eat-2), PE-Cy7-conjugated anti-mouse CD9 (MZ3) antibodies were purchased from BioLegend (San Diego, CA). Cells from collagenase digested lung tissue were prepared for flow cytometry as described before. Following preparation of single cell suspension, cells were treated with Fc Block and then stained with the following antibodies. PE-conjugated anti-mouse CD45 (30-F11), PE-Cy7-conjugated anti-mouse CD4 (GK1.5), and eFlour450-conjugated anti-mouse CD69 (H1.2F3) antibodies were purchased from Life Technologies (Grand Island, NY). Unstained control, single color staining, and calibration beads were used to calibrate the machine and adjust compensation. The stained samples were imaged at 60 x magnification with extended depth of field (EDF). The data were acquired on channels Ch01, Ch03, Ch06, Ch07, Ch09, Ch11 and Ch12. Ch01 and Ch09 were used as bright field channels whereas Ch12 was used for side-scatter. A total of 5,000 events were acquired for each sample, with three technical replicates per sample. The acquired data were analyzed using IDEAS software version 6.2 (EMD Millipore, Billerica, MA).

**BALF differential cell count**

BALF was collected for Diff-Quik staining (EMD Millipore, Bilerica, MA) from mice after i.n. adoptive transfer of MDRC-derived sEVs. BALF differential analysis was performed following Diff-Quik staining and assessing standard morphological criteria to quantitate BALF cells on cytospin slides. At least 300 cells were examined in each cytospin slide. Numbers of eosinophils, macrophages, neutrophils and lymphocytes were calculated based on the percentage of each cell population in the slides.

**Flow cytometry**

Lung tissues were harvested and digested with collagenase B. Red blood cells were removed by ACK lysis buffer. Fc receptors were blocked with 3% BSA in PBS containing 2.4G2 antibody (anti-mouse CD16/CD32; BD Pharmingin), followed by staining with the antibodies below. Phycoerythrin (PE)–conjugatedanti-mouse-Gr-1 (RB6-8C5), anti-mouse CD25 (PC61.5), anti-mouse CD62L (MEL-14), allophycocyanin (APC)–conjugated anti-mouse-CD206 (MR6F3), anti-mouse IL-4 (11B11), PerCP-Cyanine (Cy) 5.5 conjugated anti-mouse-Ly6C (HK1.4), anti-mouse FoxP3 (FJK-16s), PerCP-eFluor 710-conjugated anti-mouse CD170 (Siglec F, clone 1RNM44N), PE-Cy5-conjugated anti-mouse MHC-II (I-A/I-E, M5/114.15.2), and PE-Cy7 conjugated anti-mouse-CD4 (GK1.5) antibodies were purchased from Life Technologies (Grand Island, NY). BV605-conjugated anti-mouse-F4/80 (T45-2342), APC-Cy7–conjugated anti-mouse-CD11b (M1/70), anti-mouse-CD3 (145-2C11), Alexa Fluor 700-conjugated anti-mouse-Ly6G (1A8), and Alexa Fluor 647-conjugated anti-Mouse CD101 (Igsf2, 307707) antibodies were purchased from BD Bioscience (San Jose, CA). PerCP-Cy5.5 conjugated anti-mouse IL-33Rα (IL1RL1, ST2, DIH9) and anti-mouse IL-17A (TC11-18H10.1), PE-Cy7-conjugated anti-mouse-CD45 (30-F11), anti-mouse CD4 (GK1.5), anti-mouse CD125 (IL-5Rα, DIH37), Pacific Blue-conjugated anti-mouse lingeage cocktail (including CD3 (17A2), B220 (RA3-6B2), CD11b (M1/70), TER-119 (Ter-119), Gr-1(RB6-8C5)), anti-mouse CD4 (GK1.5), anti-mouse CD8a (53-6.7), anti-mouse CD4 (GK1.5), anti-mouse CD11c (N418), anti-mouse NK1.1 (PK136), anti-mouse FcεRIα (MAR-1), anti-mouse IFN-γ (XMF1.2), Brilliant Violet 421 conjugated anti-mouse CD193 (CCR3, J073E5), APC-conjugated anti-mouse CD90.2 (30-H12), anti-mouse CD127 (A7R34), and APC-Cy7 conjugated anti-mouse CD3 (17A2) and anti-mouse CD278 (ICOS, C398.4A) antibodies were purchased from BioLegend (San Diego, CA). Data were collected with LSR-II flow cytometer (Becton Dickinson) and analyzed with FlowJo software (version 8.5.2; TreeStar, Ashland, OR).

**Quantification and Statistical Analysis**

Statistical analysis was performed using GraphPad Prism 5.04, Metaboanalyst 3.0, or R 3.5.2 (64-bit). NanoString data was formatted to be compatible with Metaboanalyst to perform multivariate statistics using Metaboanalyst’s interactive and intuitive platform. Descriptions of the statistical tests used are elaborated in the figure legends. Significance was defined as a p-value lower than 0.05, and denoted as: * < 0.05; ** < 0.01; *** < 0.001; and **** < 0.0001. Error bars represent standard error of the mean unless otherwise noted in the figure legend as standard deviation.

**Data and software availability**

The NanoString custom panel gene expression data has been deposited in the Gene Expression Omnibus (GEO) under accession ACCESSION# GSE144813 and #GPL28122

**Reagent and resource sharing**

**Key Resources**

| **Reagent or Resource** | **Source** | **Identifier** |
| --- | --- | --- |
| **Chemicals & Reagents & Kits** | | |
| MitoSOX Red Mitochondrial Superoxide Indicator | Thermo Fisher Scientific | Cat: M36008 |
| MitoTracker Green | Thermo Fisher Scientific | Cat: M7514 |
| VeriQuest SYBR Green qPCR Master Mix | Thermo Fisher Scientific | Cat: 75600 |
| PrimeScript 1^st^ Strand cDNA Synthesis kit | Takara Bio | Cat: 6100A |
| Ficoll Paque | GE Healthcare | Cat: 45-001-749 |
| Human CD4^+^ T cells Enrichment Cocktail | STEMCELL Technologies | Cat: 15062 |
| HLA Myeloid Cell Enrichment Cocktail | STEMCELL Technologies | Cat: 15272H |
| RPMI | Corning | Cat: 15-040-CV |
| FBS | Atlas Biologicals | Cat: F-0500-DR |
| Penicillin-Streptomycin | MP Biomedicals, Inc | Cat: ICN1674049 |
| Glutamine | Corning | Cat: MT25005CI |
| Total Exosome Isolation Kit | Thermo Fisher Scientific | Cat: 4478362 |
| Trizol | Thermo Fisher Scientific | Cat: 15596018 |
| Ethanol 100% | Decon Labs | Cat: 2716GEA |
| Isopropanol | Sigma | Cat: I9516-500ML |
| RNAse Free Molecular Grade Water | Fisher Scientific | Cat: BP561-1 |
| Chloroform | Sigma | Cat: 472476-500ML |
| PureLink RNA Mini Kit | Thermo Fisher Scientific | Cat: 12183018A |
| Recombinant Human IL2 | R&D Systems | Cat: 202-IL-010 |
| Lipofectamine 3000 | Thermo Fisher Scientific | Cat: L3000001 |
| Mini-PROTEAN TGX StainFree 4-15% | BioRad | Cat: 456-8086 |
| Glycine | Fisher Scientific | Cat: BP381-1 |
| Tris Base | Fisher Scientific | Cat: BP152-1 |
| Sodium Chloride | Fisher Scientific | Cat: BP358-1 |
| Tween 20 | Fisher Scientific | Cat: BP337-500 |
| Non-fat Dry Milk | Lab Scientific | Cat: M0841 |
| MitoTEMPOL | Dr. Balaraman Kalyanaraman, University of Wisconsin | N/A |
| Rotenone | Sigma | Cat: R-8875 |
| Antimycin A | Sigma | Cat: A-8674 |
| Oligomycin | Sigma | Cat: O-4876 |
| Thenoyltrifluoroacetone (TTFA) | Sigma | Cat: T-9888 |
| Miliplex MAP Human cytokine/chemokine | EMD Millipore | Cat: HCYTOMAG-60K |
| **Viral Vectors** | | |
| CellLight Mitochondria-GFP, BacMam 2.0 | Thermo Fisher Scientific | Cat: C10508 |
| CellLight Tubulin-RFP, BacMam 2.0 | Thermo Fisher Scientific | Cat: C10503 |
| CellLight Lysosomes-RFP, BacMam 2.0 | Thermo Fisher Scientific | Cat: C10504 |
| CellLight Early Endosome-RFP, BacMam 2.0 | Thermo Fisher Scientific | Cat: C10587 |
| **Oligonucleotides** | | |
| Human IL17 A (forward primer)  5`-TACTACAACCGATCCACCTC-3` | Cytogence Genie  <https://genie.cytogence.com> | Primer ID: 301342 |
| Human IL17A (reverse primer)  5`-GAGTTCATGTGGTAGTCCAC-3` | Cytogence Genie  <https://genie.cytogence.com> | Primer ID: 301342 |
| Human IL4 (forward primer)  5`-ACGGACACAAGTGCGATATC-3` | Cytogence Genie  <https://genie.cytogence.com> | Primer ID: 302718 |
| Human IL4 (reverse primer)  5`-CTTCTCATGGTGGCTGTAGA-3` | Cytogence Genie  <https://genie.cytogence.com> | Primer ID: 302718 |
| Human DNM1L (forward primer)  5`-GCTCCAGGACGTCTTCAACA-3` | This Paper | N/A |
| Human DNM1L (reverse primer)  5`-TAGCACTGAGCTCTTTCCGC-3` | This Paper | N/A |
| Human ACTB (forward primer)  5`-TGCTATCCAGGCTGTGCTAT-3` | Hecker, et al., 2014 | N/A |
| Human ACTB (reverse primer)  5`-AGTCCATCACGATGCCAGT-3` | Bernard, et a., 2015 | N/A |
| GATA-3 (reverse primer)  5’-TCGGTTTCTGGTCTGGATGCCT-3’ | Eurofins Genomics | (Gene Accession#NM_001002295) |
| GATA-3 (forward primer)  5’ACCACAACCACACTCTGGAGGA-3’ | Eurofins Genomics | (Gene Accession#NM_001002295) |
| RORC (forward primer)  5’GAGGAAGTGACTGGCTACCAGA 3’ | Eurofins Genomics | (Gene Accession#NM_005012) |
| RORC (reverse primer)  5’GCACAATCTGGTCATTCTGGCAG 3’ | Eurofins Genomics | (Gene Accession#NM_005012) |
| RELA (forward primer)  5’ TGAACCGAAACTCTGGCAGCTG 3’ | Eurofins Genomics | (Gene Accession#NM_021975) |
| RELA (reverse primer)  5’ CATCAGCTTGCGAAAAGGAGCC 3’ | Eurofins Genomics | (Gene Accession#NM_021975) |
| GAPDH (forward primer)  5’ GTCTCCTCTGACTTCAACAGCG 3’ | Eurofins Genomics | (Gene Accession#NM_002046) |
| GAPDH (reverse primer)  5’ACCACCCTGTTGCTGTAGCCAA 3’ | Eurofins Genomics | (Gene Accession#NM_002046) |
| ON-TARGETplus SMARTpool Human DNM1L | Dharmacon | Cat: L-012092-00-0005 |
| **Software** | | |
| GraphPad Prism v5.04 | <https://www.graphpad.com/> | N/A |
| FlowJo X | <https://www.flowjo.com/> | N/A |
| IDEAS 6.2 | Luminex Corporation | N/A |
| R 3.5.2 64-bit | <https://www.r-project.org/> | N/A |
| Metaboanalyst 3.0 | <https://www.metaboanalyst.ca/> | N/A |
| FIJI ImageJ 1.52n | <https://fiji.sc/> | N/A |
| Nikon NIS-Element | Nikon | N/A |
