## Supplementary Figures for "Small Extracellular Vesicle Signaling and Mitochondrial Transfer Reprograms T Helper Cell Function in Human Asthma"

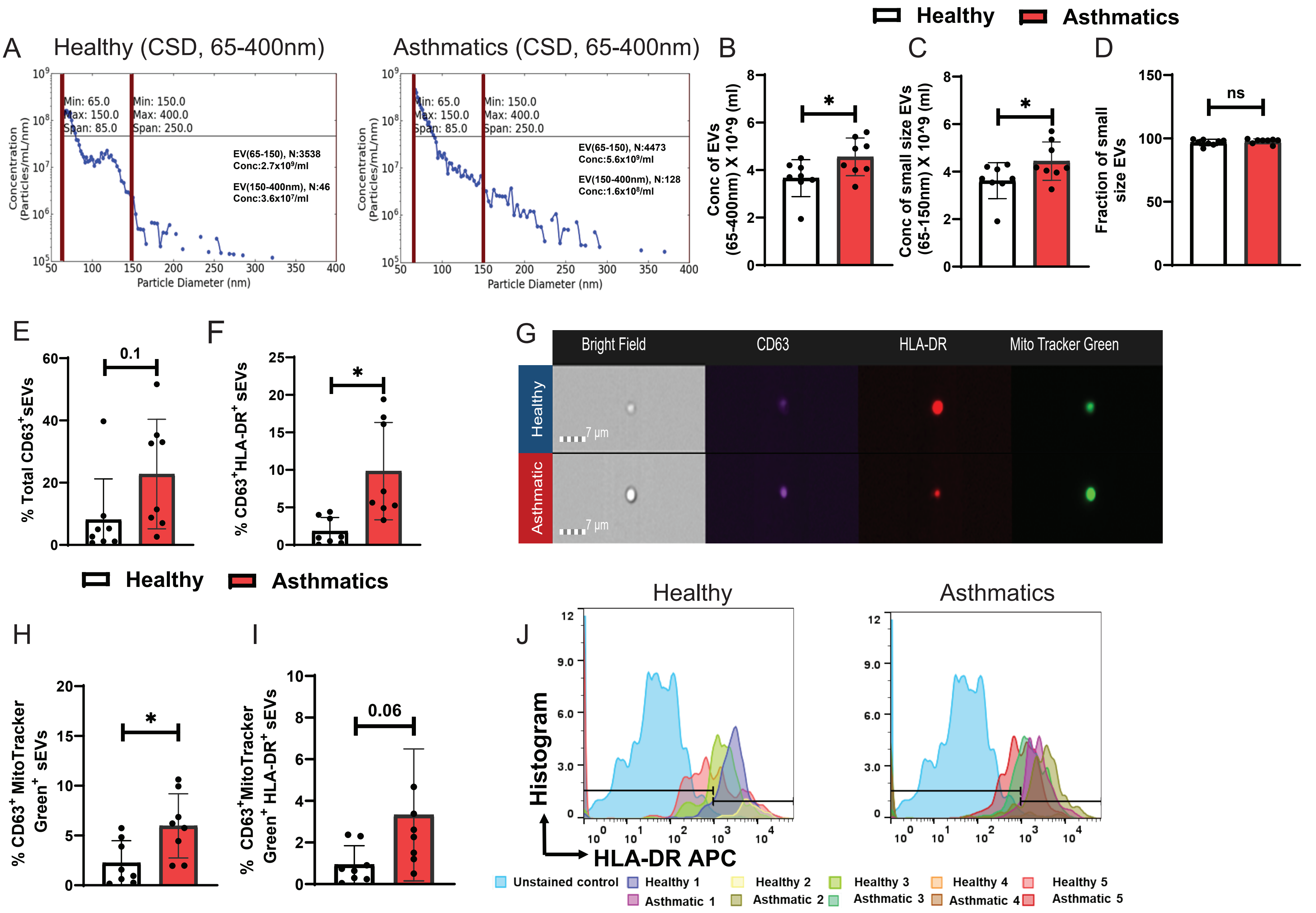
Healthy
Asthmatics
A
Healthy (CSD, 65-400nm)
Asthmatics (CSD, 65-400nm)
Min: 65.0
Max: 150.0
Span: 85.0
Min: 150.0
Max: 400.0
Span: 250.0
EV(65-150), N:3538
Conc:2.7x10⁹/ml
EV(150-400nm), N:46
Conc:3.6x10⁷/ml
EV(65-150), N:4473
Conc:5.6x10⁹/ml
EV(150-400nm), N:128
Conc:1.6x10⁸/ml
Concentration (Particles/mL/nm)
Particle Diameter (nm)
B
Conc of EVs (65-400nm) X 10^9 (ml)
C
Conc of small size EVs (65-150nm) X 10^9 (ml)
D
Fraction of small size EVs
ns
E
% Total CD63+sEVs
0.1
F
% CD63+HLA-DR+ sEVs
G
Bright Field
CD63
HLA-DR
Mito Tracker Green
Healthy
Asthmatic
7 µm
H
% CD63+ MitoTracker Green+ sEVs
I
% CD63+MitoTracker Green+ HLA-DR+ sEVs
0.06
J
Healthy
Asthmatics
Histogram
HLA-DR APC
Unstained control
Healthy 1
Healthy 2
Healthy 3
Healthy 4
Healthy 5
Asthmatic 1
Asthmatic 2
Asthmatic 3
Asthmatic 4
Asthmatic 5

**Supplementary Figure 1** – BALF CD63$^+$ sEVs and MitoTrackerGreen$^+$ sEVs within these sEVs have Class II expression. BALF sEVs isolated from Healthy controls and Asthmatics were assessed by Spectradyne's nCS1$^{TM}$ Particle Analyzer that delivers accurate nanoparticle size and concentration. (A) Representative acquired CSD image of quantification of BALF sEVs, Red Box highlights the sEV gate that is quantitated in B-D. (B) Concentration of all BALF EVs comparing Healthy controls and Asthmatics. (C) Concentration of BALF sEVs comparing Healthy controls and Asthmatics. (D) Percent sEVs of total BALF EVs comparing Healthy controls and Asthmatics. (E) Percent total CD63$^+$ sEVs isolated from BALF comparing Healthy Controls and Asthmatics. (F) Percent of CD63$^+$ HLA-DR$^+$ sEVs from BALF comparing Healthy controls and Asthmatics. (G) Representative image strips from ImageStream analyses showing HLA-DR$^+$ and MitoTracker Green$^+$ BALF sEVs in Healthy controls and Asthmatics. (H) Quantitation of % CD63$^+$ MitoTracker Green$^+$ BALF sEVs by ImageStream. (I) Quantitation of % CD63$^+$MitoTrackerGreen$^+$HLA-DR$^+$ BALF sEVs by ImageStream. (J) Overlaid Histograms of CD63$^+$MitoTracker Green$^+$ gated BALF sEVs showing HLA-DR expression in Healthy controls and Asthmatics. BALF EVs were isolated from n=8 study subjects/group and represented as each data point in B-F and H-I. Data shown in J is n=5 for Healthy controls and Asthmatics. Mann Whitney T test for comparison between Controls and Asthma groups $^*p<0.05$.

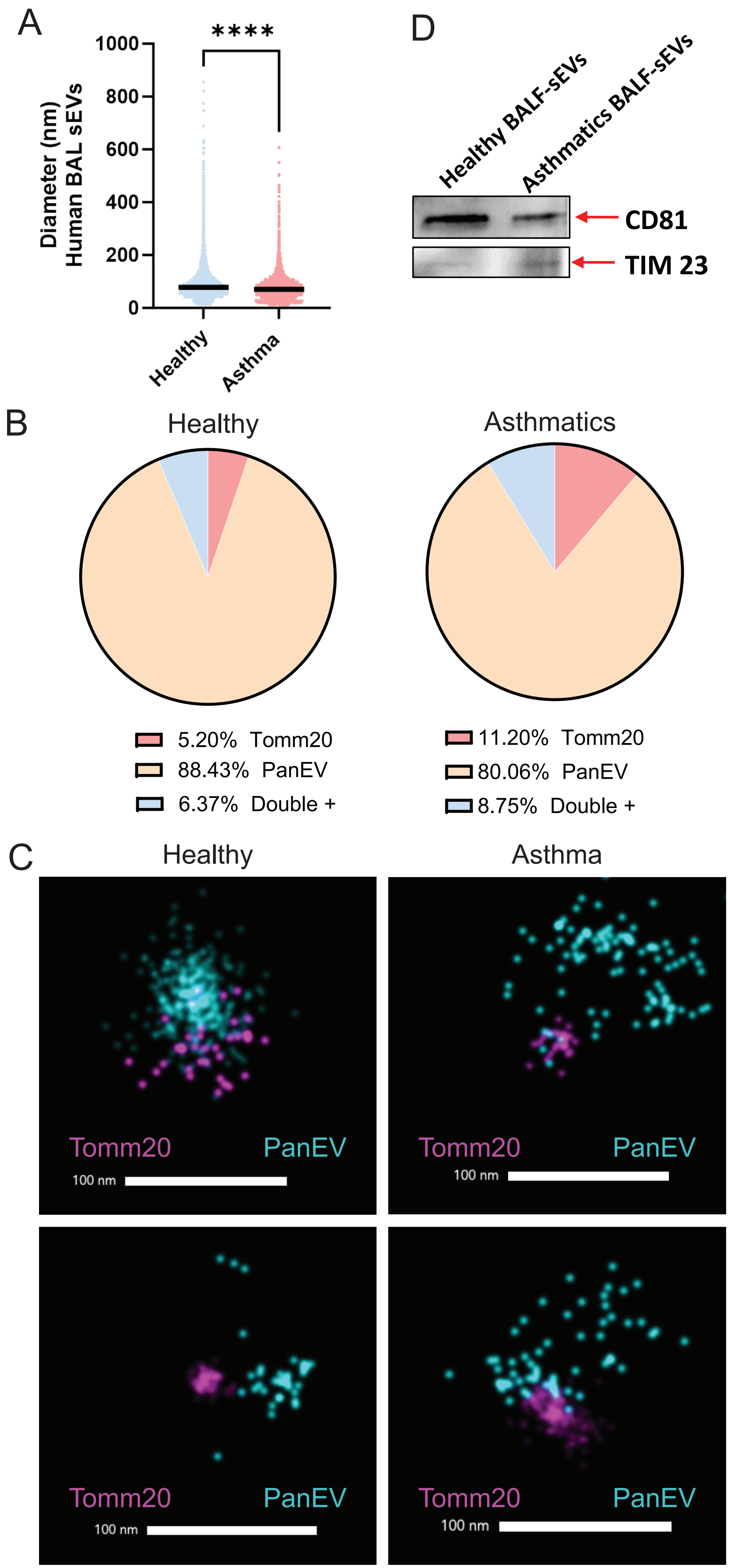
A
****
Diameter (nm)
Human BAL sEVs
1000
800
600
400
200
0
Healthy
Asthma
D
Healthy BALF-sEVs
Asthmatics BALF-sEVs
CD81
TIM 23
B
Healthy
Asthmatics
5.20% Tomm20
88.43% PanEV
6.37% Double +
11.20% Tomm20
80.06% PanEV
8.75% Double +
C
Healthy
Asthma
Tomm20
PanEV
100 nm

**Supplementary Figure 2** – Human BALF- sEVs express PanEV markers and Tomm20. Purified BALF-sEVs from Healthy human controls and Asthmatics were captured and stained using the Oxford Nanoimaging (ONi) EV Profiler Kit 2. Imaging was completed using the AutoEV function on the ONi Nanoimager and utilized direct stochastic optical reconstruction microscopy for high resolution images. All analyses was completed in the CODI software. (A) Quantitation of diameter of BALF-sEVs from Healthy controls and Asthmatics. (B) Pie charts showing % of sEVs expressing Tomm20 and Pan EV markers in samples from A, determined by Oni image analyses of 10000 events collected for n=3 replicates from pooled BALF-sEVs isolated from n=6/group of Healthy controls and Asthmatics. (C) Representative high resolution images from nanoimager showing sEVs expressing Tomm 20 and PanEV markers in BALF-sEVs from Heathy controls and Asthmatics. Scale bar=100 nm. Kolmogorov Smirnov test for comparison between Healthy controls and Asthmatics, ****p<0.001. (D) Western Blot Analyses of BALF sEVs from Healthy and Asthmatics isolated as above, lysed and electrophoresed on SDS PAGE and probed with anti-CD81 and anti-TIM23 antibody. Western Blot shows expression of CD81 and TIM 23.

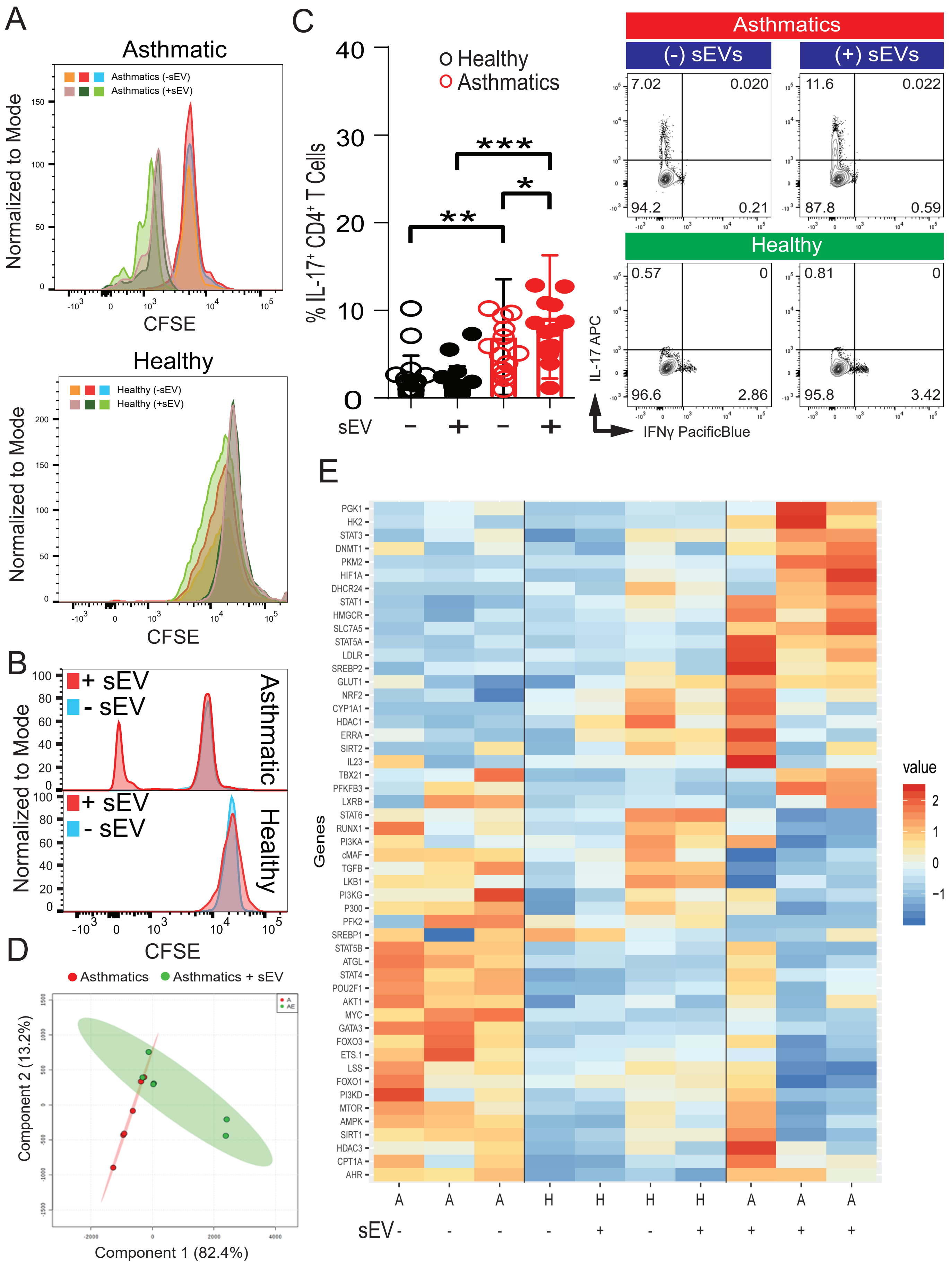
A
Asthmatic
Asthmatics (-sEV)
Asthmatics (+sEV)
Normalized to Mode
CFSE
Healthy
Healthy (-sEV)
Healthy (+sEV)
B
+ sEV
- sEV
Asthmatic
Healthy
C
Healthy
Asthmatics
% IL-17+ CD4+ T Cells
sEV
Asthmatics
(-) sEVs
(+) sEVs
7.02
0.020
94.2
0.21
11.6
0.022
87.8
0.59
Healthy
0.57
0
96.6
2.86
0.81
0
95.8
3.42
IL-17 APC
IFNγ PacificBlue
D
Asthmatics
Asthmatics + sEV
Component 2 (13.2%)
Component 1 (82.4%)
E
Genes
value
PGK1
HK2
STAT3
DNMT1
PKM2
HIF1A
DHCR24
STAT1
HMGCR
SLC7A5
STAT5A
LDLR
SREBP2
GLUT1
NRF2
CYP1A1
HDAC1
ERRA
SIRT2
IL23
TBX21
PFKFB3
LXRB
STAT6
RUNX1
PI3KA
cMAF
TGFB
LKB1
PI3KG
P300
PFK2
SREBP1
STAT5B
ATGL
STAT4
POU2F1
AKT1
MYC
GATA3
FOXO3
ETS.1
LSS
FOXO1
PI3KD
MTOR
AMPK
SIRT1
HDAC3
CPT1A
AHR
A A A H H H H A A A
sEV - - - - + - + + + +

**Supplementary Figure 3** – BALF sEVs from the airways of asthmatics promote proliferation and activation of CD4+ T cells. (A-D) Peripheral and airway $CD4^+$ T cells cultured with and without autologous BALF sEVs for 7 days in the presence of 50 IU/ml rhIL-2 in RPMI supplemented with 10% sEV depleted human AB serum. (A) Representative histogram of peripheral $CD4^+$ T cell CFSE dilution in the –sEV and +sEV groups following culture illustrating proliferation displayed for each group as an overlay. (B) Representative histogram of airway $CD4^+$ T cell CFSE dilution illustrating proliferation displayed for each group as an overlay. (C) sEVs from the BALF fluid were co-cultured with autologous peripheral human $CD4^+$ T cells and percent Th17 subsets analyzed. Left, percent of Th17 cells (assessed by IL-17 expression), and representative flow plots shown on the right. $CD4^+$ T cells were gated on IL-$4^{neg}$ and then plotted IL-17 versus IFNγ. Mann Whitney T test for comparison between healthy and asthmatics, Wilcoxon matched-pairs signed rank test for intergroup (no sEVs vs sEV), *<0.05, **<0.01, ***<0.001. (D) PCA analysis was conducted on gene expression data obtained from NanoString. PCA analysis was performed using R and MetaboAnalyst 3.0. (E) Heatmap of gene expression data obtained from NanoString. Hierarchical clustering and heatmap was generated using R (libraries used: ggplot2, reshape2, ggdendro, grid; A=Asthmatic, H=Healthy Normal).

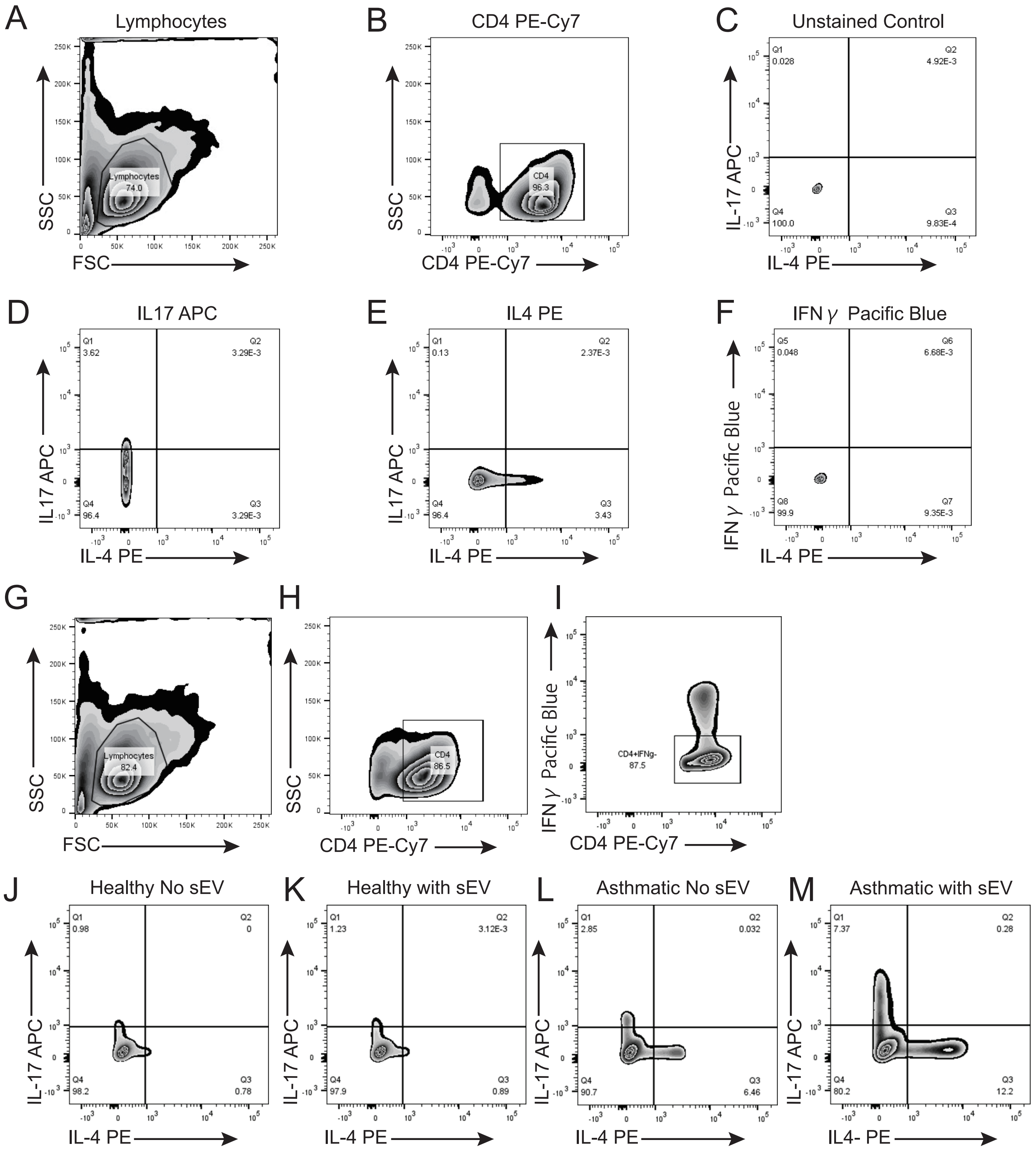

A
Lymphocytes
Lymphocytes 74.0
SSC
FSC
B
CD4 PE-Cy7
CD4 96.3
SSC
CD4 PE-Cy7
C
Unstained Control
Q1 0.028
Q2 4.92E-3
Q4 100.0
Q3 9.83E-4
IL-17 APC
IL-4 PE
D
IL17 APC
Q1 3.62
Q2 3.29E-3
Q4 96.4
Q3 3.29E-3
IL17 APC
IL-4 PE
E
IL4 PE
Q1 0.13
Q2 2.37E-3
Q4 96.4
Q3 3.43
IL17 APC
IL-4 PE
F
IFN γ Pacific Blue
Q5 0.048
Q6 6.68E-3
Q8 99.9
Q7 9.35E-3
IFN γ Pacific Blue
IL-4 PE
G
Lymphocytes 82.4
SSC
FSC
H
CD4 86.5
SSC
CD4 PE-Cy7
I
CD4+IFNg- 87.5
IFN γ Pacific Blue
CD4 PE-Cy7
J
Healthy No sEV
Q1 0.98
Q2 0
Q4 98.2
Q3 0.78
IL-17 APC
IL-4 PE
K
Healthy with sEV
Q1 1.23
Q2 3.12E-3
Q4 97.9
Q3 0.89
IL-17 APC
IL-4 PE
L
Asthmatic No sEV
Q1 2.85
Q2 0.032
Q4 90.7
Q3 6.46
IL-17 APC
IL-4 PE
M
Asthmatic with sEV
Q1 7.37
Q2 0.28
Q4 80.2
Q3 12.2
IL-17 APC
IL4- PE

**Supplementary Figure 4** – Gating Strategy and Flow Cytometry Controls for T cells from Healthy controls and Asthmatics co-cultured with autologous BALF-EVs and polarized to Th17 and Th2 subsets. (A-B) Cells were gated initially for Lymphocytes (A) followed by CD4 gate (B), (C-F) Unstained and Single color controls,(G-M) Gating strategy for sEV-co-cultured and polarized Th2 and Th17 cells from Healthy controls and Asthmatics as described above, (G-H) Lymphocyte gate (G) followed by CD4 gate (H), (I) $CD4^+$ cells were gated for $IFN\gamma^{neg}$ cells, (J-M) Representative flow plots for gates for Th2 and Th17 cells polarized in co-cultures of autologous peripheral T cells in presence or absence of BALF sEVs from Healthy controls and Asthmatics.

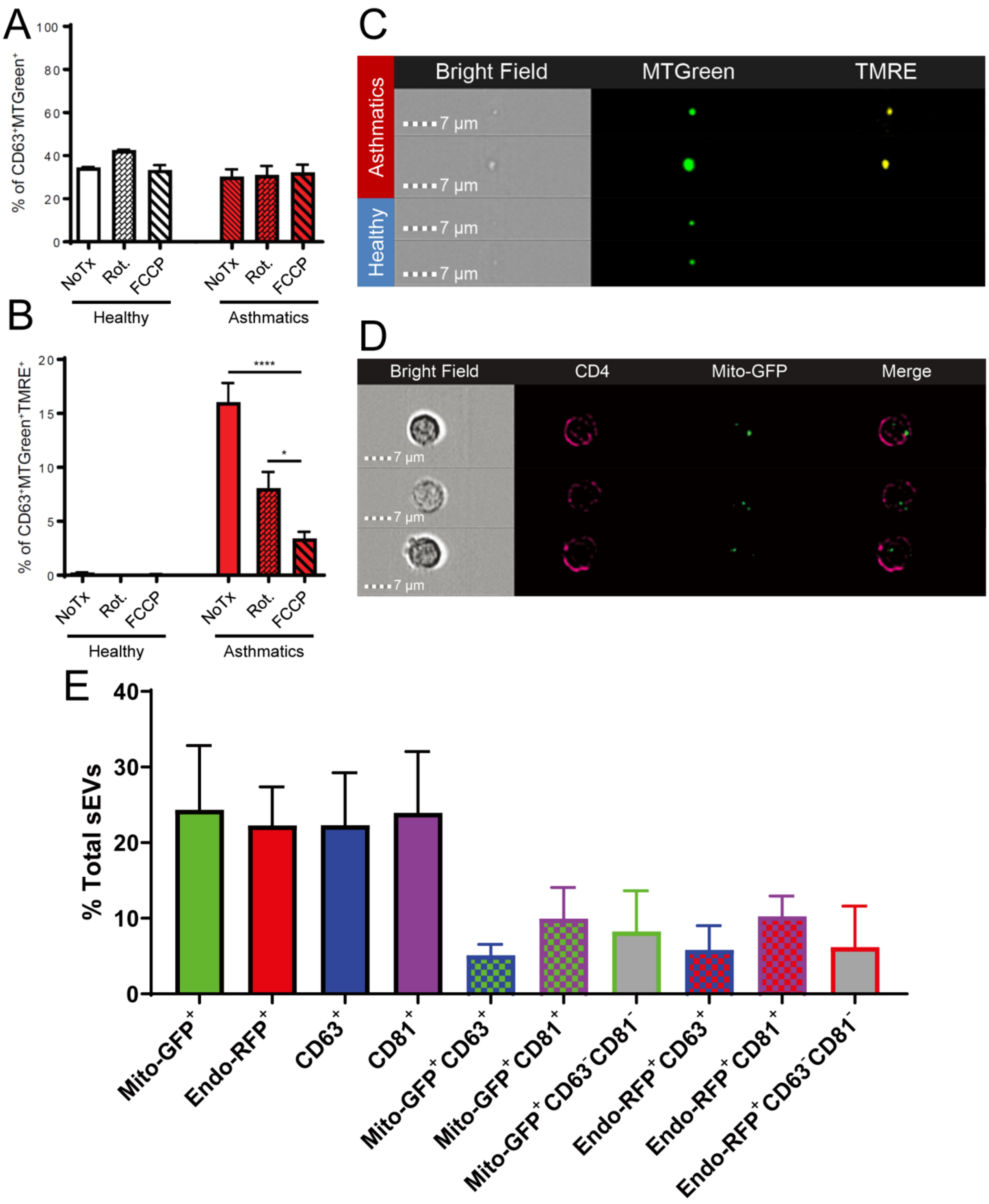


**Supplementary Figure 5** – Mitochondrial membrane potential was observed only in MDRC-derived sEVs from asthmatics. (A-C) Airway MDRCs were labeled with MitoTracker Green and sEVs purified from the conditioned media 48 hours later. Purified MDRC sEVs were labeled with TMRE and analyzed on the ImageStream flow cytometer. (A) The % of MitoTracker Green MDRC sEVs that are $CD63^{+}$. (B) The % of $TMRE^{+}$ MDRC sEVs that are also positive for CD63 and MitoTracker Green. (C) Representative image strip of MDRC-derived sEVs analyzed by ImageStream. (D-E) MDRC-derived sEVs contain mitochondria and are internalized by autologous peripheral $CD4^{+}$ T cells. (D) Representative image strip of peripheral $CD4^{+}$ T cells internalizing Mito-$GFP^{+}$ sEVs generated from MDRCs. (E) Characterization of sEVs from MDRCs that were transduced with CellLight Mito-GFP, CellLight Endo-RFP. Transduced MDRCs were cultured for 48 hours before isolation of sEVs from the conditioned media. MDRC sEVs were probed with anti-CD63 and anti-CD81 and characterized on ImageStream flow cytometry.

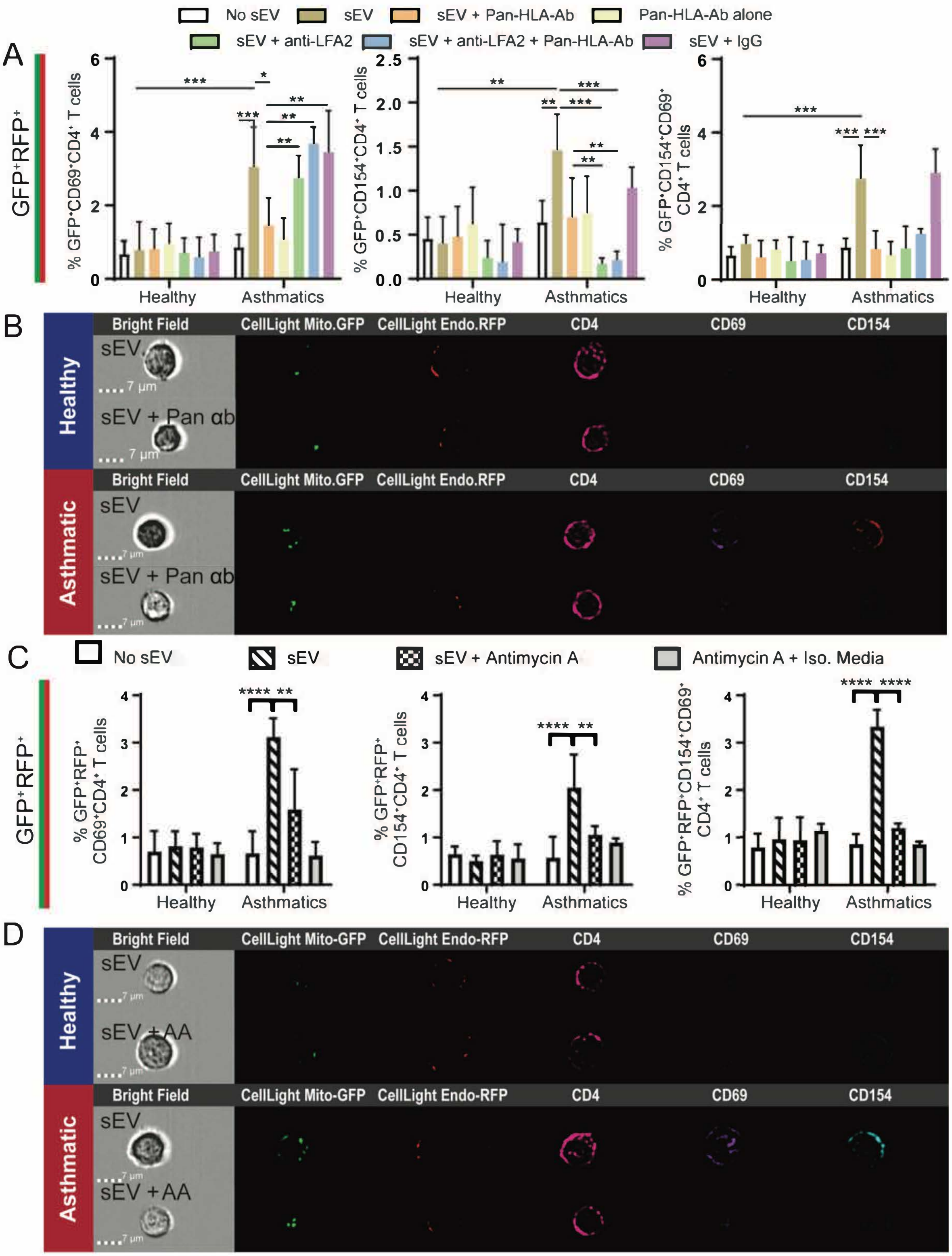


**Supplementary Figure 6** – Blockade of class II molecules using a pan-HLA antibody (HLA-DR/DP/DQ) results in loss of MDRC sEV mediated activation of autologous peripheral CD4$^+$ T cells. (A) Graphs illustrating the percentage of CD69$^+$, CD154$^+$, or CD69$^+$CD154$^+$ T cells that internalized Mito-GFP$^+$ Endo-RFP$^+$ MDRC sEVs. Mann Whitney T, *<0.05, **<0.01, ***<0.001, ****<0.0001. (B) Representative image strips from ImageStream illustrating that Mito-GFP$^+$ Endo-RFP$^+$ MDRC sEVs activate T cells in asthmatics, while treatment with pan-HLA-Ab abrogates this activation by MDRC sEVs. (C-D) Inhibition of complex III by antimycin A in MDRC sEVs diminishes autologous peripheral CD4$^+$ T cell activation in asthmatics. Results for CD4$^+$ T cells that have internalized both Mito-GFP$^+$ MDRC sEVs and Mito-RFP$^+$ MDRC sEVs. (C) Graphs illustrating the percentage of CD69$^+$, CD154$^+$, or CD69$^+$CD154$^+$ T cells that internalized Mito-GFP$^+$ Endo-RFP$^+$ MDRC sEVs. Mann Whitney T, **<0.01, ***<0.001, ****<0.0001. (D) Representative image strips from ImageStream analysis illustrating that Mito-GFP$^+$ Endo-RFP$^+$ MDRC sEVs activate T cells in asthmatics, and antimycin blocks activation in asthmatics.

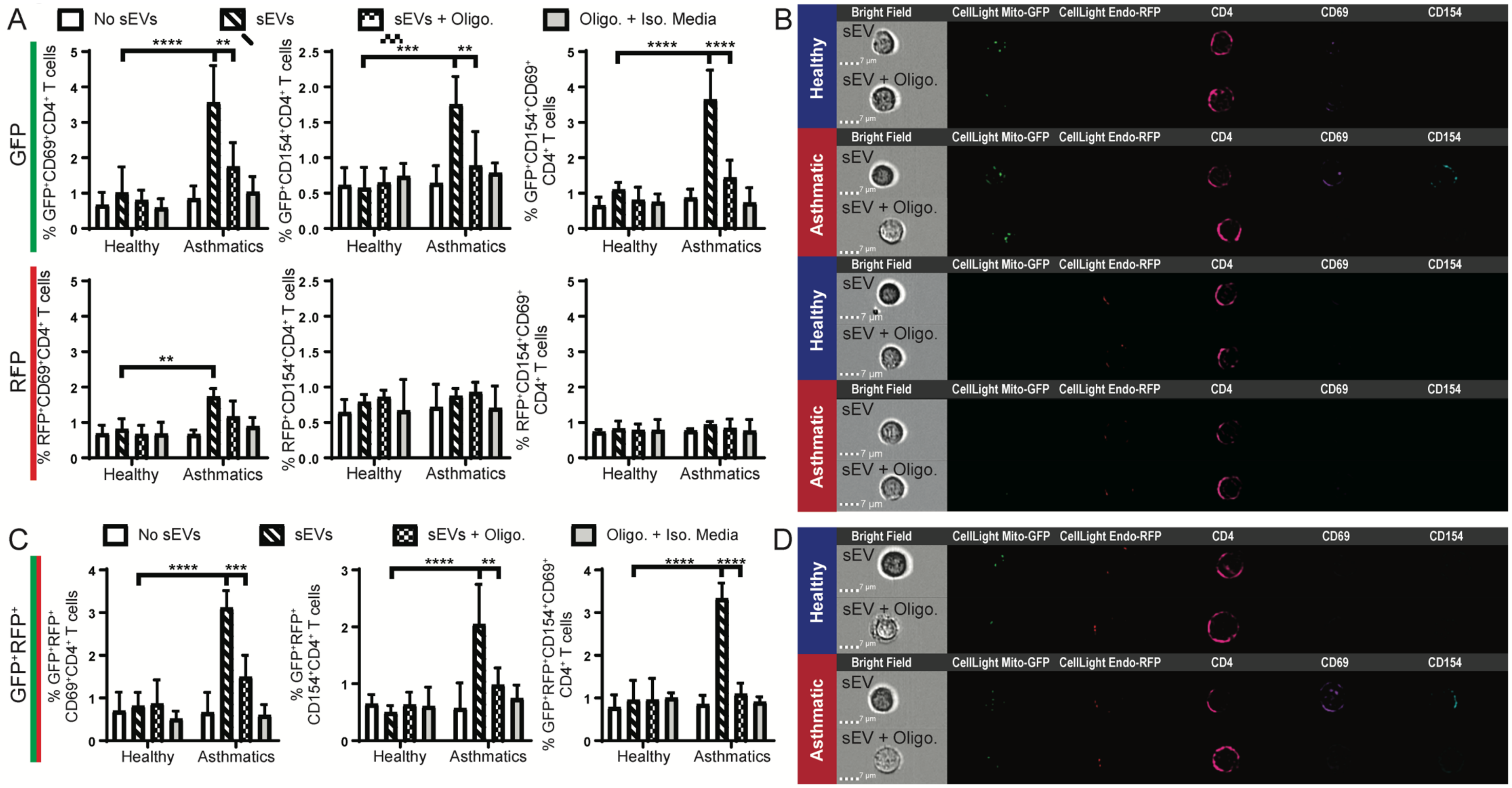


**Supplementary Figure 7** – Inhibition of complex V by oligomycin in MDRC sEVs diminishes autologous peripheral CD4⁺ T cell activation in asthmatics. MDRCs were transduced with CellLight Mito-GFP and CellLight Endo-RFP, and sEVs were purified from the supernatant 48 hours later. Purified MDRC sEVs were co-cultured with autologous peripheral CD4⁺ T cells for 24 hours in the presence of rhIL-2 (50 IU/ml) in a ration of 1:10 T cells:sEVs. MDRC sEVs were pre-treated with oligomycin (10 µM) overnight or untreated prior to co-culture with T cells. sEVs were washed and re-purified using the Invitrogen Total Exosome Isolation kit. ImageStream flow cytometry was used to assess early activation (CD69) and antigen-specific activation (CD154). (A) Graphs illustrating the percentage of CD69⁺, CD154⁺, or CD69⁺CD154⁺ T cells that internalized either Mito-GFP⁺ MDRC sEVs (top row) or Endo-RFP⁺ MDRC sEVs (bottom row). Mann Whitney T, **<0.01, ***<0.001, ****<0.0001. (B) Representative image strips from ImageStream analysis illustrating that Mito-GFP⁺ MDRC sEVs activate T cells in asthmatics, and oligomycin blocks activation in asthmatics. (C) Graphs illustrating the percentage of CD69⁺, CD154⁺, or CD69⁺CD154⁺ T cells that internalized either Mito-GFP⁺ Endo-RFP⁺ MDRC sEVs. Mann Whitney T, **<0.01, ***<0.001, ****<0.0001. (D) Representative image strips from ImageStream analysis illustrating that Mito-GFP + Endo-RFP⁺ MDRC sEVs activate T cells in asthmatics, and oligomycin blocks activation in asthmatics.

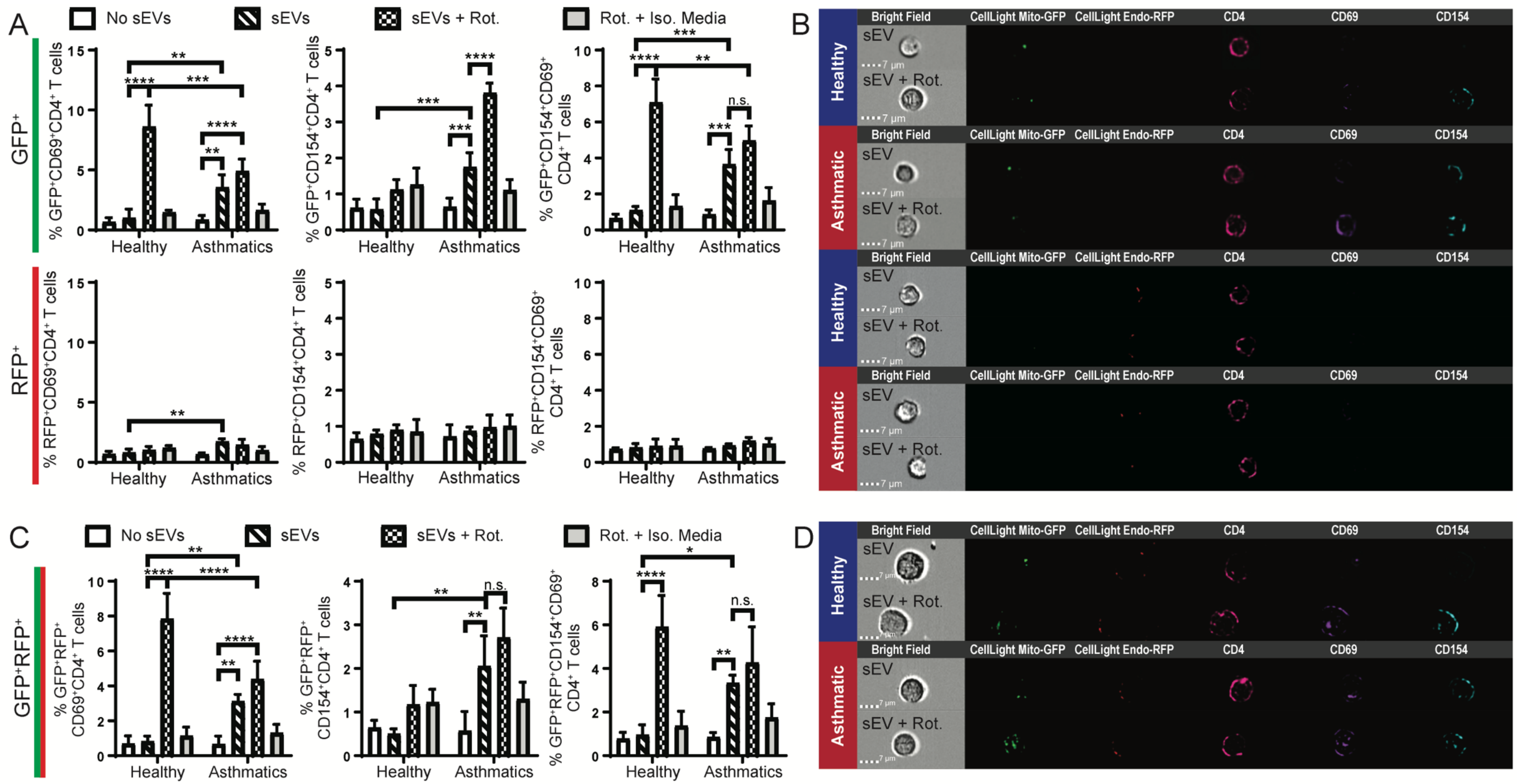


**Supplementary Figure 8** – Inhibition of complex I by rotenone in sEVs enhances autologous peripheral CD4$^+$ T cell activation in both healthy and asthmatics. MDRCs were transduced with CellLight Mito-GFP and CellLight Endo-RFP, and sEVs were purified from the supernatant 48 hours later. Purified MDRC sEVs were co-cultured with autologous peripheral CD4$^+$ T cells for 24 hours in the presence of rhIL-2 (50 IU/ml) in a ration of 1:10 T cells:sEVs. MDRC sEVs were pre-treated with rotenone (10 μM) overnight or untreated prior to co-culture with T cells. sEVs were washed and re-purified using the Invitrogen Total Exosome Isolation kit. ImageStream flow cytometry was used to assess early activation (CD69) and antigen-specific activation (CD154). (A) Graphs illustrating the percentage of CD69$^+$, CD154$^+$, or CD69$^+$CD154$^+$ T cells that internalized either Mito-GFP$^+$ MDRC sEVs (top row) or Endo-RFP$^+$ MDRC sEVs (bottom row). Mann Whitney T, **<0.01, ***<0.001, ****<0.0001. (B) Representative image strips from ImageStream analysis illustrating that Mito-GFP$^+$ MDRC sEVs activate T cells in asthmatics, and rotenone promotes activation in T cells from healthy and asthmatic subjects. (C) Graphs illustrating the percentage of CD69$^+$, CD154$^+$, or CD69$^+$CD154$^+$ T cells that internalized Mito-GFP$^+$ Endo-RFP$^+$ MDRC sEVs. Mann Whitney T, **<0.01, ***<0.001, ****<0.0001. (D) Representative image strips from ImageStream analysis illustrating that Mito-GFP$^+$ Endo-RFP$^+$ MDRC sEVs activate T cells in asthmatics, and rotenone promotes activation in healthy and asthmatic T cells.

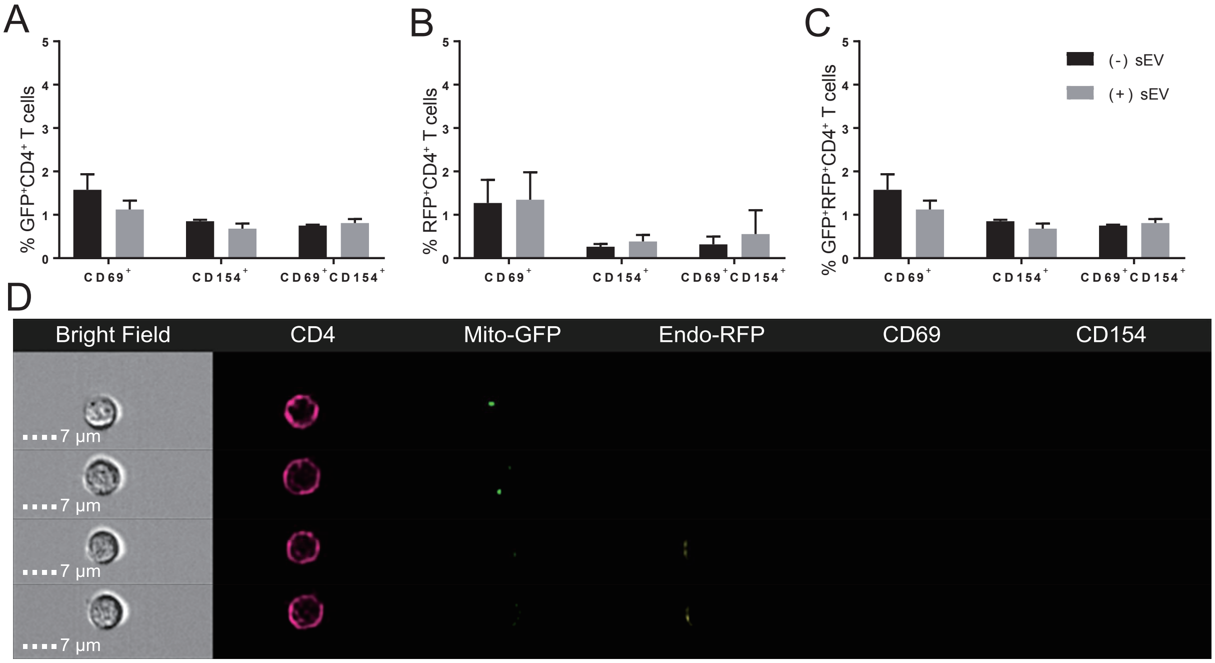


**Supplementary Figure 9** – sEVs derived from CD45 negative cells did not activate autologous peripheral CD4⁺ T cells. CD45$^{neg}$ non-immune cells were transduced with CellLight Mito-GFP and CellLight Endo-RFP. Transduced cells were cultured for 48 hours and then sEVs isolated from the conditioned media. Purified CD45$^{neg}$ sEVs were co-cultured with autologous peripheral CD4⁺ T cells for 24 hours in the presence of rhIL-2 (50 IU/ml) in a ration of 1:10 T cells:sEVs. ImageStream flow cytometry was used to assess early activation (CD69) and antigen-specific activation (CD154). (A-C) Graphs illustrating the percentage of CD69⁺, CD154⁺, or CD69⁺CD154⁺ T cells that internalized either (A) Mito-GFP⁺ sEVs, (B) Endo-RFP⁺ sEVs, or (C) both Mito-GFP⁺ and Endo-RFP⁺ sEVs. (D) Representative image strips from ImageStream analysis illustrating that activation is not observed despite internalization of non-immune cell-derived Mito-GFP⁺ sEVs by CD4⁺ T cells.

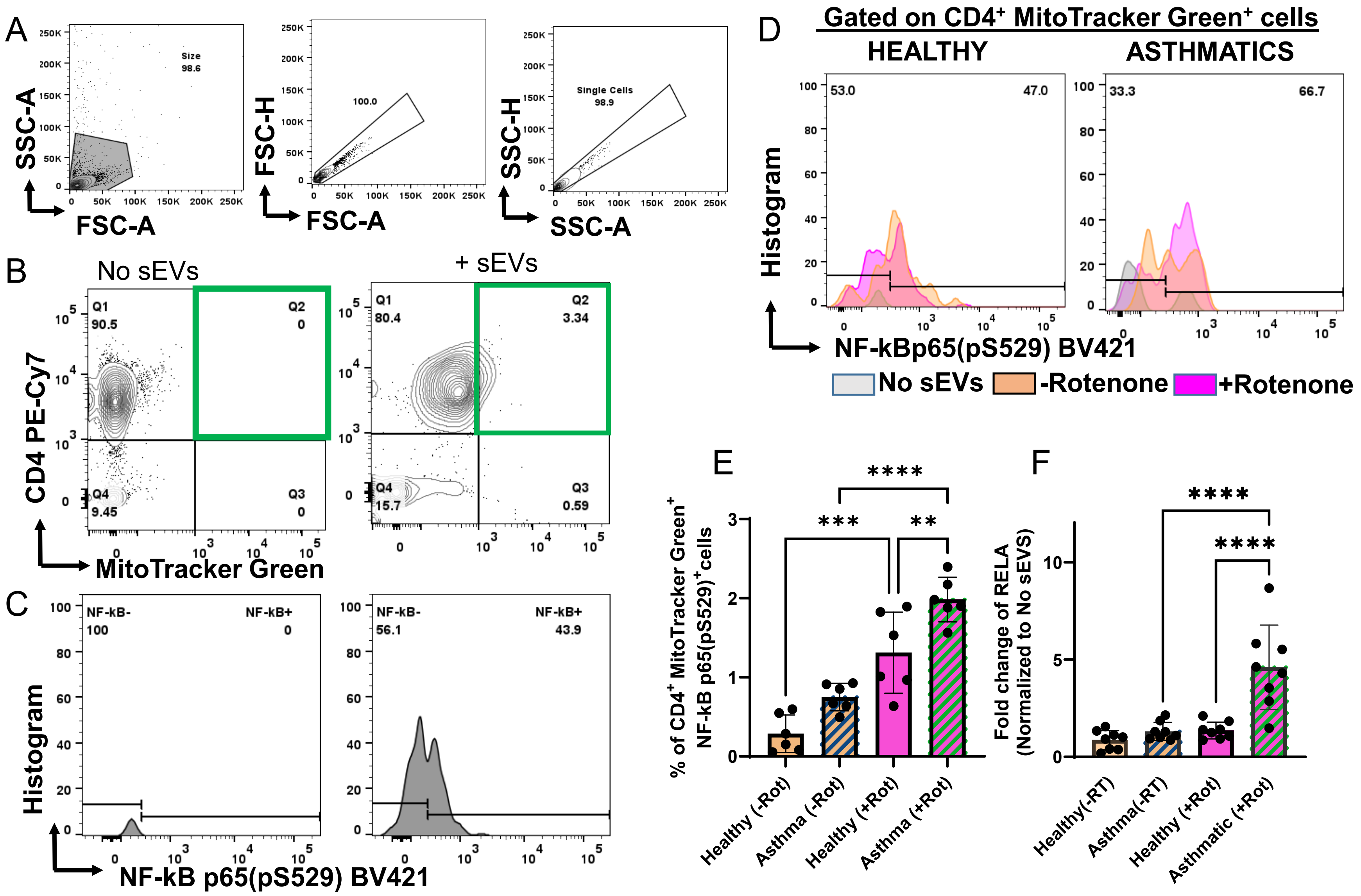


**Supplementary Figure 10** – NF-KB signaling in CD4 T cells is enhanced upon co-culture with Human BALF sEVs treated with rotenone, a Complex I inhibitor. Purified human BALF sEVs from healthy and asthmatic subjects were treated with 10 μM rotenone or vehicle control, washed, purified and labeled with MitoTrackerGreen. Autologous $CD4^{+}$ T cells were co-cultured with these sEVs at a 10 sEV: 1 T cell ratio and 24 hours later assessed for % NF-kB p65(pS529)$^{+}$ cells within the $CD4^{+}$MitoTrackerGreen$^{+}$ lymphocytes. (A) Gating strategy for ssc-fsc and singlets (B) Gating Strategy for $CD4^{+}$MitoTrackerGreen$^{+}$ lymphocytes (C) Gating strategy for NF-kB p65(pS529)$^{+}$ cells (D) Overlaid histogram of NF-kB p65(pS529)$^{+}$ cells in CD4 T cells co-cultured with no sEVs, vehicle treated sEVs and rotenone treated sEVs for 24 hours in healthy (n=6) and asthmatic (n=6) subjects. (E) Quantitation of % $CD4^{+}$MItoTrackerGreen$^{+}$ cells that are NF-kBp65(pS529)$^{+}$ from (D). (F) Fold change in RELA expression in peripheral $CD4^{+}$ T cells from healthy (n=8) and asthmatic (n=8) subjects following co-culture with vehicle treated and rotenone treated MitoTracker Green labeled sEVs. Data were normalized to GAPDH and no sEV controls. Gene Data analysis was performed using the 2-ΔΔCT method. One way ANOVA analyses with multiple comparison between treatments and study groups, **p<0.001, ***p=0.0001****p<0.001.

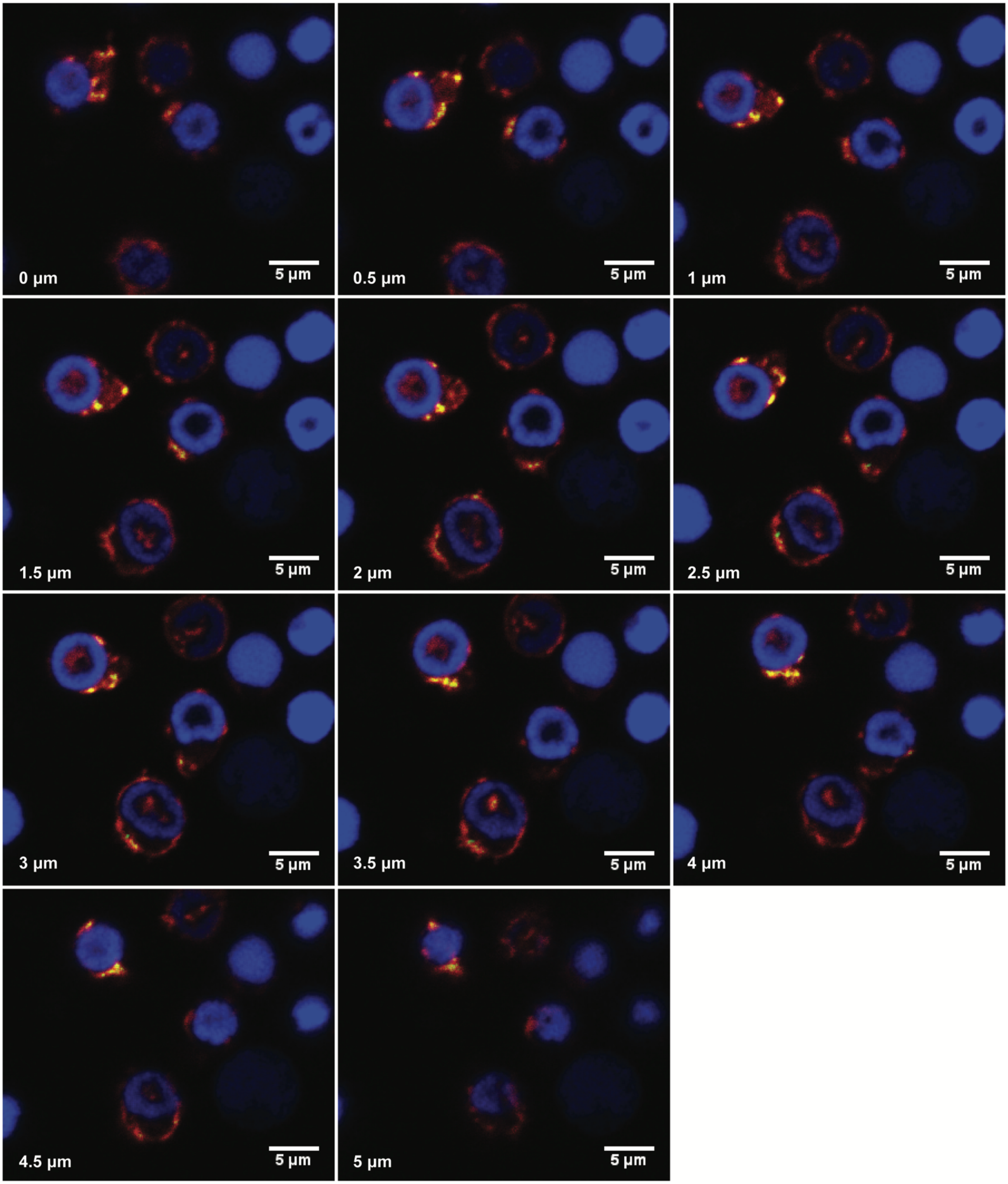


**Supplementary Figure 11** – Internalized Mito-GFP$^+$ MDRC sEVs co-localizes with cytosolic actin in T cells. Each panel represents a 0.5 µm slice in depth (z-stack). Confocal image illustrating internalization of MDRC sEVs inside the cytoplasm of the T cells. MDRCs were transduced with CellLight Mito-GFP and sEVs were purified from the supernatant 48 hours later. Mito-GFP$^+$ MDRC sEVs were co-cultured with T cells (1:10 T cell:sEV) for 24-hours in the ibidi µ-dish. Cells were fixed with 2% PFA and permeabilized with 0.5% triton-X in PBS. The fixed T cells were labeled with Phalloidine-Rhodamine and DAPI. Cells were imaged on the Nikon A1 confocal.

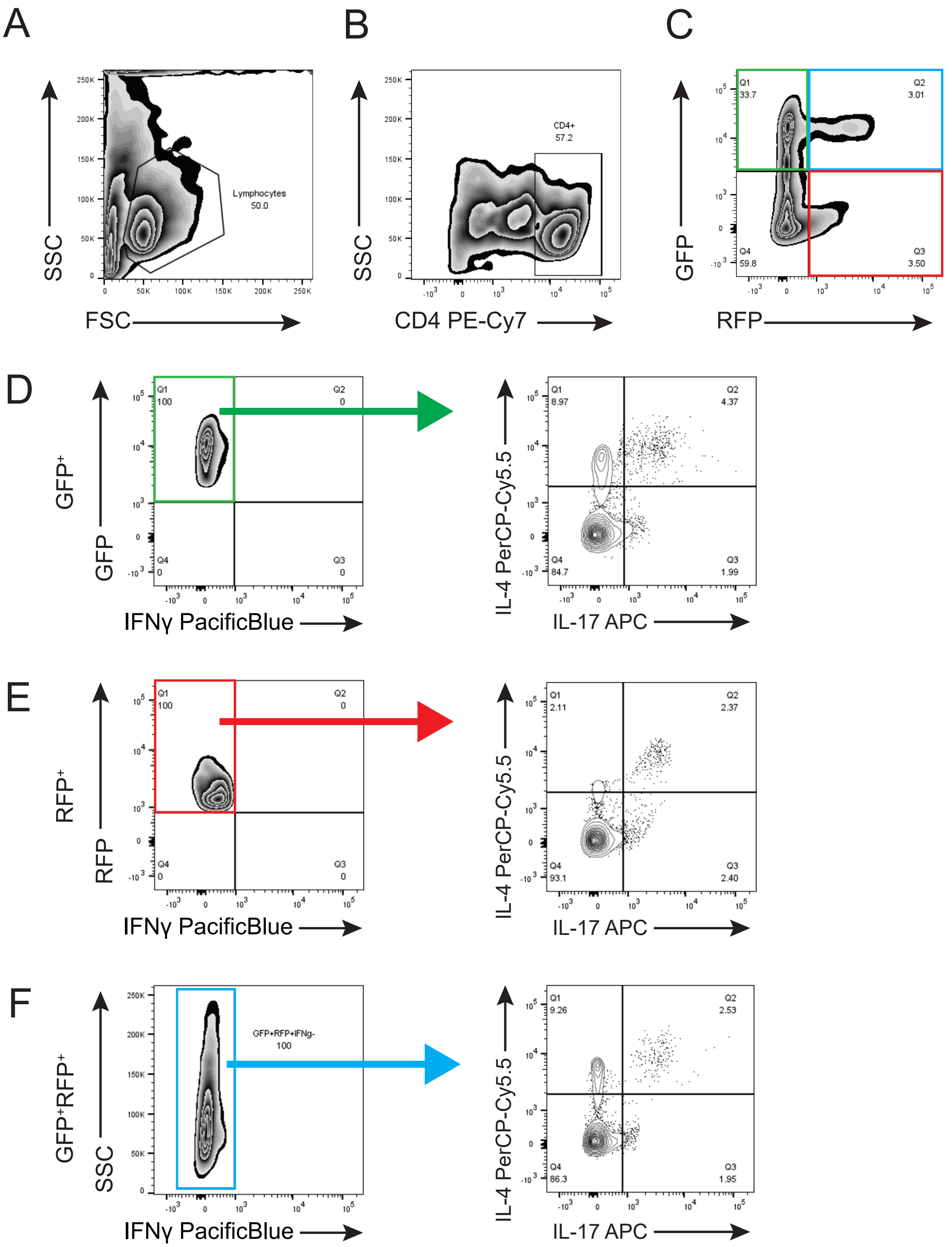


**Supplementary Figure 12** – Gating Strategy for T cells from Healthy controls and Asthmatics that internalize MDRC-derived Mito-GFP$^+$ sEVs and polarize to Th17 and Th2 subsets. Airway MDRCs were transduced with CellLight Mito-GFP and CellLight Endo-RFP, and sEVs were purified from the supernatant 48 hours later. Purified MDRC sEVs were co-cultured with autologous peripheral CD4$^+$T cells for 7 days in the presence of rhIL-2 (50 IU/ml) in a ratio of 1:10 T cells:sEVs. T helper subsets were assessed by flow cytometry 7 days later by staining intracellular for IL-4 (Th2) and IL-17(Th17) cytokines. (A-C) Cells were gated initially for CD4 followed by GFP$^+$, RFP$^+$ or GFP$^+$RFP$^+$ gate. (D) The IFNγ$^{neg}$ were gated within the GFP$^+$ gate and shown as IL-4$^+$ cells or IL-4$^{neg}$IL-17$^+$ cells, or IL-4$^+$IL-17$^+$. (E) The IFNγ$^{neg}$ were gated within the RFP$^+$ gate and shown as IL-4$^+$ cells or IL-4$^{neg}$IL-17$^+$ cells, or IL-4$^+$IL-17$^+$. (F) The IFNγ$^{neg}$ were gated based on SSC within the GFP$^+$RFP$^+$ gate and shown as IL-4$^+$ cells or IL-4$^{neg}$IL-17$^+$ cells or IL-4$^+$IL-17$^+$cells.

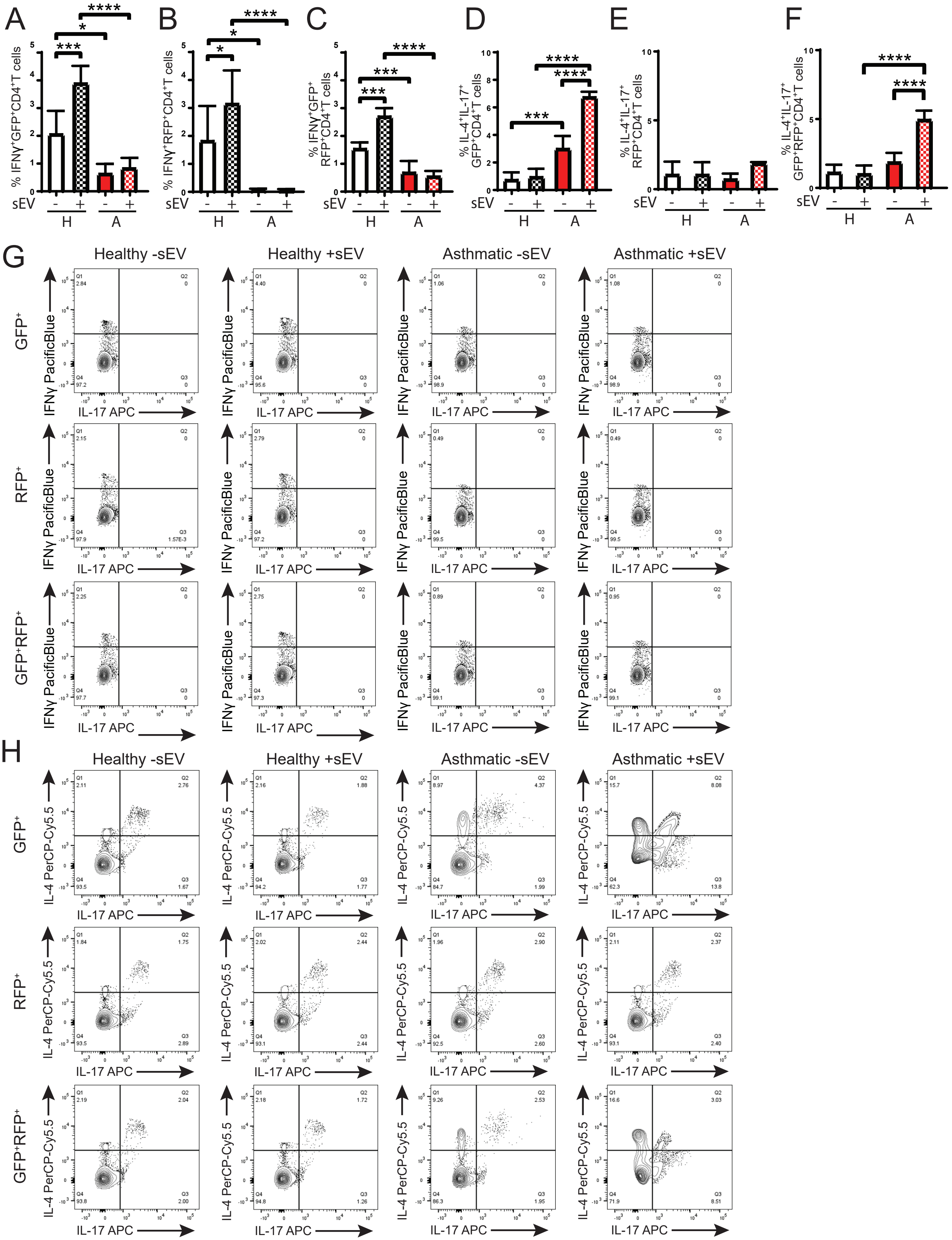

A
B
C
D
E
F
% IFNγ+GFP+CD4+T cells
% IFNγ+RFP+CD4+T cells
% IFNγ+GFP+ RFP+CD4+T cells
% IL-4+IL-17+ GFP+CD4+T cells
% IL-4+IL-17+ RFP+CD4+T cells
% IL-4+IL-17+ GFP+RFP+CD4+T cells
sEV
H
A
G
H
Healthy -sEV
Healthy +sEV
Asthmatic -sEV
Asthmatic +sEV
GFP+
RFP+
GFP+RFP+
IFNγ PacificBlue
IL-4 PerCP-Cy5.5
IL-17 APC

**Supplementary Figure 13** – Autologous peripheral $CD4^+$ T cells that internalize MDRC-derived Mito-$GFP^+$ sEVs purified from healthy subjects polarize to Th1 hybrid subset but not in asthmatics. Autologous peripheral $CD4^+$ T cells in asthmatics that internalize MDRC-derived Mito-$GFP^+$ sEVs polarize also to Th2/Th17 hybrid subset. Purified MDRC sEVs were co-cultured with autologous peripheral $CD4^+$ T cells for 7 days in the presence of rhIL-2 (50 IU/ml) in a ration of 1:10 T cells:sEVs. T helper subsets were assessed by flow cytometry 7 days later by staining intracellular for IFN$\gamma$ (Th1), IL-4 (Th2) and IL-17 (Th17) cytokines. (A-C, & G) Cells were first gated on $GFP^+$, $RFP^+$, or $GFP^+RFP^+$, then IL-$4^{neg}$ and IL-$17^{neg}$for each. For the $GFP^+RFP^+$, SSC was used for y-axis and gated on IL-$4^{neg}$IL-$17^{neg}$ and the IFN$\gamma^+$ population plotted and quantitated. Mann Whitney T test for comparison between healthy and asthmatics, Wilcoxon matched-pairs signed rank test for intergroup (no sEVs vs sEV), *<0.05, **<0.01, ***<0.001, ****<0.0001. (D-F, & H) Co-culture was performed as in (A-C). T helper subsets were assessed by flow cytometry 7 days later by staining intracellular for IL-4 (Th2) and IL-17 (Th17) cytokines. Cells were first gated on $GFP^+$, $RFP^+$, or $GFP^+RFP^+$, then gated on IFN$\gamma^{neg}$ cells as in Figures 7 and S12, and then plotted IL-4 versus IL-17 and determined % IL-$4^+$IL-$17^+$cells. Mann Whitney T test for comparison between healthy and asthmatics, Wilcoxon matched-pairs signed rank test for intergroup (no sEVs vs sEV), *<0.05, **<0.01, ***<0.001, ****<0.0001.

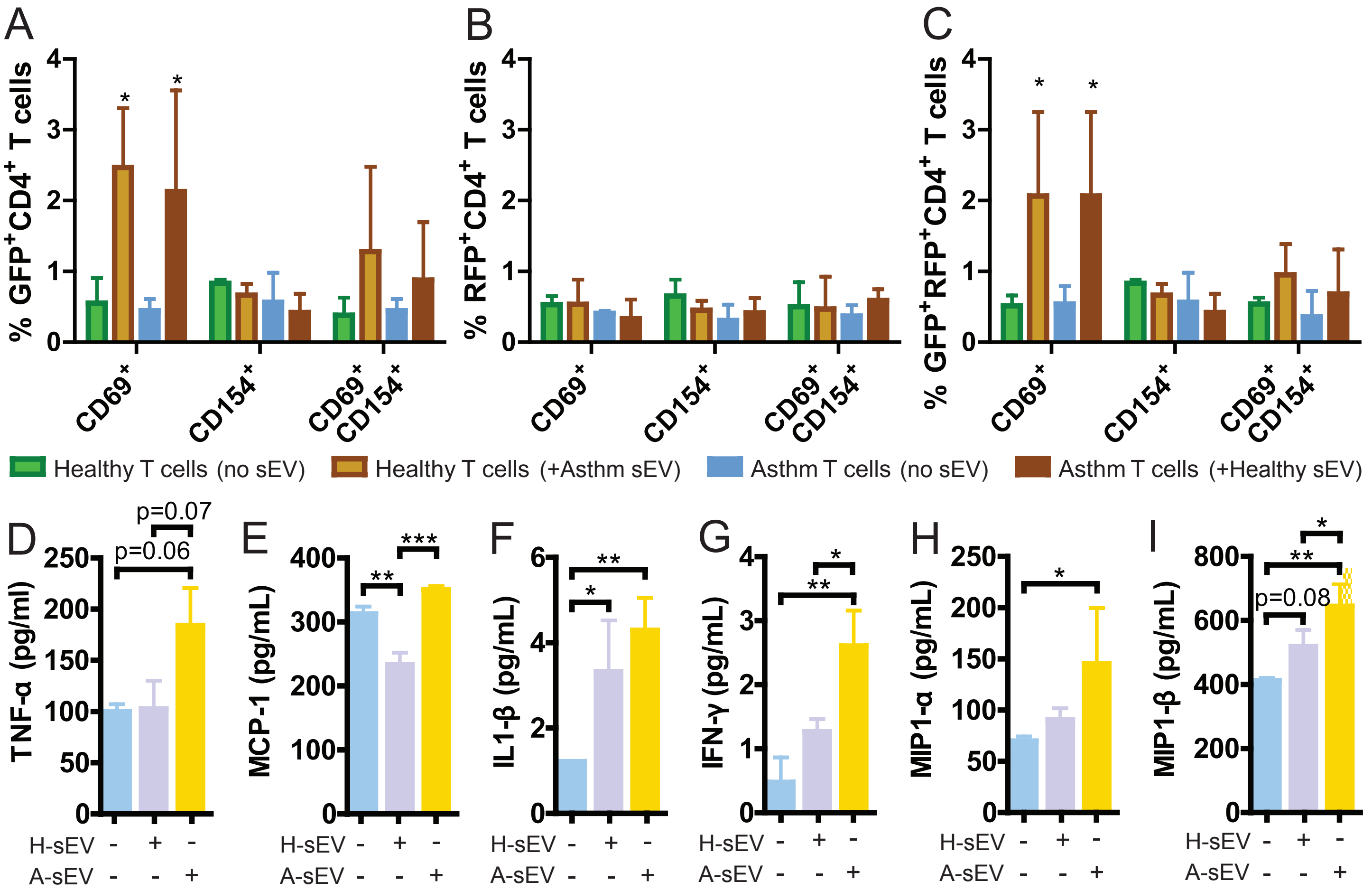


**Supplementary Figure 14** – (A-C) MDRCs were transduced with CellLight Mito-GFP and CellLight Endo-RFP, and sEVs were purified from the supernatant 48 hours later. Purified MDRC sEVs were then co-cultured with their allogenic T cells (i.e. healthy sEV on asthmatic T cells, and vice versa; at a 1:10 T cell:sEV) for 24 hours. Activation was assessed by CD69 and CD154 expression using the ImageStream cytometry (T-test comparison to no-sEV healthy control and no-sEV asthma control; *<0.05). (D-I) Airway sEVs were purified from BALF using a described ultracentrifugation method. Purified sEVs were cultured with THP-1 cells (1:10 cell:sEV) for 24 hours. Supernatants were harvested for BioRad Bioplex analysis of cytokines for (D) TNF-α, (E) MCP-1, (F) IL1- β, (G) IFNγ, (H) MIP1-α, and (I) MIP1-β. One-way ANOVA, *<0.05, **<0.01, ***<0.001.

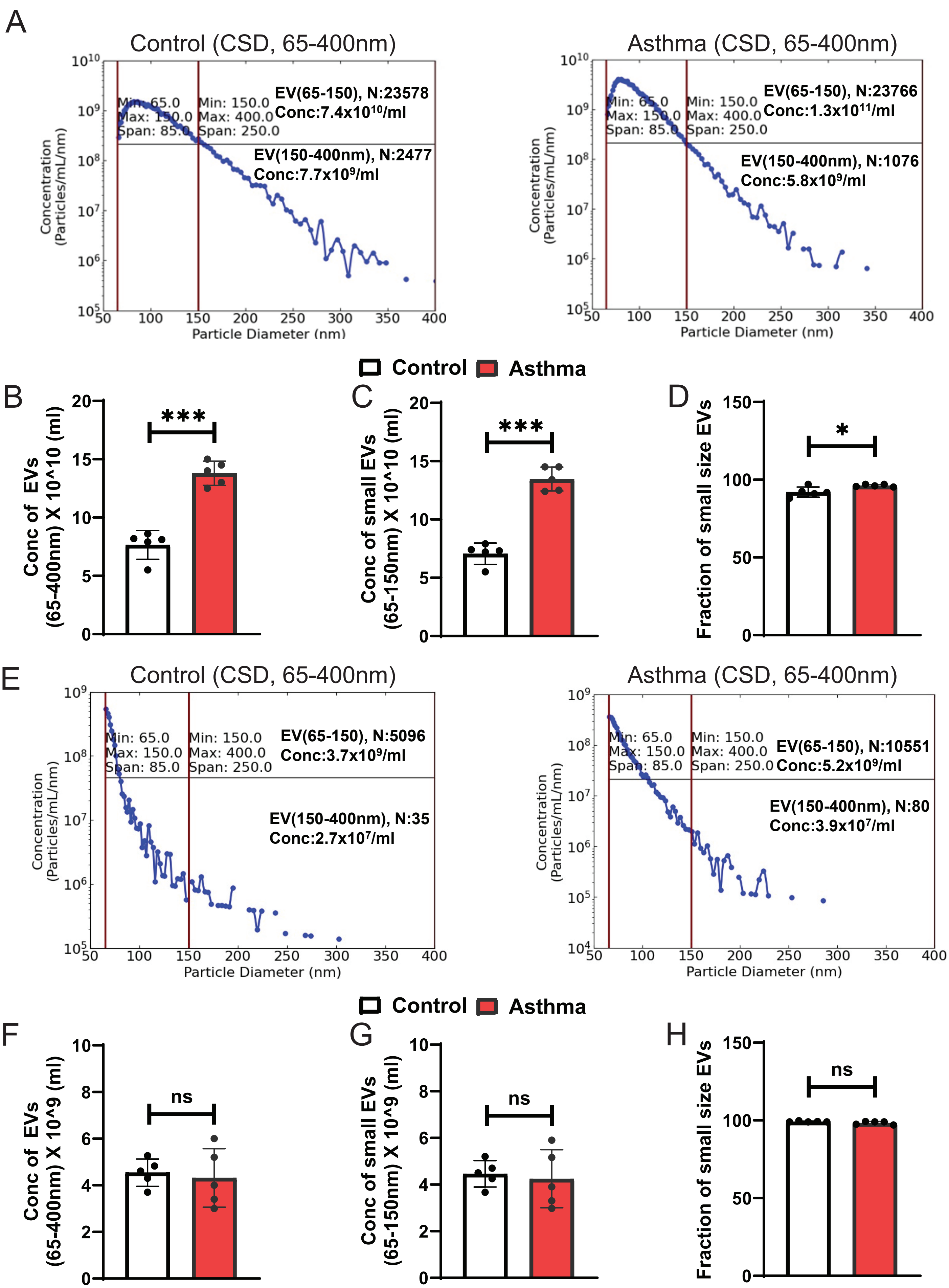

A
Control (CSD, 65-400nm)
EV(65-150), N:23578
Conc:7.4x10¹⁰/ml
EV(150-400nm), N:2477
Conc:7.7x10⁹/ml
Min: 65.0
Max: 150.0
Span: 85.0
Min: 150.0
Max: 400.0
Span: 250.0
Concentration (Particles/mL/nm)
Particle Diameter (nm)
Asthma (CSD, 65-400nm)
EV(65-150), N:23766
Conc:1.3x10¹¹/ml
EV(150-400nm), N:1076
Conc:5.8x10⁹/ml
Control
Asthma
B
Conc of EVs (65-400nm) X 10^10 (ml)
***
C
Conc of small EVs (65-150nm) X 10^10 (ml)
***
D
Fraction of small size EVs
*
E
Control (CSD, 65-400nm)
EV(65-150), N:5096
Conc:3.7x10⁹/ml
EV(150-400nm), N:35
Conc:2.7x10⁷/ml
Asthma (CSD, 65-400nm)
EV(65-150), N:10551
Conc:5.2x10⁹/ml
EV(150-400nm), N:80
Conc:3.9x10⁷/ml
F
Conc of EVs (65-400nm) X 10^9 (ml)
ns
G
Conc of small EVs (65-150nm) X 10^9 (ml)
ns
H
Fraction of small size EVs
ns

**Supplementary Figure 15** – Murine BALF EVs and MDRC-derived EVs are small size EVs. (A) Representative acquired CSD images of quantitation by Spectradyne's nCS1 nanoparticle analyzer of BALF sEVs isolated from MitoQC mice sensitized by OVA and challenged with PBS (Control) and from MitoQC mice sensitized and challenged with OVA (Asthma), Red Box highlights the sEV gate that is quantitated in B-D. (B) Concentration of all BALF EVs comparing Control and Asthma groups. (C) Concentration of BALF sEVs comparing Control and Asthma groups. (D) Percent sEVs of total BALF EVs comparing Control and Asthma. (E) Representative acquired CSD images of quantitation by Spectradyne's nCS1 nanoparticle analyzer of MDRC sEVs isolated from MitoQC mice sensitized by OVA and challenged with PBS (Control) and from MitoQC mice sensitized and challenged with OVA (Asthma), Red Box highlights the sEV gate that is quantitated in F-H. (F) Concentration of all MDRC EVs comparing Control and Asthma groups. (G) Concentration of MDRC sEVs comparing Control and Asthma groups. (D) Percent sEVs of total MDRC EVs comparing Control and Asthma. EVs were isolated from n=6 mice/group from BAL and MDRCs and pooled to obtain EVs for each data point. Data represents 5 replicate experiments. Mann Whitney T test for comparison between Controls and Asthma groups *p<0.05, ****p<0.001.

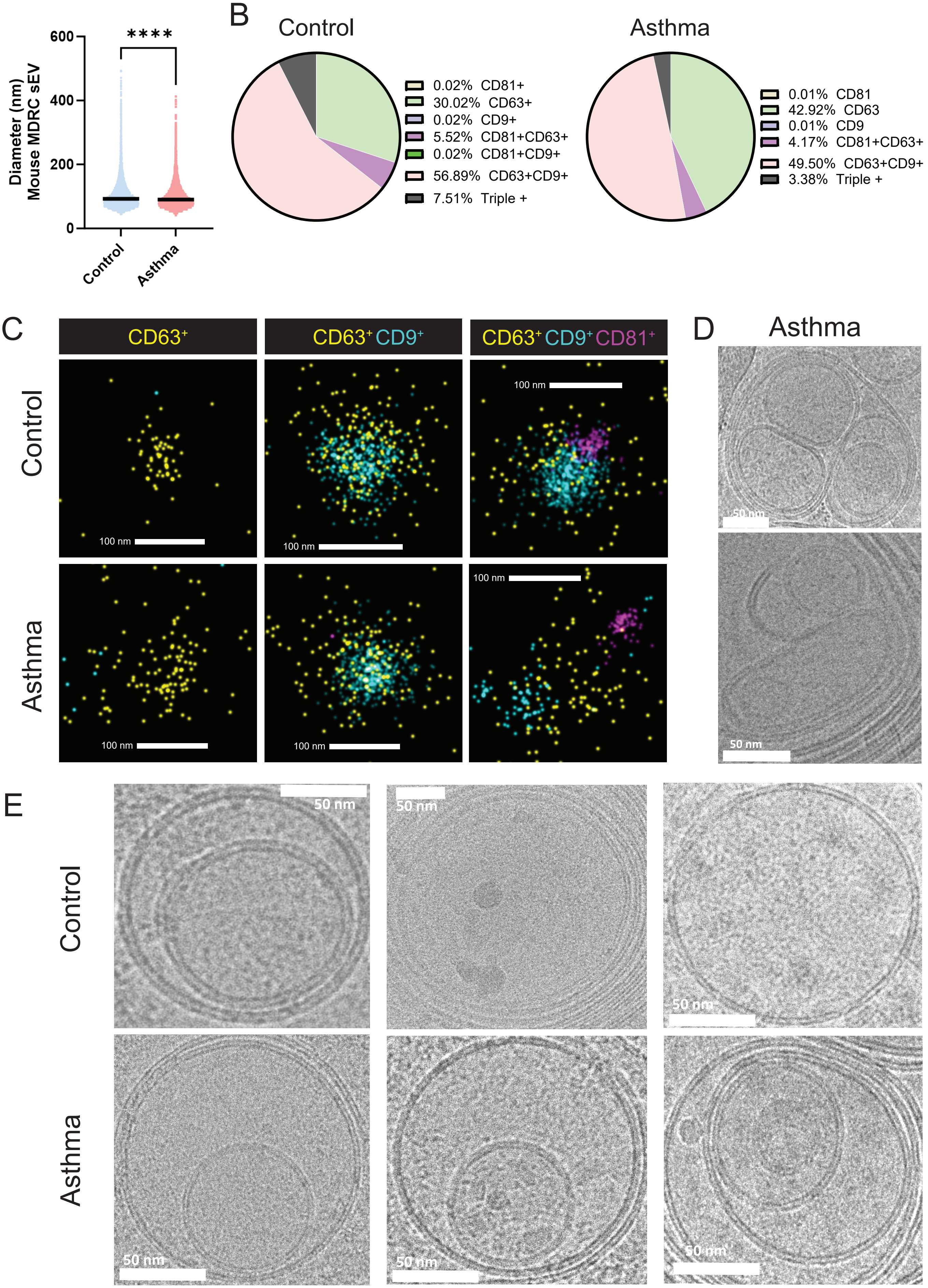

Diameter (nm) Mouse MDRC sEV
600
400
200
0
****
Control
Asthma
B
Control
0.02% CD81+
30.02% CD63+
0.02% CD9+
5.52% CD81+CD63+
0.02% CD81+CD9+
56.89% CD63+CD9+
7.51% Triple +
Asthma
0.01% CD81
42.92% CD63
0.01% CD9
4.17% CD81+CD63+
49.50% CD63+CD9+
3.38% Triple +
C
CD63+
CD63+ CD9+
CD63+ CD9+ CD81+
Control
Asthma
100 nm
D
Asthma
50 nm
E
Control
Asthma
50 nm

**Supplementary Figure 16** – Murine MDRC EVs are small size EVs and they express tetraspanins. Purified MDRC EVs from MitoQC mice were captured and stained using the Oxford Nanoimaging (ONi) EV Profiler Kit 2. Imaging was completed using the AutoEV function on the ONi Nanoimager and utilized direct stochastic optical reconstruction microscopy for high resolution images. All analyses was completed in the CODI software. (A) Quantitation of diameter of pooled MDRC EVs isolated from MitoQC mice sensitized by OVA and challenged with PBS (Control) and from MitoQC mice sensitized and challenged with OVA (Asthma). (B) Pie charts showing % of CD9, CD81 and CD63 expressing MDRC sEVs from samples in (A) determined by Oni image analyses of 25000 events collected n=3 replicates. (C) Representative high resolution images showing sEVs expressing CD63, CD9 or CD81 isolated from MDRCs from n=5 mice/group of Controls and Asthma groups as above. Scale bar= 100 nm. (D) Representative Cryo-Electron microscopy images showing Multi-vesicular inclusions in MDRC sEVs from Asthma group, Scale bar= 50nm. (E) Representative Cryo-Electron microscopy images showing lipid bilayered sEVs with electron dense inclusions in purified MDRC sEVs from Control and Asthma groups above, Scale bar= 50nm, Kolmogorov Smirnov test for comparison between Controls and Asthma groups, ****p<0.001.

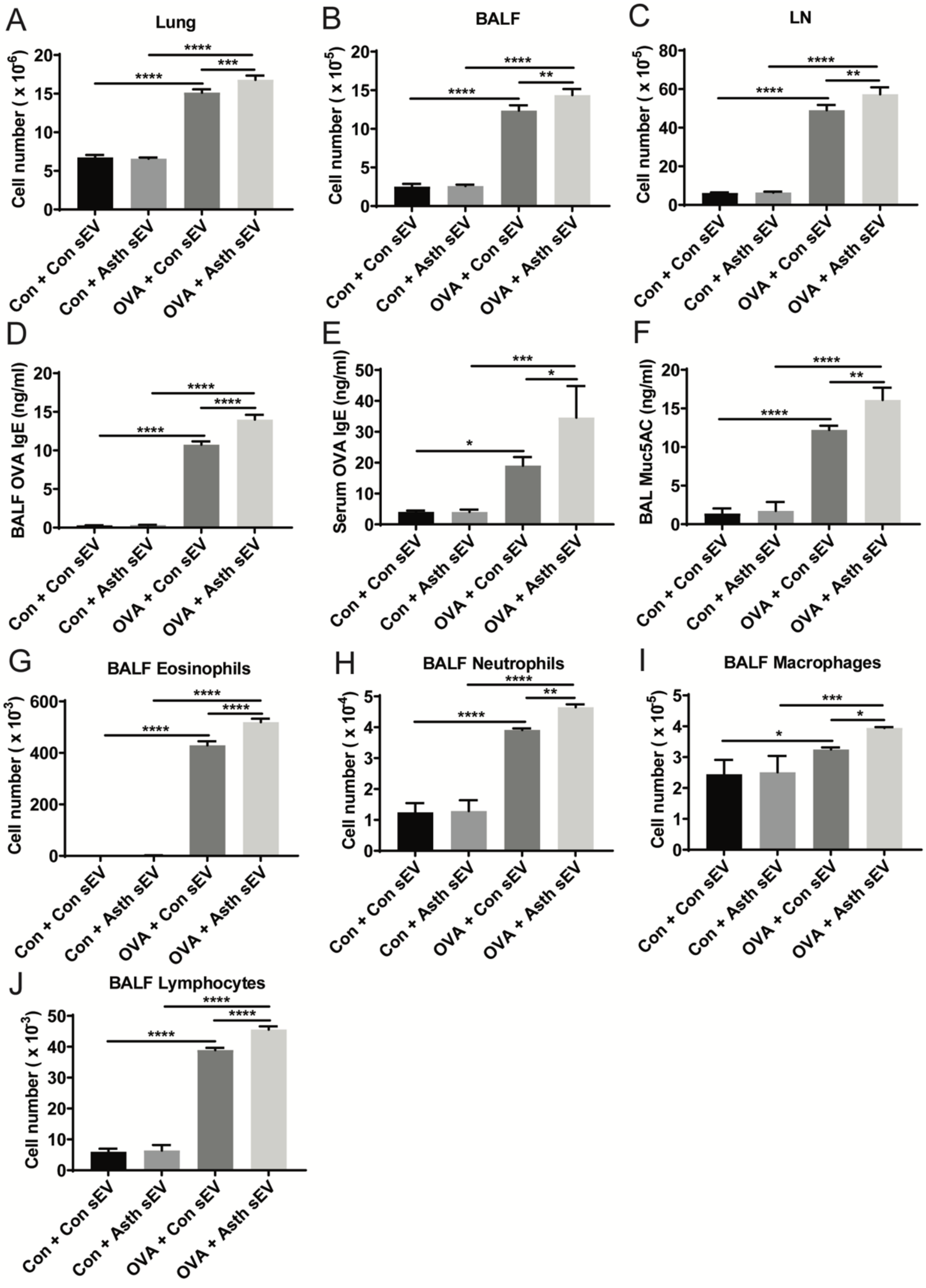


**Supplemental Figure 17 -** Intranasal (i.n.) transfer of asthmatic lung MDRC-derived sEVs exacerbates airway inflammation. Mice were sensitized by intraperitoneal injection on d0 and d7 with 50 µg of alum-adsorbed OVA. On d14, d15 & d16 mice were challenged once i.n. with 15 µg OVA in 30 µl PBS or PBS alone. On d16, 4 hrs after i.n. challenge with OVA or PBS, i.n. delivery of lung proinflammatory MDRC-derived sEVs ($1 \times 10^8$ particles/mouse in 30 µl PBS) from control or OVA challenged donor Mito-QC mice was carried out. (A). Total lung cells from sEV recipient controls and OVA challenged mice at two days after transfer of sEVs. (B). Total BALF cells from sEV-recipient controls and OVA challenged mice at two days after sEV transfer. (C) Total cells of draining LN from sEV-recipient controls and OVA challenged mice at two days after sEV transfer. (D). OVA-IgE levels in BALF harvested at two days after sEV delivery detected by ELISA. (E). OVA-IgE levels in sera at two days after sEV delivery detected by ELISA. (F). Muc5AC levels in BALF by ELISA at two days after-sEV delivery. BALF cells were collected and stained by Diff-Quik. Differential analysis was performed using standard morphological criteria on cytospin slides. 300 cells were examined in each cytospin slide. Numbers of eosinophils (G), neutrophils (H), macrophages (I), and lymphocytes (J) were calculated based on the percentage of each cell population. Statistical significance was evaluated using one-way ANOVA with Tukey's multiple comparison testing. * $P < 0.05$, ** $P < 0.01$, *** $P < 0.005$, **** $P < 0.001$.

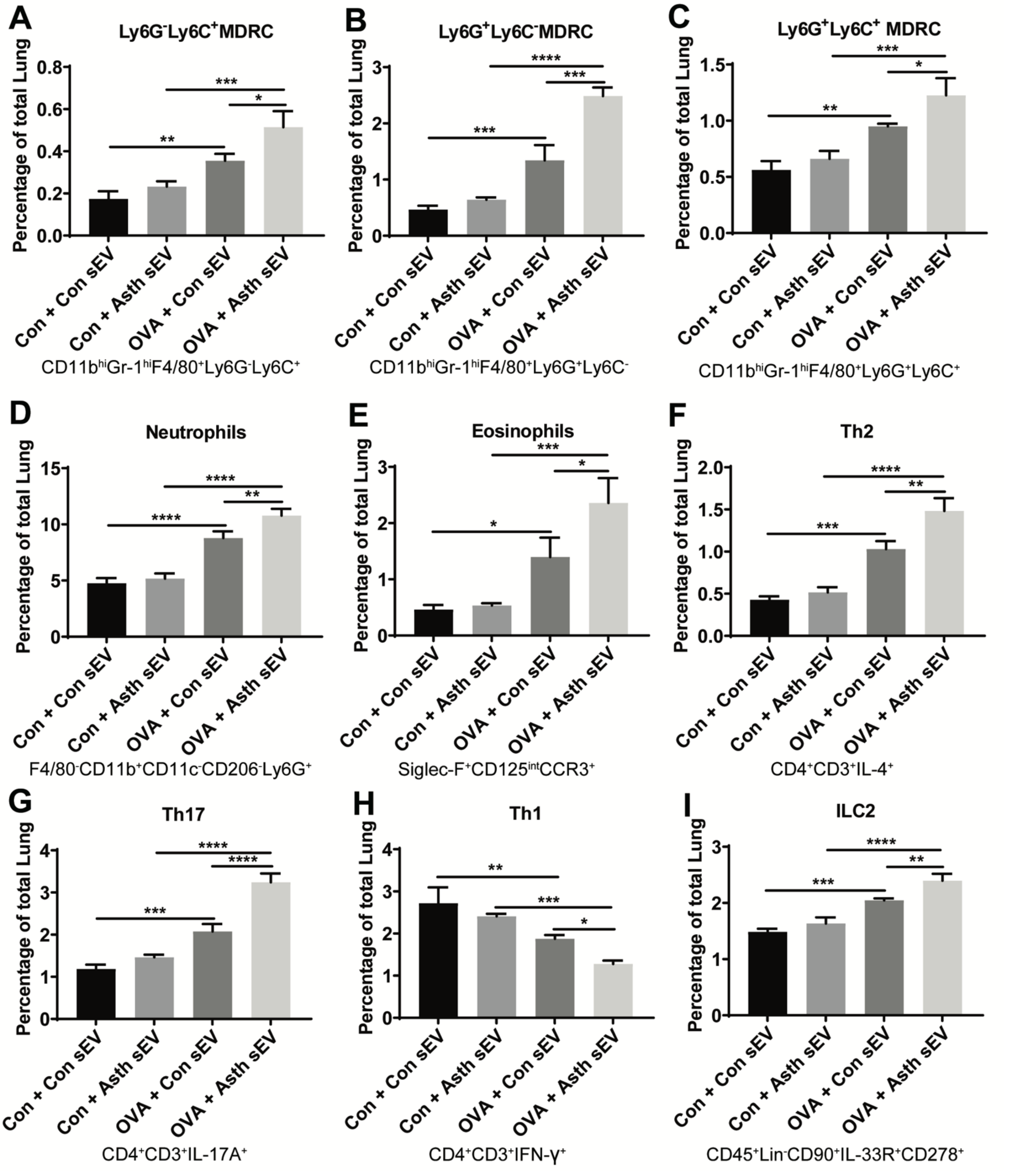


**Supplemental Figure 18** - Intranasal transfer of pro-inflammatory lung MDRC-derived sEVs from sensitized and challenged donor Mito-QC mice with asthma exacerbates Th2 and Th17 responses and allergic airway inflammation in sensitized and challenged recipients. Mice were sensitized and challenged as in Supplemental Figure 8. On d16, i.n. delivery of lung MDRC-derived sEVs (1 x 108 particles/mouse in 30 μl PBS) from control or OVA challenged Mito-QC mice were carried out as before. Lung infiltration of immune cells in lung tissue was determined by FACS analyses. Cell numbers of Ly6G$^-$Ly6C$^+$ MDRCs (A), Ly6G$^+$Ly6C$^-$ MDRCs (B), Ly6G$^+$Ly6C$^-$ MDRCs (C), Neutrophils (D), Eosinophils (E), Th2 cells (F), Th17 cells (G), Th1 cells (H), and ILC2 cells (I) in the lung tissue were determined by flow cytometry. Statistical significance was evaluated using one-way ANOVA with Tukey' s multiple comparison testing. * $P < 0.05$, ** $P < 0.01$, *** $P < 0.005$, **** $P < 0.001$.

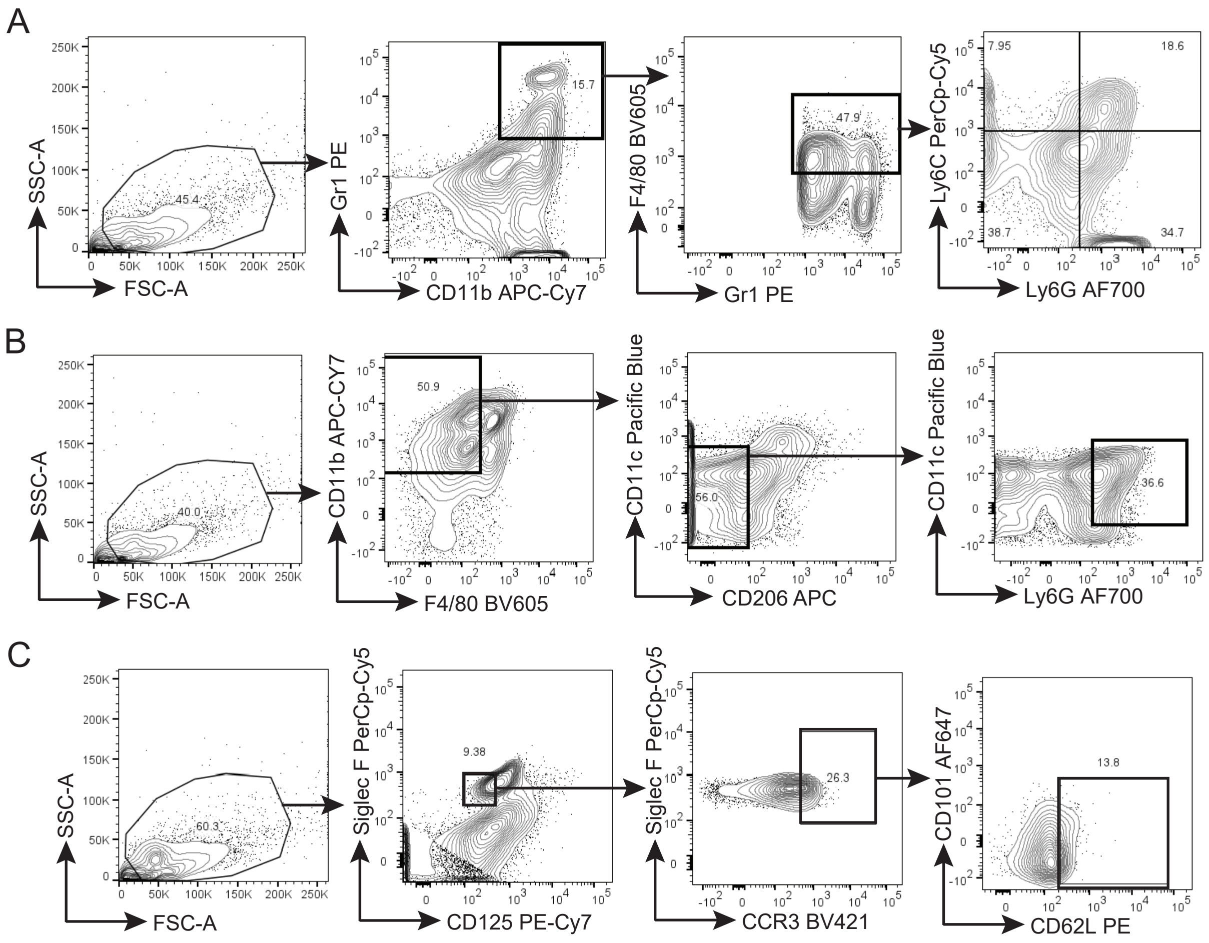


**Supplementary Figure 19** – Gating strategies for myeloid cell populations in the lung tissues of sensitized and challenged mice that are recipients of MDRC-derived sEVs from control or OVA challenged-Mito-QC mice. Mice were sensitized by intraperitoneal injection on d0 and d7 with 50 µg of alum adsorbed OVA. On d14, d15 & d16 mice were challenged once i.n. with 15 µg OVA in 30 µl PBS or PBS alone. On d16, i.n. delivery of lung MDRC-derived sEVs (1 x 108 particles/mouse in 30 µl PBS) from control or OVA challenged Mito-QC mice were carried out as before. Lung infiltration of immune cells in lung tissue was determined by FACS analyses. (A) Gating strategy for MDRC-subpopulations in the lung tissue. (B) Gating strategy for neutrophils in the lung tissue. (C) Gating strategy for eosinophils in the lung tissue.

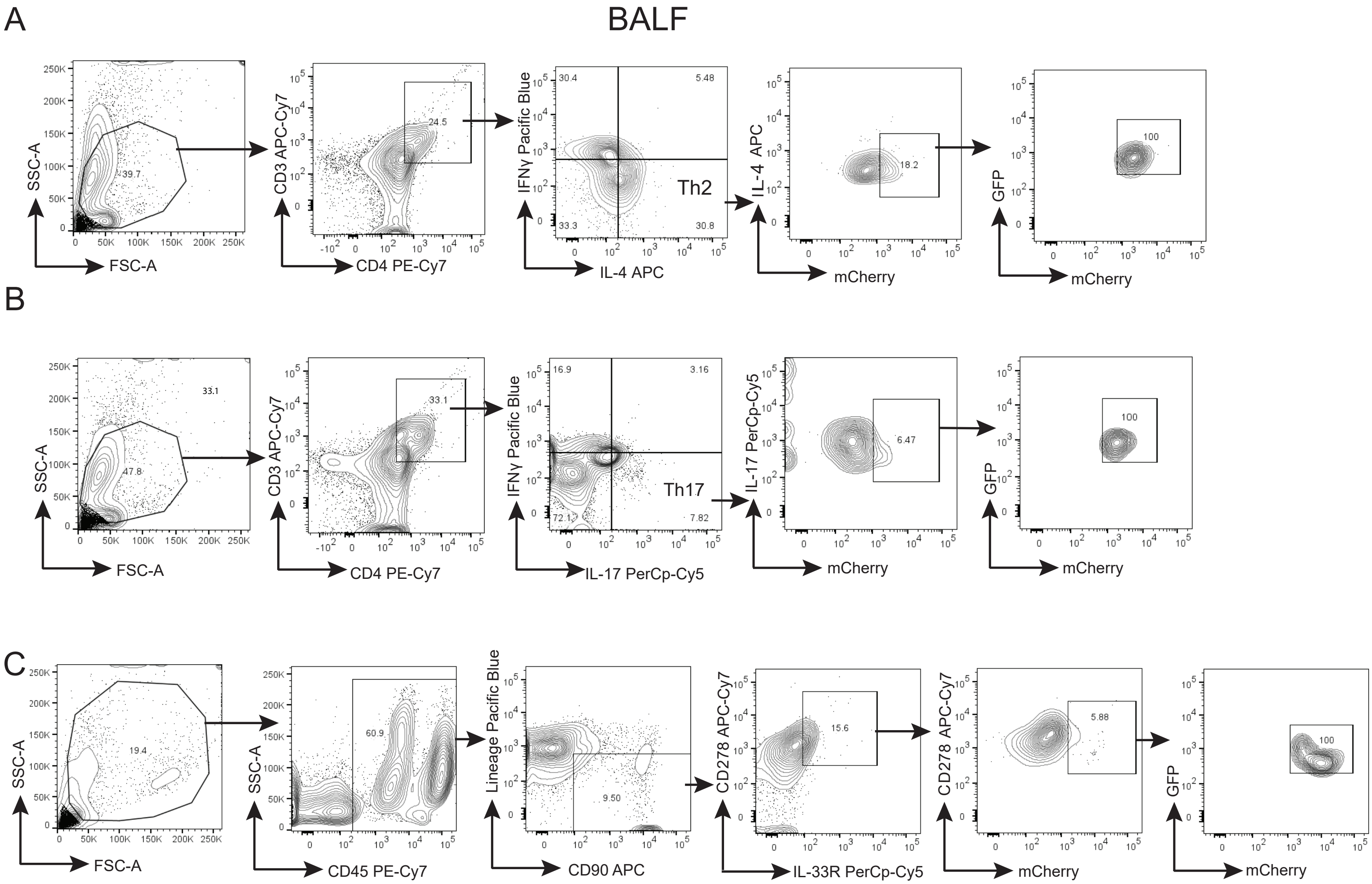


**Supplemental Figure 20** - Gating strategies for mCherry$^+$GFP$^+$ Th2 and Th17 cells in the BALF of sensitized and challenged recipients. Mice were sensitized by intraperitoneal injection on d0 and d7 with 50 µg of alum-adsorbed OVA. On d14, d15 & d16 mice were challenged once i.n. with 15 µg OVA in 30 µl PBS or PBS alone. On d16, i.n. delivery of lung MDRC-derived sEVs (1 x 108 particles/mouse in 30 µl PBS) from control or OVA challenged Mito-QC mice were carried out as before. Infiltration of immune cells in BALF was determined by FACS analyses. (A) Gating strategy for Th2 cells showing mCherry$^+$ and mCherry$^+$GFP$^+$ cells in the BALF. (B) Gating strategy for Th17 cells showing mCherry$^+$ and mCherry$^+$GFP$^+$ cells in BALF. (C) Gating strategy for ILC2 cells showing mCherry$^+$GFP$^+$ cells in the BALF.

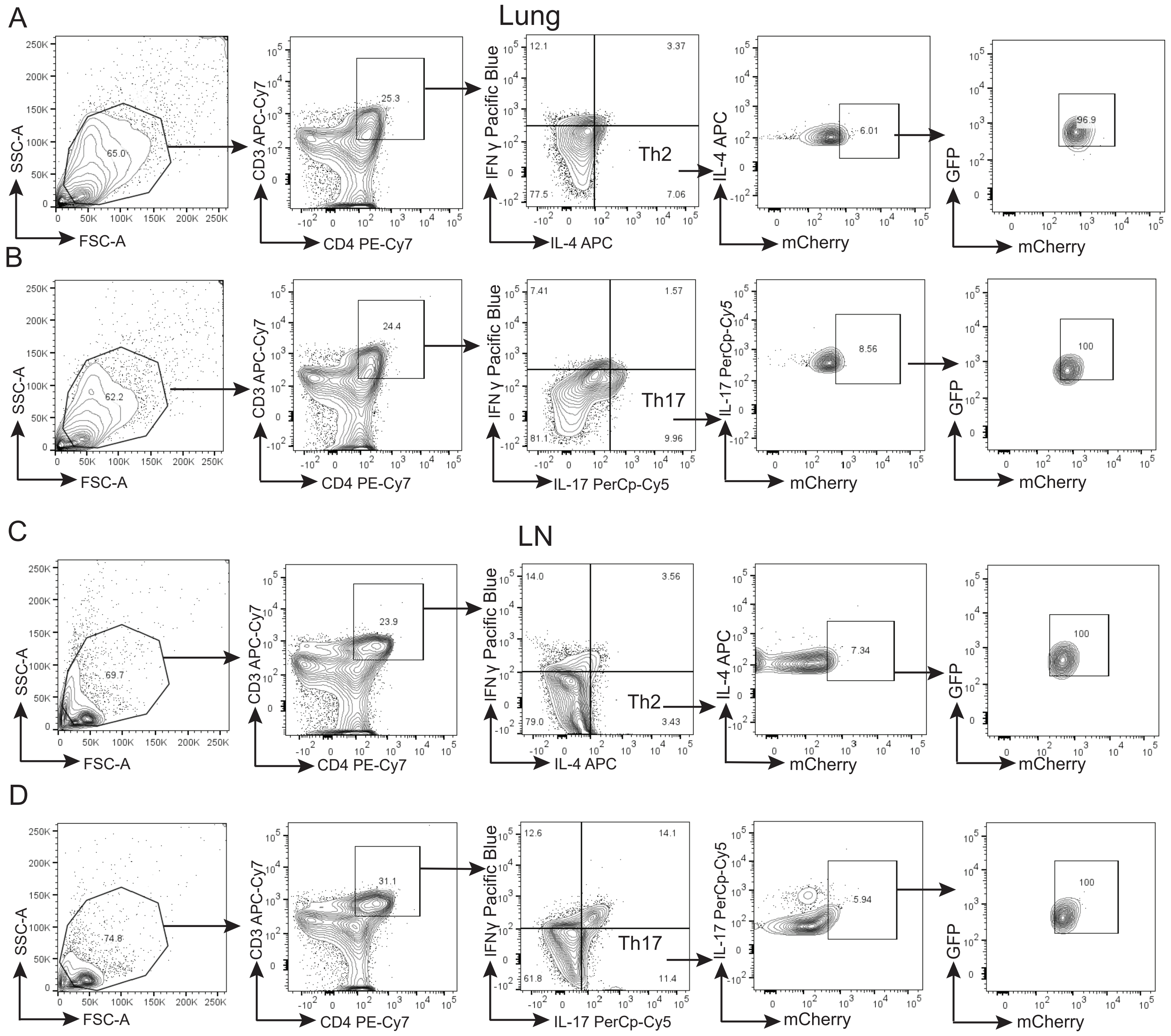


**Supplemental Figure 21** - Gating strategies for mCherry$^+$GFP$^+$ Th2 and Th17 cells in the lung tissue and draining LN of sensitized and challenged recipients. Mice were sensitized by intraperitoneal injection on d0 and d7 with 50 µg of alum-adsorbed OVA. On d14, d15 & d16 mice were challenged once i.n. with 15 µg OVA in 30 µl PBS or PBS alone. On d16, i.n. delivery of lung MDRC-derived sEVs (1 x 108 particles/mouse in 30 µl PBS) from control or OVA challenged Mito-QC mice were carried out as before. Lung infiltration of immune cells in lung tissue was determined by FACS analyses. (A) Gating strategy for Th2 cells showing mCherry$^+$ and mCherry$^+$GFP$^+$ cells in the lung. (B) Gating strategy for Th17 cells showing mCherry$^+$ and mCherry$^+$GFP$^+$ cells in the lung. Immune cell populations in draining LN tissue were detected by FACS analyses. (C) Gating strategy for Th2 cells showing mCherry$^+$GFP$^+$ population in the LN. (D) Gating strategy for Th17 cells showing mCherry$^+$ and mCherry$^+$GFP$^+$ population in the LN.

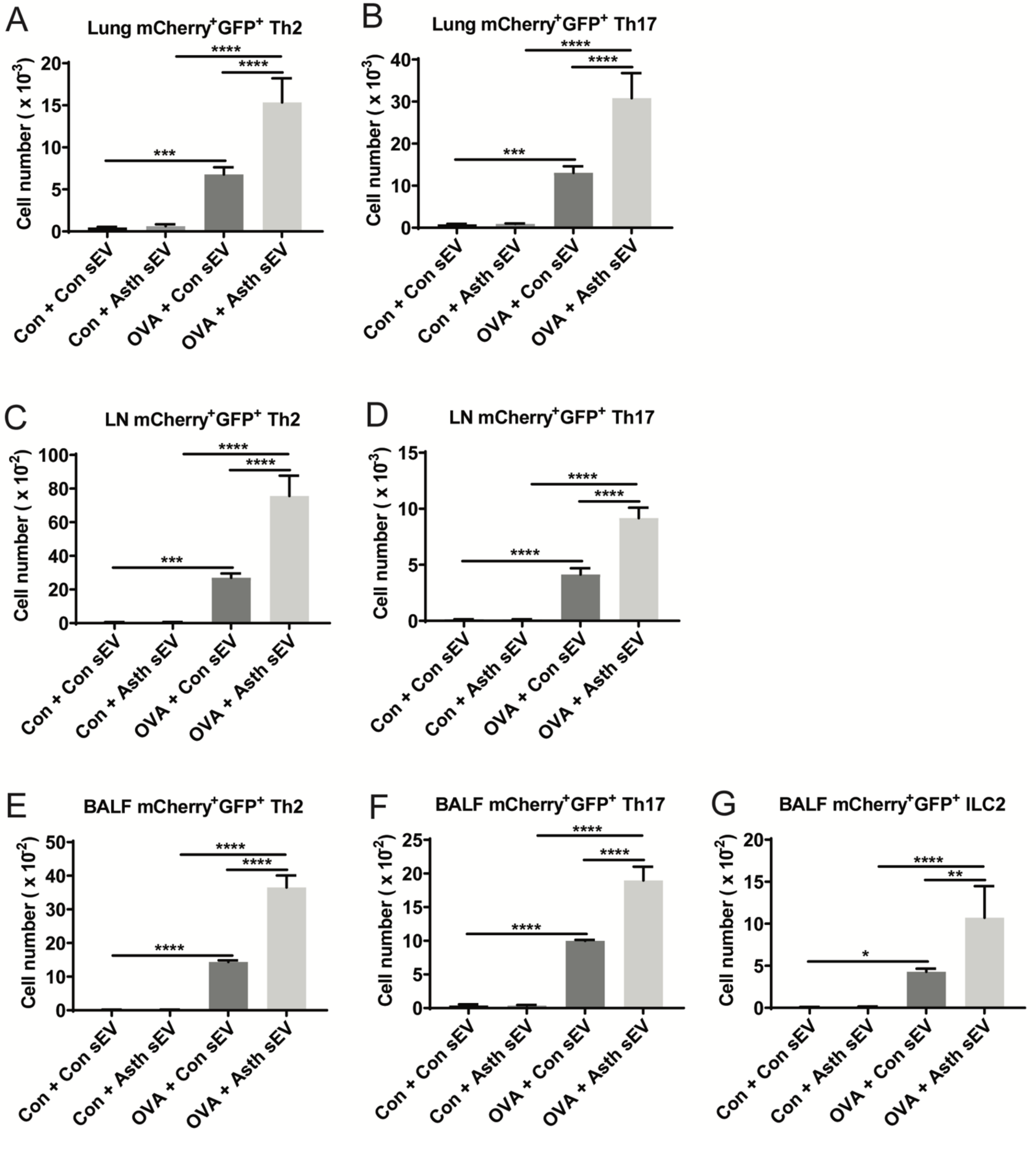


**Supplemental Figure 22 -** Intranasal transfer of pro-inflammatory lung MDRC-derived sEVs from sensitized and challenged donor Mito-QC mice with asthma enhances mCherry$^+$GFP$^+$ Th2 and Th17 cell infiltrations in the lung tissue, draining LNs and BALF in sensitized and challenged recipients. Mice were sensitized by intraperitoneal injection on d0 and d7 with 50 µg of alum-adsorbed OVA. On d14, d15 & d16 mice were challenged once i.n. with 15 µg OVA in 30 µl PBS or PBS alone. On d16, i.n. delivery of lung MDRC-derived sEVs (1 x $10^8$ particles/mouse in 30 µl PBS) from control or OVA challenged Mito-QC mice were carried out as before. Infiltration of immune cells in the lung, BALF and draining LNs was determined by FACS analyses. Cell numbers of mCherry$^+$GFP$^+$ Th2 (A) and Th17 (B) in the lung tissue were determined by flow cytometry. Cell numbers of mCherry$^+$GFP$^+$ Th2 (C) and Th17 (D) in draining LNs were determined by flow cytometry. Cell numbers of mCherry$^+$GFP$^+$ Th2 (E), Th17 (F) and ILC2 (G) in BALF were determined by flow cytometry. Statistical significance was evaluated using one-way ANOVA with Tukey's multiple comparison testing. * $P < 0.05$, ** $P < 0.01$, *** $P < 0.005$, **** $P < 0.001$.
